# Bovine viral diarrhoea virus loses quasispecies diversity rapidly in culture

**DOI:** 10.1101/2020.01.09.900332

**Authors:** George C. Russell, Ruth N. Zadoks, Kim Willoughby, Claudia Bachofen

**Affiliations:** Moredun Research Institute, Pentlands Science Park, Midlothian, EH26 0PZ, UK

**Keywords:** pestivirus, next-generation-sequencing, genome variation, evolution, quasispecies

## Abstract

Bovine viral diarrhoea (BVD) is an important disease of cattle with significant impacts on animal health and welfare. The wide host range of the causative pestiviruses may lead to formation of virus reservoirs in other ruminant or wildlife species, presenting a concern for the long-term success of BVD eradication campaigns. It is likely that the quasispecies nature of these RNA viruses contributes to their interspecies transmission by providing genetic plasticity. Understanding the spectrum of sequence variants present in BVD persistently infected (PI) animals is therefore essential for studies of virus transmission. To analyse quasispecies diversity without amplification bias, we extracted viral RNA from serum of a PI cow, and from cell culture fluid after three passages of the same virus in culture, to produce cDNA without amplification. Sequencing of this material using Illumina 250bp paired-read technology produced full-length virus consensus sequences from both sources and demonstrated the quasispecies diversity of this Pestivirus A type 1a field strain within serum and after culture. We report the distribution and diversity of over 800 single nucleotide polymorphisms and provide evidence for a loss of diversity after only three passages in cell culture, implying that cultured viruses cannot be used to understand quasispecies diversity and may not provide reliable molecular markers for source tracing or transmission studies.

Additionally, both serum and cultured viruses could be sequenced as a set of 25 overlapping PCR amplicons that demonstrated the same consensus sequences and the presence of many of the same quasispecies variants. The observation that aspects of the quasispecies structure revealed by massively parallel sequencing are also detected after PCR and Sanger sequencing suggests that this approach may be useful for small or difficult to analyse samples.

**Impact statement:** Bovine viral diarrhoea viruses are globally important cattle pathogens, which impact performance due to acute infection and BVD-induced immunosuppression. Eradication of BVD in cattle is widely pursued but is hampered by the production of persistently infected (PI) calves – the offspring of cows infected in early pregnancy – which shed virus constantly and drive BVD spread. Genetic variation in BVD viruses is an important feature of their biology, allowing them to adapt to changing conditions and to infect different hosts. Inaccurate virus replication produces a population of viruses with slightly different sequences, a quasispecies, some of which may grow better in other hosts or in culture. Analysing virus sequence variation may help us understand how the virus evolves within and between its hosts. In this paper we show that a BVD virus strain loses quasispecies diversity quickly when cultured and that these changes can be detected even in small diagnostic samples, implying that cultured viruses do not perfectly represent the field strains they were isolated from and therefore may not provide reliable molecular markers for source tracing or transmission studies.

**Data Summary:** Pestivirus A genome sequences used in this article are as follows:

| strain | accession number |
| --- | --- |
| 180 | HQ174292 |
| 8844 | HQ174293 |
| GS5 | KJ541471 |
| NADL | M31182 |
| SD1 | M96751 |
| SingerArg | DQ088995 |
| MRI103 | LR699799 |
| 526 | HQ174294 |
| 8824 | HQ174295 |
| CC13B | KF772785 |
| CP7-5A | AF220247 |
| IBSP4ncp | KJ620017 |
| KE9 | EF101530 |
| Osloss | M96687 |
| ILLNC | U86600 |
| 10JJ-SKR | KC757383 |
| Carlito | KP313732 |

Sequence data associated with this manuscript has been submitted to the European Nucleotide Archive (www.ebi.ac.uk/ena/) with accession numbers as follows:

Consensus genome sequences:

MRI103 serum NGS: LR699799

MRI103 culture NGS: LR699800

MRI103 serum Sanger: LR699801

MRI103 culture P3 Sanger: LR699802

MRI103 culture P5 Sanger: LR699803

NGS raw data

Serum dataset: ERR3624580

Culture dataset: ERR3624581

## Introduction

Bovine viral diarrhoea (BVD), caused by viruses of the genus *Pestivirus*, family *Flaviviridae*, is an important cattle pathogen causing immunosuppression, ill-thrift and a lethal form of diarrhoea called mucosal disease (1). The BVD viruses have a positive sense single-stranded RNA genome of about 12500 nucleotides that is replicated in an error-prone manner, generating a wide range of related virus sequences within an infected individual – a “quasispecies cloud” (2, 3). BVD viruses exist as two major types, of which in Britain type 1 prevails and type 2 is only rarely identified (4, 5). Recently, however, pestivirus nomenclature has been revised so that BVDV type 1 (now Pestivirus A) and BVDV type 2 (Pestivirus B) are considered distinct virus species (6). Pestivirus A genotypes are defined by sequence similarity, particularly in the 5’ untranslated region (5’-UTR) with at least 23 genotypes of Pestivirus A being recognised (4, 7-10).

Due to the economic impact of this infection, several European nations, including Scotland, have ongoing control and/or eradication programmes. During such programmes, which mainly target bovine species, re-infection of previously BVD-free herds may occur, depending on the degree of biosecurity that can be maintained and the number of persistently infected (PI) animals remaining in the national cattle population. However, it is also possible that reinfection may occur from infected wildlife or non-bovine livestock species, particularly deer and small ruminants (11-14). Thus, the broad host range of BVD viruses (all even-toed ungulate species may be infected) may hinder the long-term success of BVD eradication. Indeed, with prevalence in pigs reported to be increasing internationally in recent years (15-17), the risk of spread among a wider range of livestock species should also be considered. In this situation, molecular epidemiology is a useful approach to finding sources of infection and routes of virus transmission (18-21).

The quasispecies nature of this RNA virus may contribute to interspecies transmission by providing genetic plasticity. It was shown previously that transmission of BVD from cattle to goats and between cattle and sheep led to nucleotide changes in the coding region of E2, the major receptor binding protein and host species determinant, and changes to the binding of monoclonal antibodies (22-24). Furthermore, in vitro experiments using cell cultures of bovine and ovine origin showed that such adaptive mutations were reversible within just 3 cell culture passages and were therefore most likely due to selection of pre-existing variants (25). Analysis of the spectrum of virus variants present in PI animals may reveal important information on the quasispecies dynamics involved in interspecies transmissions, and may help to reveal unusual reservoirs and transmission routes of BVD during the final stages of eradication campaigns (11,12). Thus, understanding quasispecies dynamics of pestiviruses and how this may give rise to variants or emerging viruses would enhance our understanding of interspecies transmission between wildlife and livestock species. Genome sequencing of BVD viruses has been possible for at least 20 years. Initial strains were sequenced using cDNA cloning and PCR (26-29), whilst more recent work has used reverse transcription (RT-) PCR and direct sequencing or next generation sequencing (NGS) (30-41). However, many pestivirus genome sequences were derived from isolates that were cultured before analysis and therefore do not necessarily represent the sequence diversity of the virus strain infecting its host. Few studies have taken advantage of NGS technology to look directly at quasispecies diversity in culture or *in vivo* (42) and none have looked at how the quasispecies diversity of BVD viruses changes during the initial passages of virus culture. In this paper we describe the direct sequencing of a Pestivirus A strain from serum and of the same strain after three passages in culture. In each case care was taken to remove potential sources of bias in the dataset, particularly avoiding amplification of the source material before construction of the sequencing libraries.

## Methods

### Viruses and culture

Laboratory strain NADL was obtained from the Moredun Virus Surveillance Unit and was grown on bovine turbinate cells in Iscove’s modified Dulbecco’s medium (IMDM), by inoculation at a multiplicity of infection (MOI) of 0.1. Viral cultures were expanded by serial subculture through three passages to produce sufficient material for virus concentration and RNA extraction.

A BVD PI cow (MRI103), which was held at the Scottish Centre for Production Animal Health & Food Safety, University of Glasgow for teaching purposes, was humanely euthanized by intravenous injection to prevent the spread of disease after confirmation of PI status. As part of the necropsy, samples of whole blood were collected and serum was stored at −20°C in aliquots of up to 50 ml. Serum samples were used as the source of virus for these studies after about two years of storage without thawing. The virus strain was characterized by RT-PCR and sequencing of 5’UTR and Npro gene fragments, which confirmed that this isolate was a BVDV1a genotype (43). This strain had also been used to infect rabbits, demonstrating that it could propagate *in vivo* and induce virus neutralizing antibodies (44). In order to define the effects of low-passage culture on the quasispecies diversity of BVD viruses, this isolate was passaged up to five times on bovine turbinate cells at MOI of 0.1.

### RNA extraction and purification

Prior to RNA extraction, virus particles from serum and culture medium samples were concentrated. The uncultured MRI103 serum sample was concentrated by ultracentrifugation of 50 ml of serum at 120 000 ×g for 3h at 4 °C. The pelleted virus was resuspended in 1.3 ml of phosphate buffered saline (PBS).

The medium from cultured virus was first clarified by low speed centrifugation before concentration of virus particles by ultrafiltration using Centricon Plus-70 Centrifugal Filter Units, (Merck, 100 KDa cutoff). Approximately 350 ml of culture fluid was concentrated 100-fold before RNA extraction.

Concentrated virus particles were treated with RNase A before extraction of RNA using Trizol LS (Ambion, Life Technologies). The aqueous fraction of the Trizol lysate was precipitated with ethanol and was resuspended in 10 mM Tris pH 8.0 before purification using the QIAamp viral RNA mini kit (QIAGEN, Manchester, UK) in the absence of carrier RNA. Purified RNA was treated with RTS DNase (Cambio, Cambridge, UK) before cDNA synthesis.

### Analysis of viral RNA

The degree of contamination of the viral RNA preparations with residual host or cell-culture material was analysed by real-time PCR and RT-PCR using a triplex assay for BVD (45), β-actin (46) and mitochondrial DNA (Table 1). These assays allowed the yield of viral RNA to be compared with potential host genomic or mitochondrial DNA contamination, respectively. The recovery of the full-length NADL viral genome was analysed by a set of long RT-PCR assays that covered the full length of the virus in either three or five fragments (not shown). The primer sequences for this analysis were not compatible with amplification of BVD strain MRI103, so the likely recovery of full-length RNA for this genome was inferred from NADL RNA purified in parallel.

**Table 1.**
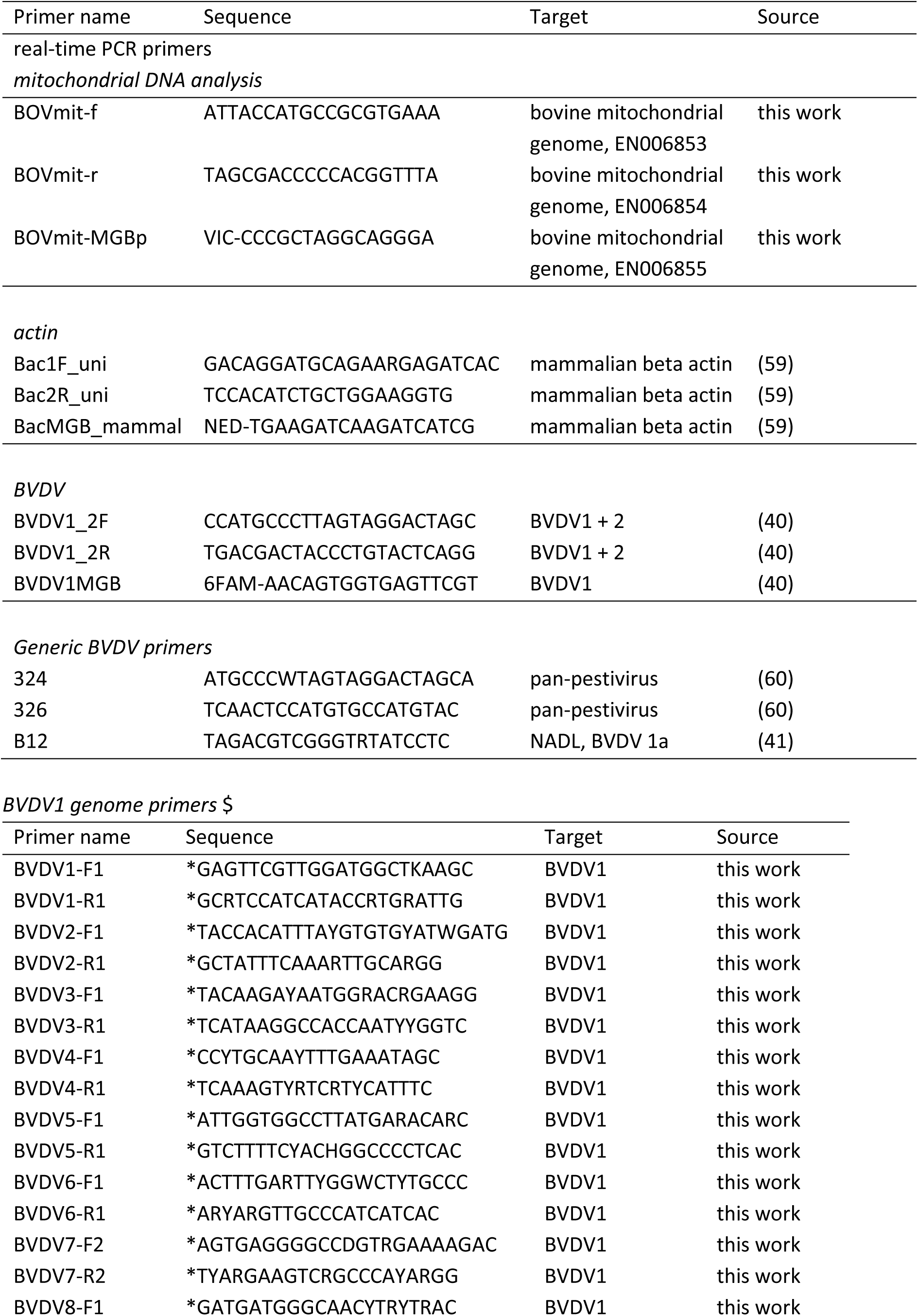
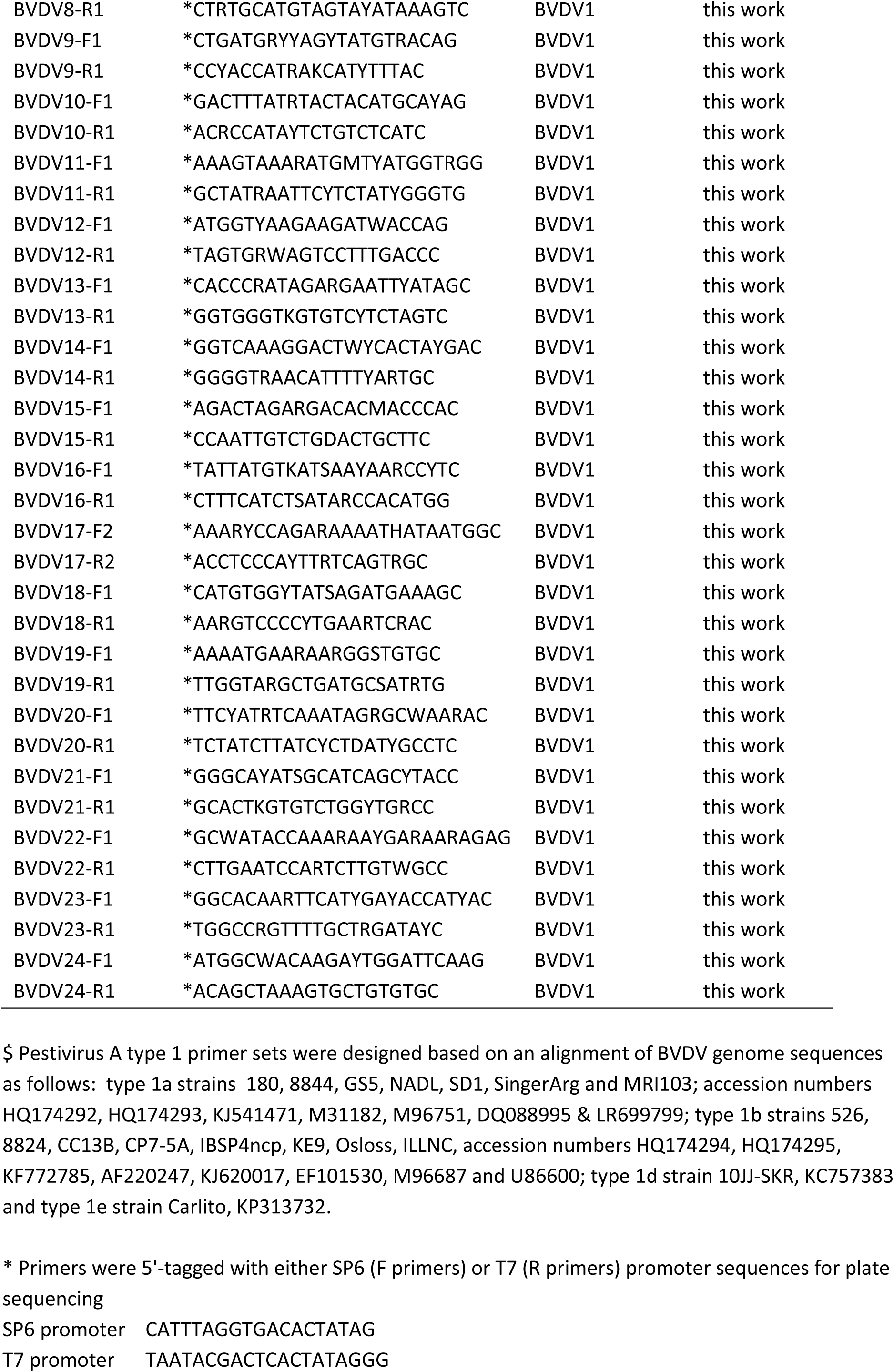
Primers used in this work.

### cDNA synthesis

Initial experiments showed that priming cDNA synthesis with random primers led to a poorer yield of full-length cDNA, with clear reduction in representation of the 3’-end of the genome. In order to improve coverage of full-length genomes, two approaches were used. In the first approach, a Pestivirus A genotype 1a-specific primer (B12; Table 1) was used to prime cDNA synthesis from the 3’-end of the viral genome, whilst in the second approach, viral RNA samples were polyadenylated using polyA polymerase before cDNA synthesis primed by anchored oligo dT primers. In each case, synthesis of double-stranded cDNA was done using the Superscript III cDNA synthesis system (Life Technologies), with RNAseH, *E. coli* DNA polymerase and *E. coli* DNA ligase in the second strand mix.

The presence of double-stranded BVD-specific cDNA after synthesis was assayed qualitatively by end-point PCR before and after restriction endonuclease digestion of the cDNA, using primers that amplified either conserved regions of the virus (all samples) or the entire NADL genome as a set of overlapping amplicons. Restriction enzyme treatment of cDNA clearly reduced the yield of BVD PCR products (not shown).

### Sequencing

Six viral cDNA samples were sequenced by The Genome Analysis Centre (TGAC; now Earlham Institute) as part of their Capacity and Capability Challenge (CCC) programme. The samples were extracted from the two viruses, NADL and MRI103, with MRI103 purified from serum and after three passages in culture (P3), and each cDNA was primed either with oligo-dT or with the BVD-specific 3’-UTR primer B12 (47). The cDNA samples were therefore named: NADL-dT; NADL-B12; MRI103-dT; MRI103-B12; MRI-P3-dT and MRI-P3-B12.

Library generation used the TruSeq CHiP protocol (Illumina) because of the small amounts of cDNA available (< 15 ng DNA per sample) and each library was tagged with barcoding primers to allow multiplex analysis of all six samples in a single MiSeq run with 250 base paired-end reads. Data was returned as FastQ format files.

### Sequence assembly

The individual datasets were mapped against the bovine and pestivirus genomes to get an estimate of read coverage before assembly. An initial mapping of reads from the oligo-dT primed serum derived sequences to the Pestivirus A isolate SD-1 genome (26) was performed at low stringency using the program BWA (48). The preliminary consensus sequence derived from this analysis allowed the reference-led assembly of the MRI103 virus genome datasets.

Assembly of the sequence data for each cDNA sample was done using the DNASTAR Lasergene SeqMan Ngen program, using the preferences for a reference-led assembly of paired-end reads. Each MiSeq sequence assembly was then transferred from SeqMan Ngen to Seqman Pro for final quality control and SNP calling.

### SNP analysis

Polymorphic positions within each consensus were called using the SNP Report feature within Seqman Pro and validated by inspection of the assembly. SNPs represented by a single base call within each assembly were discarded while all other SNP were retained for further analysis of diversity. Areas of low coverage (<100-fold) were not included in the analysis and SNPs within homopolymeric tracts at the end of reads were disregarded. The SNP reports were copied into MS Excel spreadsheets for further analysis. SNP positions were reported based on the numbering of the respective MRI103 serum or NADL (M31182) consensus sequence (29).

Sequence diversity within each assembly was measured as the Shannon Entropy (H) at each polymorphic base position, calculated as the negative sum of the product of each base call frequency and its natural log, as described by Milani and coworkers (49). Diversity for each sequence assembly was plotted as the distribution of single base entropy values or as the mean single nucleotide Shannon entropy within each of the individual coding regions.

Error rates within the sequence data were estimated from the base call quality (Phred) scores in the final edited assemblies. Average read length in the assembled data was 235 and mean read quality score was 36 with range 22-39. The minimum mean read quality score was therefore used to calculate a conservative threshold entropy score below which SNP were excluded as potential read errors. For a read quality value of 22, the estimated error rate was 0.63 %, with a corresponding H value of 0.0384. Thus only polymorphic positions with H >0.0384 were considered to be SNP in this analysis. In addition, overall entropy for each genome was calculated as the sum of all H values above the threshold value.

### Genome analysis by PCR and Sanger sequencing

As an approach to deriving close to full-length Pestivirus A genome sequences from small amounts of virus material without propagation of virus, a panel of 24 primer pairs was designed based on an alignment of 17 genome sequences representing BVD virus genotypes found in the UK (7 type 1a including MRI103, 8 type 1b, 1 type 1d, 1 type 1e; Table 1). Degenerate primers were designed using the program PrimerDesign-M (50), with the following parameters: maximum degeneracy 32-fold, primer length 20-25, overlap mid, Tm difference 5 °C. Primers were synthesised by Eurofins-MWG tagged with the SP6 promoter sequence on all forward primers and the T7 promoter sequence on all reverse primers as sequencing adaptors (Table 1).

In addition, sequences derived from amplification with the pan-pestivirus 5’UTR primers 324 and 326 (51) added approximately 56 nucleotides (nt) at the 5’end of each sequence. RNA from the original material used for next-generation sequencing of the serum and passage 3 cultured MRI103 virus was used as the template for amplification using the One-Step Ahead RT-PCR kit (Qiagen).

To demonstrate genome sequencing from small samples, RNA purified from 100 µl of MRI103 passage 5 (P5) culture medium was amplified as above. This also allowed the analysis of sequence changes introduced by continued culture.

The PCR products were sequenced using SP6 and T7 promoter primers (Eurofins MWG) and trace files were assembled using the DNASTAR Lasergene SeqMan program (DNASTAR Inc, Madison, WI, USA). All potential SNP sites were assessed visually and were considered valid if there was clear evidence of multiple base calls at the candidate site in two or more sequencing reads. Variant bases were recorded as UPPERCASE/lowercase (e.g. G/a) if all relevant reads showed the same majority base call and as International Union of Pure and Applied Chemistry (IUPAC) ambiguity codes (e.g. R for G/A) if the majority base could not be identified from the available reads.

## Results

### Viral RNA purification and cDNA synthesis

MRI103 viral RNA was purified from approximately 50 ml of serum or 330 ml of culture medium and double stranded cDNA was synthesised using either anchored oligo dT primers or a BVD-specific primer (B12, Table 1). A triplex real-time PCR, using primer-probe sets for BVD, mitochondrial DNA and genomic B-actin (Table 1), demonstrated that all of the cDNA samples contained BVD-specific cDNA which was enriched compared to genomic or mitochondrial DNA (data not shown).

Multiple end point PCR assays (with and without prior treatment by restriction enzyme *Mse*I) demonstrated the presence of NADL-specific double-stranded cDNA representing the complete genome (data not shown) and suggested that the MRI103 samples (for which the NADL-specific primers did not work) were also likely to contain full-length viral cDNA.

Quality control of six cDNA samples submitted to the UK Biotechnology and Biological Sciences Research Council (BBSRC) Genome Analysis Centre (TGAC; now Earlham Institute) suggested that the samples contained between 2 and 14 ng cDNA. All of each cDNA sample was used for library construction and after barcoding were combined into a single MiSeq run which yielded a minimum of 1.7 million read-pairs per sample.

### Virus genome Sequencing

Sequencing reads from TGAC were obtained as FastQ files of reads that passed initial quality thresholds. Per-base quality showed that many reads dropped out of the optimal quality range (>28) after about position 200, so all reads were quality trimmed before assembly. Reads that were mainly of poor quality or which contained long homopolymeric stretches were also excluded from further analysis. The reads were analysed to ascertain source (bovine, BVD or unknown) and to estimate the number of reads available for genome assembly (Table 2). This showed that only a small proportion of reads were homologous to BVD viruses in each library (0.2 % to 5.3 % for MRI103) and that the sequence data obtained from the serum-derived virus had the lowest proportion of virus-specific reads. Further analysis was therefore done to establish whether the BVD-specific data was sufficient to produce a whole genome consensus.

**Table 2.**
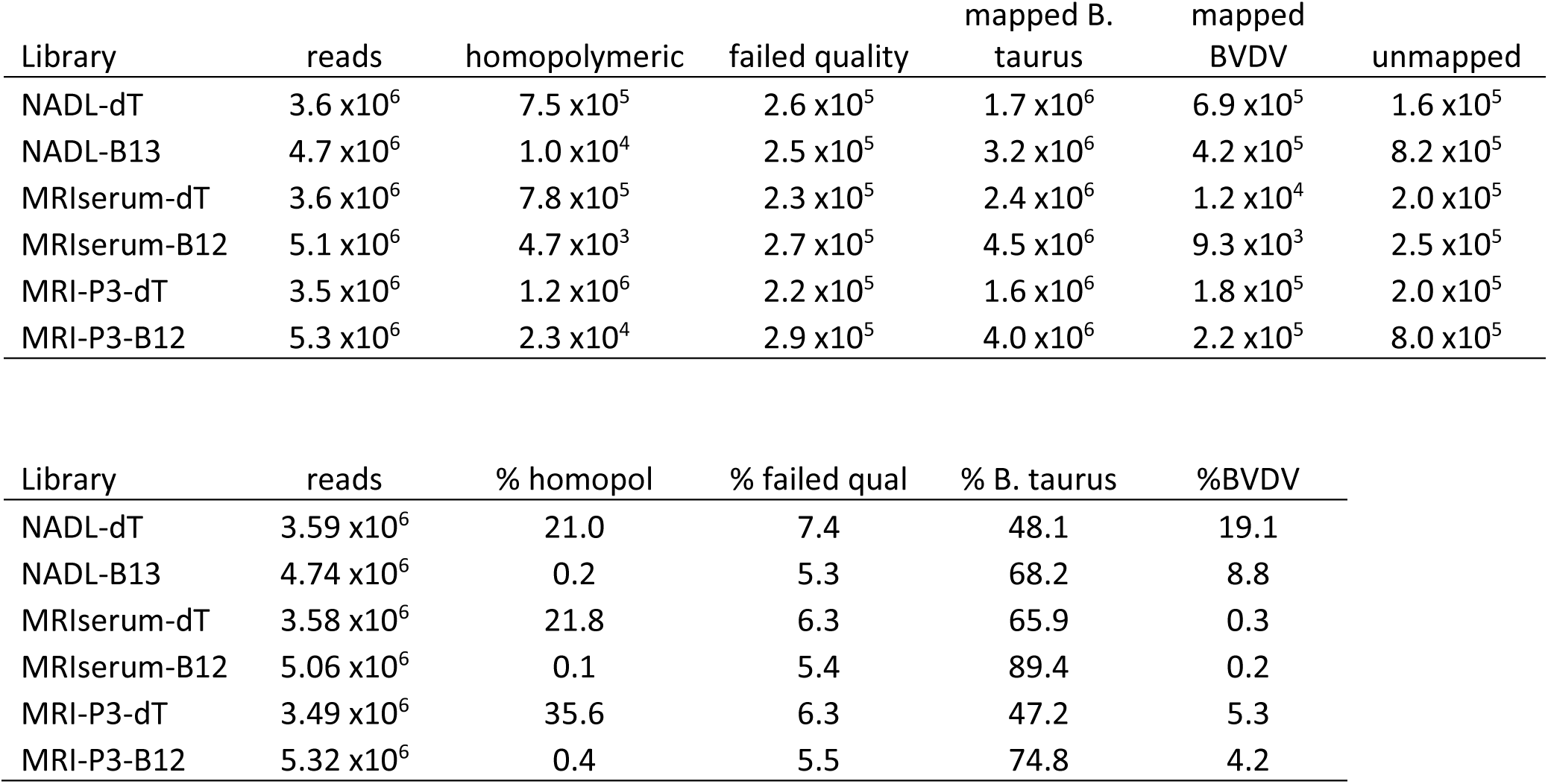
BVD virus NGS measures. Number of reads, quality control data and mapping to *Bos taurus* and BVD virus genomes. Total read numbers and percentages are given for each sequencing library.

As an initial test of assembly and to determine whether a consensus sequence could be produced for reference-led assembly of the MRI103 genome, each set of reads was aligned with the most relevant reference genome sequence using the BWA read alignment package (48). These were NADL (Accession M31182) for the NADL reads and strain SD-1 (M96751) for the MRI103 reads, based on analysis of the known 5’UTR and Npro sequences of MRI103. Alignment to NADL was done at high stringency, while alignment of MRI103 reads to the SD-1 sequence was done at lower stringency to obtain optimal coverage.

These data showed that coverage of NADL was complete with average read-depth >5000 from the oligo-dT primed cDNA, whilst read depth in the data from the B12-primed cDNA was approximately 100-fold lower. When the MRI103 reads were aligned with BVD isolate SD-1, the coverage was essentially complete in the oligo-dT primed cDNA but both coverage and read-depth were lower in the B12-primed cDNA. These data also showed that only the oligo-dT primed cDNA produced enough sequence data to cover either virus genome at read depth >100. A consensus sequence was derived from each alignment, representative of the sequencing data for NADL and MRI103. The NADL consensus was identical to the NADL reference genome and the MRI103 consensus (of 12,292 nucleotides) was 90 % identical to the SD-1 genome sequence.

All six datasets (NADL-dT; NADL-B12; MRI103-dT; MRI103-B12; MRI-P3-dT and MRI-P3-B12) were then used in reference-led assembly of the respective NADL or MRI103 genome sequences using DNASTAR Seqman Ngen. All datasets assembled to their respective reference sequence with high quality and good consistency in the placing of read pairs (supplementary data table 1). However, once again, the B12-primed cDNA datasets provided five- to ten-fold lower read depth than the oligo-dT primed cDNA data. All subsequent analyses were therefore performed using the data obtained from the oligo-dT primed cDNA samples. For each of these datasets, (NADL-dT; MRI103-dT; MRI-P3-dT), a consensus sequence was derived from the assembled reads and this was used to assess the degree of polymorphism within the virus populations assayed. Sequence coverage within the oligo-dT-primed datasets was as follows: MRI103-dT had median read depth of 190 but about 10% of the genome had read depth 50-100; MRI-P3-dT had median read depth of about 1800 and minimum read depth of just over 100. In contrast, the NADL-dT sequence dataset had read depth of over 5000 across almost the complete genome, except for the region corresponding to bases 1-24 of the reference sequence, within which read depth rose from 4 to >1000.

### Comparison of MRI103 serum and culture derived sequences

The consensus MRI103 sequences derived by next-generation sequencing (NGS) from serum and cell culture were almost identical. The serum virus consensus was 12292 nt in length and the cultured virus was 12296 nt, excluding a 3’ poly-A tail which started at the same position in both sequences and resulted from in vitro polyadenylation for oligo dT-primed cDNA synthesis. Among the 12292 aligned bases, the consensus sequence derived from the cultured virus differed at only 4 positions, compared with the serum consensus sequence (A/T, 1485; G/A, 1504; G/A, 2532; C/A, 2712; supplementary data table 2) all of which were non-synonymous (NS) changes, encoding distinct amino acids in glycoproteins Erns (1485, 1504) and E2 (2532, 2712).

To compare the NGS MRI103 genome sequences with genome sequences produced from PCR products, the first-strand cDNA used for NGS of the MRI103 serum and P3 viruses was used for genome amplification by RT-PCR using primers designed to amplify a range of Pestivirus A genotypes as a set of 24 overlapping PCR products (Table 1). The assembly of sequence data from overlapping PCR products, including the 5’UTR sequence amplified by primers 324-326, produced double-strand coverage of length 12043 nt, equivalent to nt 116-12159 of the MRI103 serum consensus. To further test the reliability of this approach, 100 µl of culture supernatant from passage 5 (P5) of MRI103 virus propagation was used as a source of viral RNA and 1 µl aliquots were used for RT-PCR. The P5 virus double-stranded sequence coverage was 12023 nt, twenty bases shorter than the serum and P3 sequences. A consensus sequence was derived from each RNA source for comparison with the NGS-derived data. Both PCR-derived consensus sequences were identical to the respective NGS-derived consensus and the P5 PCR consensus was identical to the P3 NGS consensus. The length of each PCR sequence was shorter than the NGS equivalent due to the positions of the leftmost and rightmost primer sites and the extent of reliable (double-stranded) coverage of the terminal fragments but each amplicon-derived consensus sequence included the entire polyprotein CDS.

Analysis of the MRI103 genome sequence by BLAST identified Pestivirus A strain Nose (AB078951) as the most similar complete genome currently available, with 93 % identity across 99 % of the genome. Translation of the MRI103 genome indicated a single continuous open reading frame (nt 372 - 12068 in the serum reference dataset) encoding 3898 amino acid residues which also had greatest similarity to strain Nose with 95 % amino acid identity.

### SNP analysis

SNP calling was done using Seqman Pro and all SNP were verified by visual inspection. This approach has been used previously to improve the reliability of SNP verification, and is especially applicable to short genomes such as RNA viruses (52). Diversity within each NGS genome assembly was measured using Shannon entropy (H), which was calculated at each candidate polymorphic base position in each sequence assembly (serum, culture and NADL). The MRI103-dT (serum) sequence had 824 SNP positions, with H ranging from the calculated error threshold of 0.0384 to 0.73. In comparison, the MRI-P3-dT culture assembly had 407 SNP positions, with values of H between 0.05 - 0.69; while the NADL-dT sequence had only 12 SNP positions with a much narrower H range of 0.10 to 0.23. Analysis of the distribution of H values in each sequence (Fig. 1) suggests that the number of low entropy variants reduced during culture, while the diversity of the remaining variants increased. This is supported by the observation that median entropy values increased (Fig. 1) from MRI103-dT serum (0.07), to MRI-P3-dT (0.11) to NADL-dT (0.14). In addition, Mann-Whitney U-test analysis of the two MRI103 distributions in Fig. 1 showed that they were significantly different from each other (P<0.0001). In addition, the total Shannon entropy in each genome (based on all positions above the error threshold) decreased with culture, from 92.9 in the serum virus, to 64.1 in the passage 3 culture virus and 1.8 in the NADL strain.

**Figure 1.**
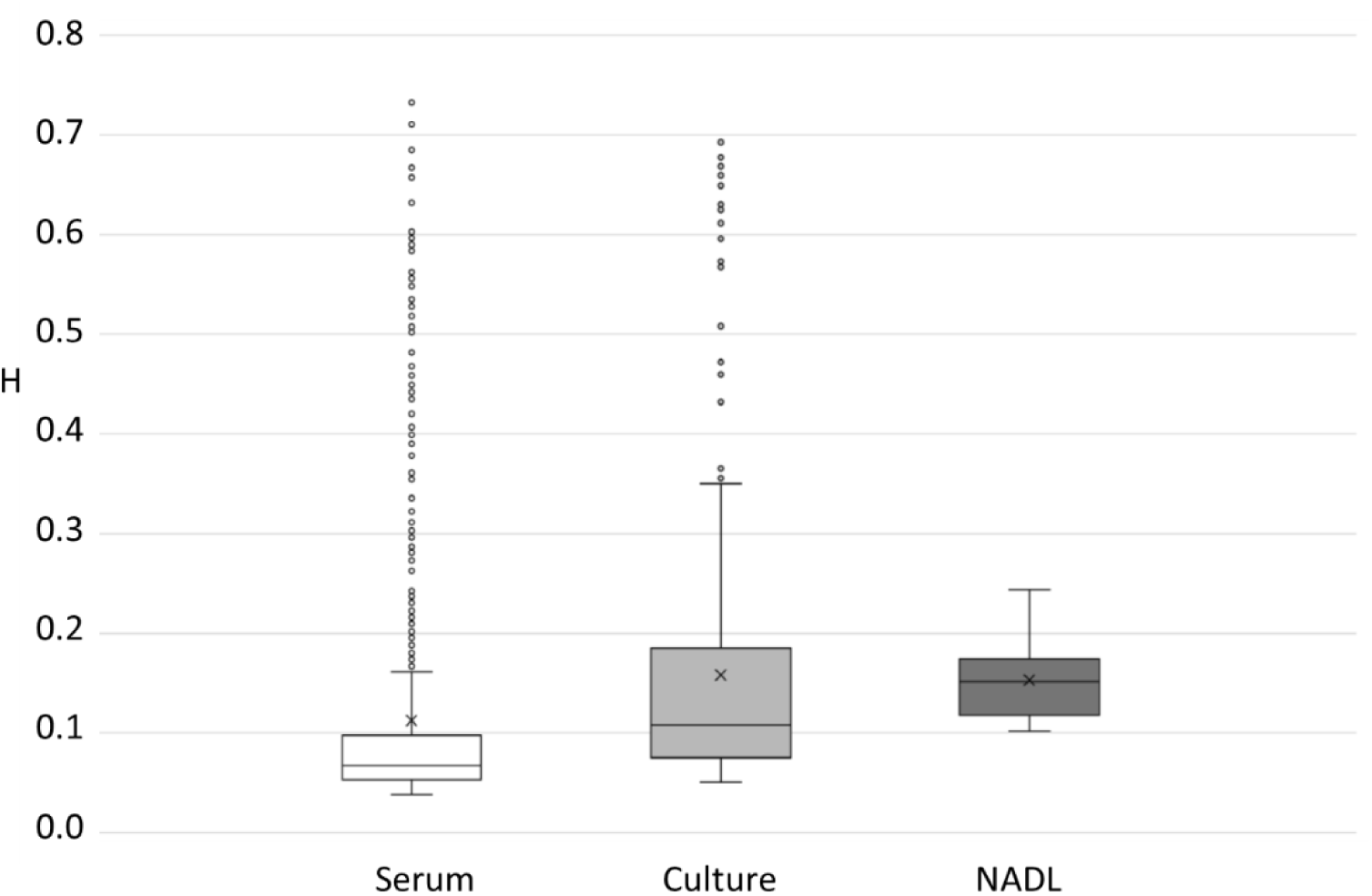
Boxplots of the Shannon Entropy values (H) for qualifying SNPs in each of the three genome sequences: MRI103-serum (open boxes, 824 SNP); MRI103-culture (light grey, 407 SNP) and NADL (dark grey, 12 SNP). For each plot the first and fourth quartiles are given as bounded vertical lines, the second and third quartiles are boxes separated by a line representing the median value, outliers are included as circles and the mean value is marked with a cross (x).

In these datasets, allele frequencies paralleled entropy values (supplementary data table 2), with MRI103-dT SNP ranging from 0.6 % to 46 %; MRI-P3-dT SNP ranging from 1 % to 41 % and NADL-dT ranging from 2.1 % to 6.3 %, in comparison with the respective consensus sequences. Indeed, where the consensus base changed in the MRI103 culture virus (positions 1485, 1504, 2532 and 2712), the frequencies of the alternate base were between 75 and 83 % (supplementary data table 2) relative to the serum consensus sequence.

Among the NGS data for MRI103, 681 SNP were found only in the serum sequence, 143 SNP positions were shared and 264 were found only in the culture sequence. Among the shared SNP, neither serum nor culture dataset had consistently higher allele frequencies suggesting that there was no consistent effect of culture passage on allele frequency.

The distribution of SNPs also varied across the MRI103 genome. Among the protein coding regions, SNPs were more frequent in the genes encoding the envelope proteins Erns, E1 and E2, compared to the non-structural genes (Fig. 2), in both the serum-derived virus and following propagation. Comparison of the Erns, E1 and E2 coding regions with the non-structural genes (NS2-NS5B) showed the glycoprotein gene sequences had significantly higher average single-base entropy values (p<0.01; Fig. 2) demonstrating a higher degree of polymorphism.

**Figure 2.**
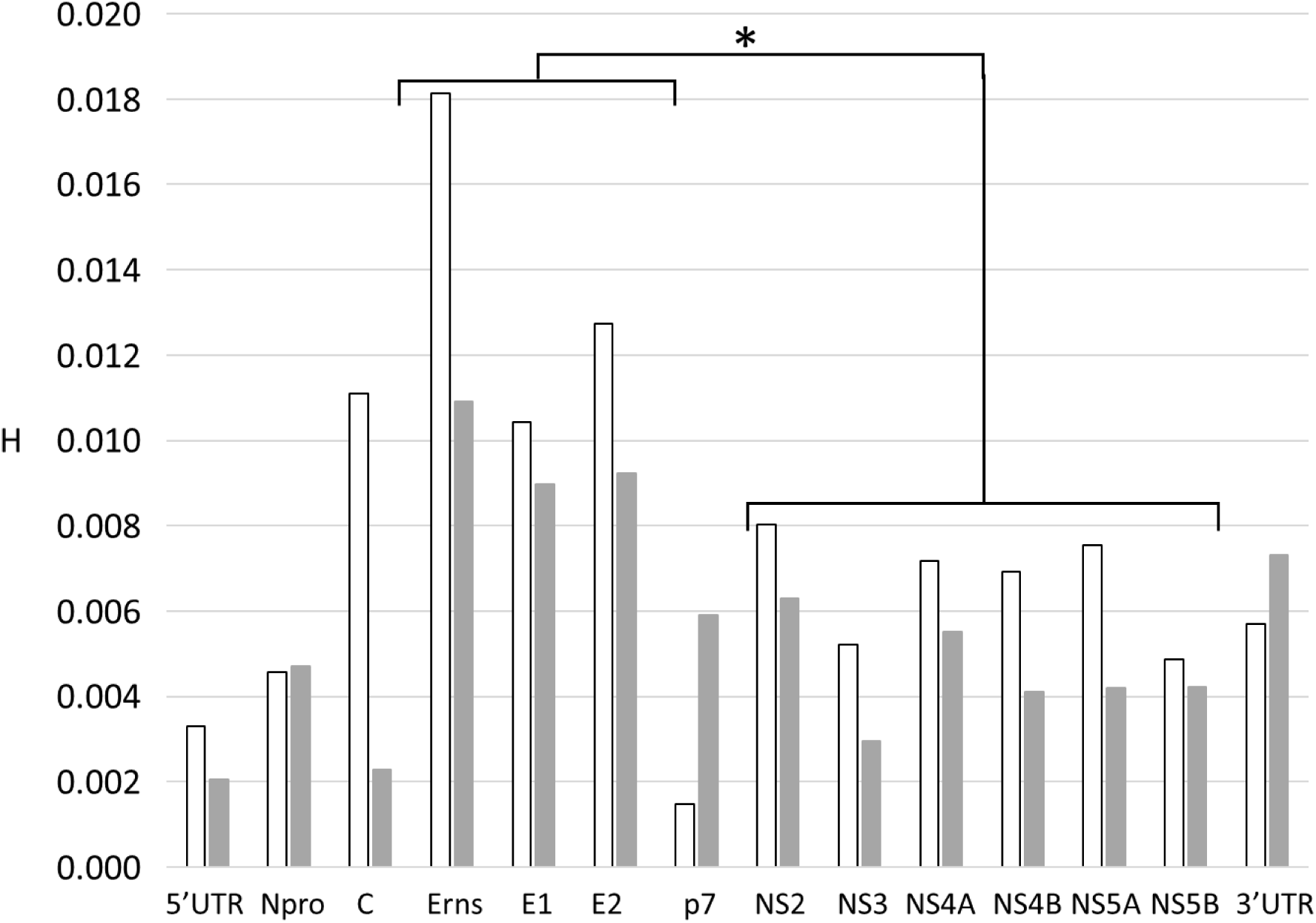
Distribution of sequence variation across the MRI103 genome. For each gene, the average single nucleotide entropy (H) was plotted. The columns are coloured as Fig 1: open = MRI103-serum; light grey = MRI103-culture. Student’s T-test comparison of the glycoprotein genes (Erns, E1, E2) and the non-structural genes (NS2-NS5B) showed that average entropy among the glycoprotein genes was significantly higher than the NS genes for both the serum and the cultured virus (p<0.01) *.

It was also clear that both synonymous (S) and non-synonymous (NS) SNP were more frequent in the MRI103 serum dataset than the culture dataset (p<0.005; Student’s T-test). There was also evidence to suggest that NS SNP in Erns, E1 and E2 were more frequent than in non-structural genes (NS2-NS5B). In the P3 culture dataset, NS SNP were more than twice as frequent in the envelope genes (p<0.01; Student’s T-test), while in the serum dataset NS SNP were also more frequent but this difference was not significant (p=0.055; Student’s T-test).

The SNP within the MRI103 datasets included 23 that encoded stop codons (19 in MRI103-dT and 4 in MRI-P3-dT, none shared; supplementary data table 2). All had minor allele frequencies below 3%, except for a single polymorphism (GAG>TAG nt 2283, within E1) that was found at 29% in the serum dataset.

### Quasispecies variation observed in PCR derived sequence

Examination of the sequence trace data from the PCR-derived genome assemblies also suggested that some quasispecies variation could be detected. Although each Sanger-derived consensus PCR sequence was identical to the respective NGS consensus, the PCR data also contained evidence of multiple base calls at many positions in the trace data.

The trace data was therefore analysed more closely to identify sites that exhibited clear quasispecies variation. Each base position within the PCR derived sequences that contained evidence of multiple base calls in at least two reads was recorded as either ambiguous (two base calls of approximately equal peak height, denoted by IUPAC ambiguity codes in supplementary data table 2); or as one major and one minor base call (one base having a higher peak in all reads, denoted as uppercase/lowercase (e.g. G/a) in supplementary data table 2).

Examples of sequencing traces from polymorphic positions (including two that represent consensus base changes between serum and cultured viruses) are shown in Fig. 3 and relate to SNP positions detailed in supplementary data table 2. The traces show either reducing diversity (nt 1504), diversity throughout (nt 2532) or increasing diversity (nt 4263) with culture passage. Position 1504 was called as R (G/A) within the serum PCR sequence but A was called as the majority base in both P3 and P5 sequences. At position 2532, all three virus sequences had evidence of distinct base calls but the majority base call appeared to change from G in serum to A in culture. In contrast, at position 4263, the serum virus had A as the majority base call, while the minority G call became more prominent in P3 and was called as R in P5.

**Figure 3.**
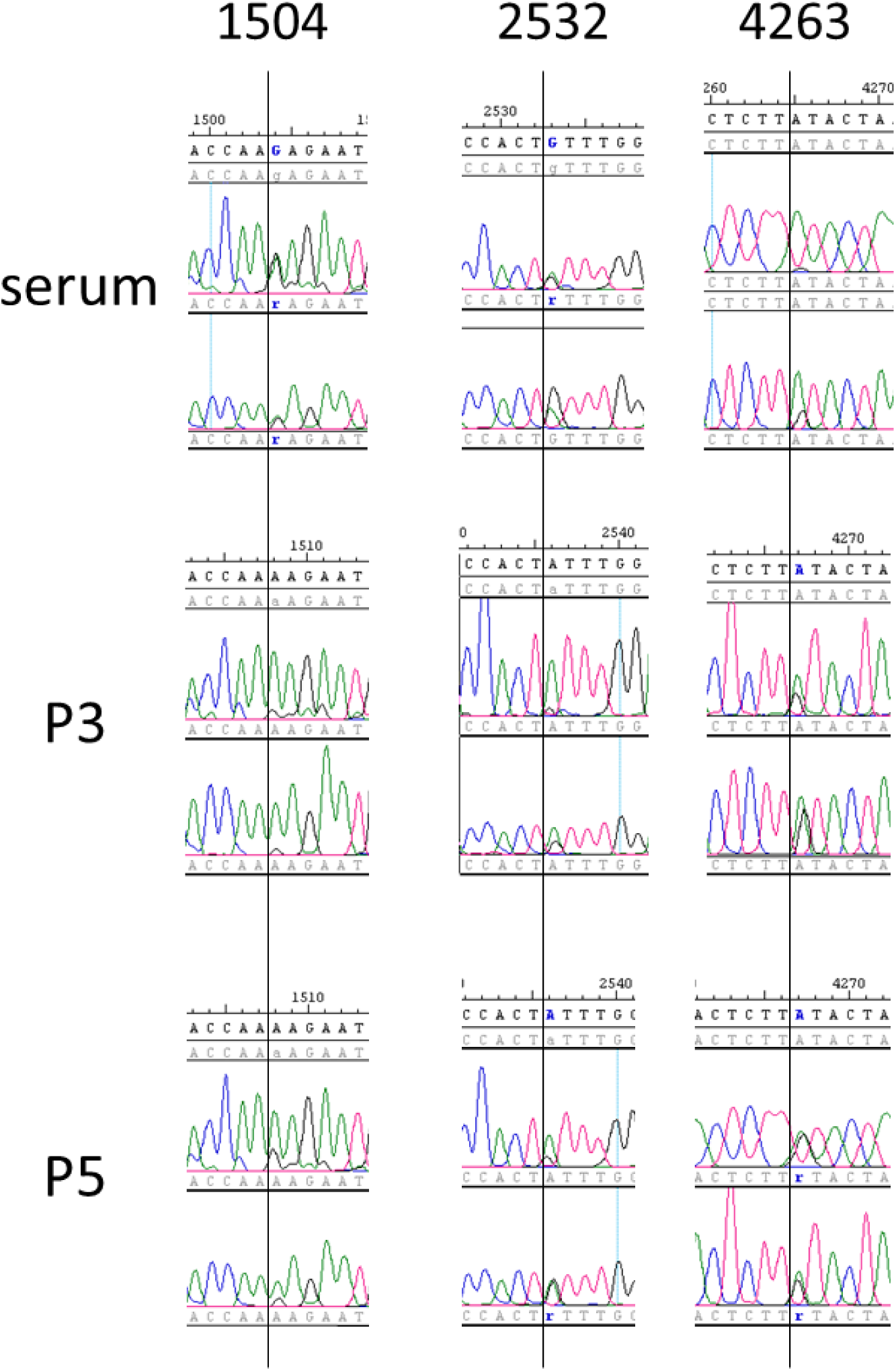
Example traces from PCR amplicon sequencing of MRI103 from serum, passage 3 (P3) and passage 5 (P5) of culture. In each example two traces from the equivalent position in each virus sequence are shown. The number above each set of traces refers to the SNP coordinates given in supplementary data table 2. The local consensus sequence and numbering is given in the top line of each panel, while base calls for each trace are given below the trace. Software-assigned potential SNP in each read and in the consensus are highlighted in blue. Additional lines containing read data were collapsed to simplify the figure.

Overall, in the serum PCR sequence, 58 positions were called as polymorphic, including 10 that were not seen in the serum NGS dataset. Among the remaining 48 positions, the corresponding SNP frequency in the serum NGS dataset varied from 4.9% to 46%. These included 68% of serum NGS SNP with frequency >10%.

Within the P3 culture PCR sequence, 33 positions were called as polymorphic, all of which were variable in the NGS data, with SNP frequency ranging from 5% to 80%. These positions included 80% of P3 culture NGS SNP with frequency >10%.

For the P5 culture PCR sequence, 57 positions were called as polymorphic. In the absence of a P5 NGS dataset, these were compared with the P3 NGS data. This showed that 12 SNP were not found in the P3 culture NGS data. Corresponding SNP frequencies for the remaining 45 SNP positions in the MRI-P3-dT dataset varied from 1.4% to 91%, including 73% of P3 culture NGS SNP with frequency >10%. The P5 PCR dataset was clearly distinct from the serum and P3 sequences and contained 17 SNP at positions that had variant frequencies below 5% in the P3 culture NGS dataset. This may reflect further changes in quasispecies structure between P3 and P5, with some variant bases becoming more highly represented in the P5 virus population.

## Discussion

The number of complete genome or polyprotein sequences for pestiviruses that are publicly available now exceeds 100 for each of Pestivirus A and B (European Nucleotide Archive: www.ebi.ac.uk/ena). Of these, most were sequenced by Sanger methods after cloning or amplification of genome segments while many of the more recently determined sequences were defined by NGS approaches. In almost all cases, however, only the consensus sequences have been published and quasispecies variation has seldom been examined. For example, Workman and co-workers (36) defined genome sequences of 19 BVD virus strains directly from plasma. These strains were genotyped within Pestivirus A and B species but information on quasispecies variation within these strains was not presented or discussed. Similarly, Neil and co-workers (40) used NGS approaches to sequence 36 virus isolates representing a range of Pestivirus B (BVDV type 2) strains but this work did not examine quasipecies diversity.

The quasispecies nature of RNA viruses may facilitate their transmission between host species by providing a reservoir of variants for selection to operate on (53, 54). This process, whereby a subpopulation of viruses can infect and adapt to a different host, is extremely important for the emergence of novel or more virulent viruses and also may lead to vaccine or diagnostic “escape mutants”. Understanding the baseline quasispecies structure of RNA virus populations may allow the risks of acquiring a wider host range or increased virulence to be assessed in advance.

Specific studies of BVD virus quasispecies variation have generally focused on individual genes, by counting clones of PCR products reflecting sequence variation within an E1/E2 fragment (55) or in the E2 and NS5B genes (56). Other studies have compared the virus consensus sequences between sequentially-infected individuals, inferring the nature and rates of quasispecies variation between hosts (39).

Recent work described a PCR and NGS approach to define consensus sequences of major genes and infer detailed phylogenetic relationships among Canadian BVD virus strains, but without analysing quasispecies variation (57). The same group used RT-PCR amplification of complete genomes from lymph node tissue to study quasispecies diversity in different BVD strains (58). This work found more variation in structural genes, in accord with results published here (Fig. 2). However, this study did not include unamplified control material or examine the influence of PCR amplification on the datasets obtained.

In this study we used a PCR-free approach to derive NGS datasets for the MRI103 strain of Pestivirus A from serum and after three passages in culture, obtaining both consensus sequences and genome level quasispecies data. We also analysed the laboratory strain NADL. The serum MRI103 virus had the largest number of variant sites within its genome, while the P3 cultured and NADL genomes showed decreasing numbers of polymorphic sites. However, the median Shannon entropy value increased with culture (Fig. 2), suggesting that the remaining variants in cultured viruses had higher individual entropy values and were represented at higher frequency within the populations. The view that overall virus diversity decreases with culture was supported both by the number of variant sites in each genome (824 in serum, 407 in P3 and 12 in NADL) and by the total entropy in each genome (sums of 93 in serum, 64 in P3 and 1.8 in NADL). The summed entropy figures may give a more accurate picture of the loss of diversity in culture, reflecting both the number of sites above the error cutoff and the level of diversity at each site.

The observation of multiple stop codons at low frequency within the MRI103 NGS datasets and a single high frequency mutation within E1 (nt 2283; 29% T) in the serum virus, raises the question of their potential function. Genomes carrying stop codons may be defective or may express truncated gene products. Genomes with polyprotein truncation may enhance the survival of the population as a whole, or be rescued by translational frameshifting, the use of alternative start codons or by complementation in cells infected with multiple viruses. Some stop codon mutations may benefit persistent infection by reducing the expression of viral antigens (59, 60), although increasing the frequency of stop codon mutations in RNA viruses was associated with significant losses in viral fitness both *in vivo* and *in vitro* (61). The observations here, that 19 stop codon variants were found in the serum virus sequence, while only four stop codon variants were seen in the P3 culture virus sequence and that none of these were shared (supplementary data table 2), support the view that the stop codon variants did not confer an advantage in culture and were lost. It will be of interest to analyse further samples *ex vivo* to determine whether high frequency stop codon variants are a common feature of pestivirus quasispecies populations.

The four consensus base changes in the cultured virus sequence were also those that had the greatest change in SNP frequency across the entire genome (supplementary data table 2). The changed consensus bases encoded amino acid changes to the virus glycoproteins Erns (nt 1485=T102S; 1504=R108K) and E2 (2532=V62I; 2712=Q122K) and in each case the substitutions lay within the exposed domains of the proteins (reviewed in 62). For Erns, the altered residues were located between the catalytic domain (residues 32-81), which has RNase activity, and the C-terminal region (residues 167-227) which mediates membrane retention and processing. These residues have not been recorded as being associated with any recognised antibody epitope (62) and the nature of the changes (Thr-Ser; Arg-Lys) maintained the respective hydrophilic and basic character of the residues and therefore may not have significant effects on Erns structure-function.

In E2 the substitutions were in the two outermost (membrane-distal) domains that possess Ig-like folds and the substitution in the outermost domain (V61I) is close to a recognised neutralising motif in E2 of Classical Swine Fever Virus (residues 64-76), which has analogous structure (62). However, like Erns, the substitutions in E2 appeared to be conservative in nature and may not cause any clear change in structure or function of E2.

However the observation that mutations in the Erns glycoprotein gene can lead to failure of diagnostic assays implies that serum-derived and culture-derived samples of the same virus could give different results in such assays (63).

Although the overall degree of variation within the cultured virus was lower than the serum virus genome, it was clear from analysis of shared SNP that culture passage did not lead to a consistent reduction in the degree of variation of individual SNP, suggesting that culture may select for increased frequency of some SNP variants but not others. In this context we also note that the twelve variant positions within the NADL genome, with SNP frequencies of 2.1 - 6.3 %, did not appear to correspond with SNP positions in the MRI103 genome, when the two genome sequences were aligned (supplementary data table 2). This may suggest that extended culture exerts selective pressure *in vitro* on sites that are not under such pressure *in vivo*.

In addition, Sanger sequencing of MRI103 RT-PCR products produced the same virus consensus sequences as the NGS analysis and identified many polymorphic sites, suggesting that PCR amplification has potential for use in quasispecies analysis. While this may be more error-prone that an amplification-free approach, it clearly has value where the amount or complexity of input material makes direct sequencing difficult. The use of PCR and clone counting approaches (55, 56) to assess quasispecies variation supports the view that read-counting within NGS data from such PCR products may yield accurate quantitative data. This suggests that analysis of quasispecies variation in pestiviruses (and potentially other RNA viruses) could focus on uncultured samples from infected individuals rather than after virus isolation, potentially taking advantage of amplification methods to derive genome level quasispecies data.

From the Sanger data, it appeared that the quasispecies variants detected in the MRI103 P5 virus included a number that were derived from low-frequency variants in the P3 virus and others that were not detected in the serum or P3 datasets. This appears to be in keeping with the suggestion above that culture exerts selective pressure on different sites from those selected *in vivo*, in addition to reducing the overall quasispecies variation in the cultured virus population.

Drivers that may act to fix sequence changes include selection pressures resulting from infection of new cell types within a host, of new individuals or of new species. Where available, data on cross-species infection suggests that success may be facilitated by the presence of ‘pre-adapted’ variants that could be selected and rapidly established in the new host (23, 54). The appearance of host-specific sequence changes in goats, sheep and cattle serially infected with BVD viruses supports this view (22-24). In addition, NGS studies of bird-to-mammal transmission of avian influenza viruses indicates that transmission to a new host creates genetic bottlenecks that select variants with suitable adaptations (53, 54). The existence of virus sub-populations within individuals is also supported by studies of HIV, where distinct, organ-specific populations were found and the differences between populations increased with time (64).

The sequencing of the NADL strain here showed that it had identical consensus sequence to the reference genome sequence determined in 1988 (29). Additionally, the NADL dataset had a lower number of variant base positions than MRI103 P3 (12 compared to 407), but these few variants had higher median diversity than either MRI103 dataset (Fig. 1; supplementary data table 2). Thus, this highly culture-adapted virus had few variant positions but these appeared to be maintained stably.

The PCR-based Sanger sequence data obtained here reflected the quasispecies variation detected by NGS, but SNPs with frequency <10% in the corresponding NGS dataset appeared to be more difficult to detect, reflecting the more qualitative nature of Sanger sequencing. The application of NGS analysis to PCR amplicon sequencing could be an easy and cost-effective way to gather information on quasispecies variation and allow evolution within chains of infection to be monitored from even small serum or tissue samples. Such a forensic use of sequencing could also be applied in situations such as the late stages of BVD eradication campaigns, where single locus sequencing may not distinguish similar viruses shared across multiple premises from true transmission events. However, it is clear that for such forensic epidemiology we will also need to improve our understanding of virus sequence variation within a chain of infection and between animals infected by the same source. The recently described Scottish BVD Biobank (43) includes such samples that could be analysed to improve our understanding of what defines a single Pestivirus A strain.

While it is clear that the amplification-free process used here, offering a no- or low-bias protocol for producing RNA virus templates for NGS, may provide a gold-standard approach for analysis of quasispecies diversity, it has the disadvantage of requiring a high volume of input material. This may however be alleviated by ongoing technical improvements in the generation of quantitative NGS datasets from small amounts of RNA. In addition, the use of PCR as a first step in quasispecies analysis needs further validation and comparison with direct NGS. Analysis of potential problems, such as uneven coverage, especially at the ends of target viruses, will help critically evaluate this potentially valuable approach. Previous work (65) used PCR to amplify the FMDV genome prior to NGS library preparation to investigate quasispecies variation in the inoculum virus and in lesions from same animal. While this approach accurately replicated the consensus inoculum reference sequence, it did not address potential limitations of the approach and there was no NGS-only dataset for comparison. It also demonstrated that the high read-depth achievable by using NGS of PCR products allowed definition of quasispecies variants with frequencies below 1%.

## Conclusion

This work provides evidence that initial culture of a BVD virus isolate drives virus evolution, producing changes in the virus consensus sequence. Here, four clear changes in MRI103 virus consensus sequence, encoding amino acid sequence changes in the Erns and E2 glycoproteins, were found after only 3 passages in culture, while 15 novel variant sites were detected in the virus after 5 passages. This was accompanied by a clear reduction in quasispecies diversity with culture, as defined by number of variant sites and total entropy, suggesting that genome sequences derived after virus isolation may not present an accurate picture of the quasispecies variation *in vivo* or the true virus consensus sequence. The very low total entropy of variant sites in the NADL genome analysed here suggests that extensive culture works to maintain a genome sequence that is well-adapted to the tissue culture environment.

While this study has demonstrated clear changes in quasi-species structure after only three passages in culture, additional experiments could enhance the future value of this work. Repetition of the culture passages and quasispecies analysis would provide useful information on the reproducibility of the changes induced by culture, while infection of BVD-naive animals with the cultured virus could help define whether the changes were reversible *in vivo*. Further analysis of genetic variation within culture-adapted viruses, in homologous and heterologous culture systems, may also help us understand how this low level of genetic variation is maintained. In any case, the clonal nature of the NADL strain compared to the impressive diversity of the virus in a BVD PI animal implies that using highly passaged lab strains to model natural infection may not be representative.

We also showed that PCR amplification of viral RNA produces amplicons in which many viral polymorphisms appear to be conserved. Further quantitative analysis, with and without PCR amplification, will help define how closely genome sequence diversity in PCR amplicons reproduces that obtained directly from viral RNA. However, the data produced here suggest that even Sanger sequence analysis of PCR-amplified virus genome fragments can reflect the quasispecies variation found in the source material, at least for more frequent variants. The combination of PCR amplification and NGS analysis of the quasispecies structure of virus genomes may be a powerful approach to allow investigation of samples that would otherwise be difficult or impossible to study.

## Author contributions

George C. Russell: conceptualisation, investigation, validation, formal analysis, resources, data curation, writing (original draft preparation; review and editing), visualisation, supervision, project administration, funding

Ruth N. Zadoks: conceptualisation, writing (review and editing), visualisation, supervision

Kim Willoughby: conceptualisation, resources, writing (review and editing), supervision

Claudia Bachofen: conceptualisation, investigation, validation, formal analysis, resources, data curation, writing (original draft preparation; review and editing), visualisation, funding

## Acknowledgements

We thank The Genome Analysis Centre (now Earlham Institute) for the provision of sequencing facilities under their Capacity and Capability Challenge programme (project CCC-123). We are grateful to staff of the Scottish Centre for Production Animal Health & Food Safety, University of Glasgow for facilitating the collection of BVD-positive blood samples. We are also grateful to the Moredun Virus Surveillance Unit for the provision of strains and support.

## Data Bibliography

1 European Nucleotide Archive (www.ebi.ac.uk/ena/)

