## supplementary data for "Bovine viral diarrhoea virus loses quasispecies diversity rapidly in culture"

### Supplementary data Table 1

Summary data from reference-led NGS assembly of the six datasets (NADL-dT; NADL-B12; MRI103-dT; MRI103-B12; MRI-P3-dT and MRI-P3-B12) using Seqman Ngen (DNASTAR Inc, Madison, WI, USA). All datasets assembled to their respective reference sequence with high quality and good consistency in the placing of read pairs. -dT and -B12 indicate the primer used for cDNA synthesis.

| sample | Total reads | Assembled reads | Average Coverage: | Average Quality (Assembled Sequences) | Assembled Pairs: | Pairs Consistent Within a Contig: | Minimum Match Percentage: |
|---|---|---|---|---|---|---|---|
| NADL-dT | 3587795 | 578995 | 10708 | 36 | 254788 | 253507 | 93 |
| NADL-B12 | 4740735 | 50141 | 793 | 36 | 10335 | 10324 | 93 |
| MRI103-dT | 3577799 | 11492 | 220 | 36 | 5281 | 5263 | 93 |
| MRI103-B12 | 5064217 | 2459 | 40 | 36 | 310 | 309 | 93 |
| MRI-P3-dT | 3491349 | 115739 | 2024 | 36 | 39242 | 39190 | 93 |
| MRI-P3-B12 | 5328689 | 15096 | 236 | 35 | 2739 | 2694 | 93 |

**Supplementary data Table 2**: Variant base sites within the BVDV MRI103 and NADL genomes, as sequenced directly from serum or culture fluid, or following PCR amplification. Column headings are as follows:

| | |
|---|---|
| Ref Position | Base position on reference sequence (serum NGS consensus) |
| Ref Base | Base encoded by reference sequence at this position |
| Serum Base | Variant base found in serum NGS dataset |
| Serum SNP % | Frequency of non-consensus base in serum NGS dataset |
| Serum Depth | Read depth at this position in serum NGS dataset |
| Serum entropy | Shannon Entropy at this nucleotide position calculated as H= -Σ(freqA*ln(freqA) +freqC*ln(freqC)+freqG*ln(freqG)+freqT*ln(freqT)), where freqN= fractional frequency of the specified base. |
| Culture Base | Variant base found in culture NGS dataset |
| Culture SNP % | Frequency of non-consensus base in culture NGS dataset |
| Culture Depth | Read depth at this position in culture NGS dataset |
| Culture entropy | Shannon Entropy at this nucleotide position. |
| Features | BVDV coding regions |
| S/NS | Nature of substitutions: S, synonymous; NS, non-synonymous; STOP, creating stop codon; NA, not applicable (substitution in untranslated region). |
| Residue change | Nature of change from consensus residue to altered residue (one-letter amino acid code; * = stop codon) |
| Serum PCR call | Ambiguous base calls detected in serum based PCR sequence. Bases are indicated as IUPAC ambiguity codes where trace values were similar for each base and as uppercase/lowercase pairs where the uppercase base had higher peak height in all traces of this position. |
| P3 PCR call | Ambiguous base calls detected in P3 culture based PCR sequence. |
| P5 PCR call | Ambiguous base calls detected in P5 culture based PCR sequence. |
| NADL position | Base position on NADL sequence (M 31182) |
| MRI103 position | Base position on MRI103 reference sequence (when aligned with NADL) |
| NADL base | Consensus base found in NADL NGS dataset |
| variant Base | Variant base found in NADL NGS dataset |
| NADL SNP % | Frequency of non-consensus base in NADL NGS dataset |
| NADL Depth | Read depth at this position in NADL NGS dataset |
| NADL entropy | Shannon Entropy at this nucleotide position |

| Ref Position | Ref Base | Serum Base | Serum SNP % | Serum Depth | Serum entropy | | Culture Base | Culture SNP % | Culture Depth | Culture entropy | | Features | S/NS | Residue change | Serum PCR call | P3 PCR call | P5 PCR call |
|---|---|---|---|---|---|---|---|---|---|---|---|---|---|---|---|---|---|
| 41 | T | | | | | | C | 2.8% | 1945 | 0.1299 | | UTR | NA | | | | |
| 58 | C | | | | | | T | 1.5% | 3239 | 0.0801 | | UTR | NA | | | | |
| 67 | C | A | 1.5% | 261 | 0.0665 | | | | | | | UTR | NA | | | | |
| 69 | A | C | 14.5% | 269 | 0.4104 | | | | | | | UTR | NA | | | | |
| 118 | G | A | 1.0% | 416 | 0.0542 | | | | | | | UTR | NA | | | | |
| 120 | G | T | 7.4% | 419 | 0.2638 | | T | 5.7% | 3723 | 0.2215 | | UTR | NA | | | | |
| 154 | G | A | 0.7% | 570 | 0.0418 | | | | | | | UTR | NA | | | | |
| 202 | C | A | 1.3% | 752 | 0.0796 | | | | | | | UTR | NA | | | | |
| 203 | G | A | 0.8% | 752 | 0.0465 | | A | 4.1% | 6074 | 0.1759 | | UTR | NA | | | | |
| 207 | A | | | | | | G | 1.5% | 6135 | 0.0824 | | UTR | NA | | | | |
| 237 | C | T | 1.7% | 772 | 0.0814 | | | | | | | UTR | NA | | | | |
| 270 | G | A | 1.0% | 714 | 0.0551 | | | | | | | UTR | NA | | | | |
| 277 | C | | | | | | T | 1.4% | 4155 | 0.0737 | | UTR | NA | | | | |
| 278 | G | A | 0.8% | 658 | 0.0447 | | | | | | | UTR | NA | | | | |
| 292 | C | T | 0.7% | 557 | 0.0426 | | | | | | | UTR | NA | | | | |
| 305 | T | A | 0.9% | 567 | 0.0437 | | | | | | | UTR | NA | | | | |
| 384 | A | G | 1.0% | 406 | 0.0461 | | | | | | | Npro | NS | TA | | | |
| 434 | T | A | 1.4% | 140 | 0.0749 | | | | | | | Npro | NS | DE | | | |
| 452 | G | A | 20.5% | 156 | 0.5074 | | A | 8.2% | 2473 | 0.2838 | | Npro | S | | | | |
| 458 | C | | | | | | T | 1.9% | 2502 | 0.0948 | | Npro | S | | | | |
| 464 | T | C | 1.3% | 152 | 0.0701 | | | | | | | Npro | S | | | | |
| 478 | G | A | 1.3% | 157 | 0.0682 | | | | | | | Npro | NS | RK | | | |
| 485 | A | G | 1.2% | 171 | 0.0637 | | | | | | | Npro | S | | | | |
| 504 | C | T | 1.1% | 184 | 0.0600 | | | | | | | Npro | S | | | | |
| 509 | G | | | | | | A | 1.0% | 2843 | 0.0595 | | Npro | S | | | | |
| 525 | G | A | 1.0% | 198 | 0.0565 | | | | | | | Npro | NS | GR | | | |

| Ref Position | Ref Base | Serum Base | Serum SNP % | Serum Depth | Serum entropy | | Culture Base | Culture SNP % | Culture Depth | Culture entropy | | Features | S/NS | Residue change | Serum PCR call | P3 PCR call | P5 PCR call |
|---|---|---|---|---|---|---|---|---|---|---|---|---|---|---|---|---|---|
| 526 | G | A | 1.0% | 203 | 0.0553 | | | | | | | Npro | NS | GE | | | |
| 531 | C | T | 1.0% | 206 | 0.0547 | | | | | | | Npro | NS | RC | | | |
| 533 | C | G | 1.0% | 207 | 0.0544 | | | | | | | Npro | S | | | | |
| 536 | T | C | 1.0% | 208 | 0.0542 | | | | | | | Npro | S | | | | |
| 542 | C | | | | | | T | 4.2% | 2918 | 0.1822 | | Npro | S | | | | |
| 558 | T | C | 1.0% | 193 | 0.0577 | | | | | | | Npro | S | | | | |
| 560 | G | T | 1.5% | 200 | 0.0609 | | | | | | | Npro | NS | LF | | | |
| 584 | G | A | 0.9% | 223 | 0.0512 | | | | | | | Npro | S | | | | |
| 591 | G | | | | | | A | 1.1% | 2628 | 0.0590 | | Npro | NS | GS | | | |
| 592 | G | A | 6.4% | 220 | 0.2369 | | A | 4.6% | 2641 | 0.1852 | | Npro | NS | GD | | | G/a |
| 626 | A | C | 0.8% | 252 | 0.0463 | | | | | | | Npro | S | | | | |
| 646 | A | T | 0.7% | 286 | 0.0417 | | | | | | | Npro | NS | DV | | | |
| 653 | G | | | | | | A | 6.4% | 2541 | 0.2425 | | Npro | S | | | | |
| 656 | T | G | 6.7% | 312 | 0.2466 | | G | 28.2% | 2562 | 0.6249 | | Npro | S | | | T/g | |
| 686 | C | | | | | | A | 1.4% | 3766 | 0.0751 | | Npro | S | | | | |
| 692 | G | A | 0.8% | 355 | 0.0488 | | | | | | | Npro | S | | | | |
| 707 | T | C | 1.4% | 361 | 0.0730 | | | | | | | Npro | S | | | | |
| 713 | G | A | 1.9% | 363 | 0.0952 | | | | | | | Npro | S | | | | |
| 767 | T | | | | | | C | 2.9% | 3865 | 0.1365 | | Npro | S | | | | |
| 797 | G | | | | | | A | 11.1% | 3857 | 0.3555 | | Npro | S | | | | |
| 812 | T | | | | | | C | 1.3% | 3958 | 0.0702 | | Npro | S | | | | |
| 815 | C | T | 1.1% | 359 | 0.0612 | | | | | | | Npro | S | | | | |
| 846 | C | T | 0.9% | 329 | 0.0401 | | | | | | | Npro | S | | | | |
| 860 | A | G | 1.0% | 312 | 0.0419 | | | | | | | Npro | S | | | | |
| 866 | C | T | 0.6% | 311 | 0.0389 | | | | | | | Npro | S | | | | |
| 883 | C | T | 0.7% | 301 | 0.0399 | | | | | | | C | NS | TI | | | |

| Ref Position | Ref Base | Serum Base | Serum SNP % | Serum Depth | Serum entropy | | Culture Base | Culture SNP % | Culture Depth | Culture entropy | | Features | S/NS | Residue change | Serum PCR call | P3 PCR call | P5 PCR call |
|---|---|---|---|---|---|---|---|---|---|---|---|---|---|---|---|---|---|
| 888 | G | A | 0.7% | 299 | 0.0402 | | | | | | | C | NS | DN | | | |
| 899 | A | G | 0.8% | 249 | 0.0467 | | | | | | | C | S | | | | |
| 902 | A | G | 1.2% | 242 | 0.0667 | | | | | | | C | S | | | | |
| 911 | A | G | 2.5% | 239 | 0.1173 | | | | | | | C | S | | | | |
| 914 | A | G | 0.8% | 244 | 0.0475 | | | | | | | C | S | | | | |
| 929 | A | G | 0.9% | 231 | 0.0497 | | | | | | | C | S | | | | |
| 938 | G | A | 0.9% | 229 | 0.0501 | | | | | | | C | S | | | | |
| 958 | C | G | 0.8% | 245 | 0.0474 | | | | | | | C | NS | PR | | | |
| 959 | C | T | 2.4% | 248 | 0.1293 | | | | | | | C | S | | | | |
| 979 | G | T | 1.6% | 253 | 0.0922 | | | | | | | C | NS | SI | | | |
| 983 | G | A | 1.6% | 252 | 0.0815 | | | | | | | C | S | | | | |
| 994 | C | T | 0.8% | 243 | 0.0477 | | | | | | | C | NS | PL | | | |
| 997 | A | T | 0.8% | 248 | 0.0469 | | | | | | | C | NS | DV | | | |
| 999 | G | T | 0.8% | 246 | 0.0472 | | | | | | | C | NS | AS | | | |
| 1007 | A | G | 0.8% | 250 | 0.0466 | | | | | | | C | NS | IM | | | |
| 1020 | G | T | 0.8% | 250 | 0.0466 | | | | | | | C | NS | VF | | | |
| 1022 | T | | | | | | C | 1.8% | 2736 | 0.0883 | | C | S | | | | |
| 1028 | C | G | 0.9% | 235 | 0.0490 | | | | | | | C | STOP | Y* | | | |
| 1029 | C | T | 1.3% | 232 | 0.0538 | | | | | | | C | STOP | Q* | | | |
| 1033 | T | A | 1.8% | 226 | 0.0749 | | | | | | | C | NS | VE | | | |
| 1036 | A | G | 5.9% | 220 | 0.2245 | | | | | | | C | NS | KR | | | |
| 1037 | G | A | 0.9% | 220 | 0.0518 | | | | | | | C | S | | | | |
| 1050 | G | A | 0.9% | 217 | 0.0524 | | A | 5.7% | 2500 | 0.2181 | | C | NS | VI | | | |
| 1054 | A | G | 1.9% | 209 | 0.0947 | | | | | | | C | NS | KR | | | |
| 1061 | A | | | | | | G | 1.1% | 2434 | 0.0610 | | C | S | | | | |
| 1063 | G | A | 0.9% | 233 | 0.0494 | | A | 3.8% | 2549 | 0.1681 | | C | NS | SN | | | |

| Ref Position | Ref Base | Serum Base | Serum SNP % | Serum Depth | Serum entropy | | Culture Base | Culture SNP % | Culture Depth | Culture entropy | | Features | S/NS | Residue change | Serum PCR call | P3 PCR call | P5 PCR call |
|---|---|---|---|---|---|---|---|---|---|---|---|---|---|---|---|---|---|
| 1070 | G | T | 1.7% | 235 | 0.0862 | | | | | | | C | NS | QH | | | |
| 1076 | T | | | | | | C | 1.3% | 2552 | 0.0730 | | C | S | | | | |
| 1079 | A | G | 0.8% | 241 | 0.0480 | | | | | | | C | S | | | | |
| 1096 | A | G | 0.9% | 221 | 0.0516 | | | | | | | C | NS | KR | | | |
| 1100 | G | A | 2.3% | 219 | 0.1089 | | | | | | | C | S | | | | |
| 1102 | C | A | 1.4% | 218 | 0.0726 | | | | | | | C | NS | PQ | | | |
| 1110 | C | T | 0.9% | 225 | 0.0508 | | | | | | | C | NS | RC | | | |
| 1112 | C | | | | | | T | 1.5% | 2633 | 0.0815 | | C | S | | | | |
| 1118 | A | G | 0.9% | 227 | 0.0505 | | | | | | | C | S | | | | |
| 1121 | T | C | 1.8% | 225 | 0.0893 | | | | | | | C | S | | | | |
| 1127 | A | G | 0.9% | 212 | 0.0534 | | | | | | | C | S | | | | |
| 1130 | A | T | 0.9% | 223 | 0.0512 | | | | | | | C | S | | | | |
| 1137 | G | A | 1.4% | 222 | 0.0558 | | | | | | | C | NS | AT | | | |
| 1139 | A | T | 1.4% | 217 | 0.0569 | | | | | | | C | S | | | | |
| 1140 | T | A | 0.9% | 216 | 0.0526 | | | | | | | C | NS | WR | | | |
| 1155 | G | A | 24.4% | 217 | 0.5559 | | | | | | | C | NS | VM | G/a | | |
| 1156 | T | C | 1.4% | 221 | 0.0718 | | | | | | | C | NS | VA | | | |
| 1159 | T | C | 1.4% | 222 | 0.0558 | | | | | | | C | NS | IT | | | |
| 1172 | T | C | 2.2% | 230 | 0.1047 | | | | | | | C | S | | | | |
| 1174 | C | T | 0.9% | 230 | 0.0499 | | | | | | | C | NS | TI | | | |
| 1190 | A | T | 0.9% | 220 | 0.0518 | | | | | | | Erns | S | | | | |
| 1217 | G | A | 1.0% | 202 | 0.0555 | | | | | | | Erns | S | | | | |
| 1220 | A | | | | | | G | 1.4% | 1718 | 0.0735 | | Erns | S | | | | |
| 1234 | A | G | 1.9% | 214 | 0.0929 | | | | | | | Erns | NS | QR | | | |
| 1237 | C | T | 2.8% | 217 | 0.1265 | | T | 3.0% | 1635 | 0.1397 | | Erns | NS | AV | | | C/t |
| 1243 | T | A | 0.9% | 220 | 0.0518 | | | | | | | Erns | NS | FY | | | |

| Ref Position | Ref Base | Serum Base | Serum SNP % | Serum Depth | Serum entropy | | Culture Base | Culture SNP % | Culture Depth | Culture entropy | | Features | S/NS | Residue change | Serum PCR call | P3 PCR call | P5 PCR call |
|---|---|---|---|---|---|---|---|---|---|---|---|---|---|---|---|---|---|
| 1244 | C | T | 0.9% | 217 | 0.0524 | | T | 7.5% | 1590 | 0.2660 | | Erns | S | | | C/t | |
| 1246 | A | G | 0.9% | 216 | 0.0526 | | | | | | | Erns | NS | QR | | | |
| 1257 | A | T | 17.4% | 218 | 0.6320 | | T | 5.8% | 1388 | 0.2846 | | Erns | NS | NY | | | |
| 1258 | A | G | 1.8% | 219 | 0.1039 | | | | | | | Erns | NS | NS | | | |
| 1259 | C | G | 11.7% | 222 | 0.4030 | | G | 13.2% | 1373 | 0.4720 | | Erns | NS | NK | C/g | | |
| 1282 | C | T | 0.9% | 223 | 0.0512 | | | | | | | Erns | NS | PL | | | |
| 1285 | A | G | 1.3% | 234 | 0.0686 | | | | | | | Erns | NS | EG | | | |
| 1286 | A | C | 14.1% | 234 | 0.4068 | | C | 40.9% | 1108 | 0.6927 | | Erns | NS | ED | A/c | A/c | M |
| 1289 | A | G | 0.9% | 235 | 0.0490 | | | | | | | Erns | S | | | | |
| 1293 | T | C | 1.7% | 239 | 0.0967 | | | | | | | Erns | NS | CR | | | |
| 1295 | T | C | 9.4% | 233 | 0.3126 | | C | 2.5% | 1060 | 0.1152 | | Erns | S | | | | |
| 1297 | C | T | 0.9% | 232 | 0.0496 | | | | | | | Erns | NS | TI | | | |
| 1304 | C | | | | | | T | 2.4% | 902 | 0.1147 | | Erns | S | | | | |
| 1307 | T | C | 1.9% | 214 | 0.0929 | | | | | | | Erns | S | | | | |
| 1312 | A | T | 1.3% | 228 | 0.0546 | | | | | | | Erns | NS | HL | | | |
| 1332 | C | | | | | | A | 2.2% | 558 | 0.0933 | | Erns | NS | LI | | | |
| 1338 | A | G | 9.0% | 233 | 0.3028 | | G | 2.6% | 491 | 0.1223 | | Erns | NS | TA | | | |
| 1360 | C | T | 1.3% | 239 | 0.0674 | | | | | | | Erns | NS | AV | | | |
| 1363 | G | | | | | | C | 1.0% | 291 | 0.0574 | | Erns | NS | ST | | | |
| 1365 | G | | | | | | A | 1.2% | 248 | 0.0509 | | Erns | NS | EK | | | |
| 1366 | A | G | 0.9% | 230 | 0.0499 | | | | | | | Erns | NS | EG | | | |
| 1369 | A | G | 1.3% | 228 | 0.0701 | | | | | | | Erns | NS | KR | | | |
| 1379 | C | | | | | | T | 1.4% | 504 | 0.0665 | | Erns | S | | | | |
| 1385 | T | C | 1.7% | 242 | 0.0842 | | | | | | | Erns | S | | | | |
| 1386 | T | C | 0.8% | 243 | 0.0477 | | | | | | | Erns | NS | CR | | | |
| 1388 | C | T | 0.8% | 243 | 0.0477 | | | | | | | Erns | S | | | | |

| Ref Position | Ref Base | Serum Base | Serum SNP % | Serum Depth | Serum entropy | | Culture Base | Culture SNP % | Culture Depth | Culture entropy | | Features | S/NS | Residue change | Serum PCR call | P3 PCR call | P5 PCR call |
|---|---|---|---|---|---|---|---|---|---|---|---|---|---|---|---|---|---|
| 1392 | C | T | 1.7% | 241 | 0.0845 | | | | | | | Erns | S | | | | |
| 1416 | C | T | 2.3% | 215 | 0.1105 | | | | | | | Erns | NS | HY | | | |
| 1417 | A | | | | | | G | 3.0% | 807 | 0.1338 | | Erns | NS | HR | | | |
| 1418 | T | C | 0.9% | 216 | 0.0526 | | | | | | | Erns | S | | | | |
| 1436 | C | T | 1.4% | 221 | 0.0561 | | | | | | | Erns | S | | | | |
| 1441 | T | A | 0.9% | 215 | 0.0528 | | | | | | | Erns | NS | IN | | | |
| 1443 | G | A | 0.9% | 213 | 0.0532 | | | | | | | Erns | NS | EK | | | |
| 1472 | C | T | 1.0% | 202 | 0.0555 | | | | | | | Erns | S | | | | |
| 1477 | C | T | 1.0% | 203 | 0.0553 | | | | | | | Erns | NS | AV | | | |
| 1478 | T | | | | | | C | 1.6% | 1190 | 0.0819 | | Erns | S | | | | |
| 1483 | T | C | 1.8% | 167 | 0.0900 | | | | | | | Erns | NS | LP | | | |
| 1484 | T | | | | | | C | 2.1% | 1216 | 0.1010 | | Erns | S | | | | |
| 1485 | A | T | 44.9% | 167 | 0.6880 | | T | 82.8% | 1218 | 0.4597 | | Erns | NS | TS | W | T/a | T/a |
| 1488 | G | T | 1.1% | 176 | 0.0622 | | | | | | | Erns | STOP | E* | | | |
| 1504 | G | A | 37.8% | 193 | 0.7107 | | A | 79.4% | 1415 | 0.5091 | | Erns | NS | RK | R | A/g | A/g |
| 1514 | A | G | 1.1% | 189 | 0.0587 | | | | | | | Erns | S | | | | |
| 1526 | G | | | | | | A | 2.7% | 1592 | 0.1242 | | Erns | S | | | | |
| 1536 | G | A | 8.2% | 195 | 0.2838 | | A | 1.7% | 1686 | 0.0894 | | Erns | NS | DN | | | |
| 1537 | A | G | 3.6% | 196 | 0.1613 | | | | | | | Erns | NS | DG | | | |
| 1538 | T | C | 13.2% | 197 | 0.3901 | | C | 8.7% | 1693 | 0.3090 | | Erns | S | | T/c | | |
| 1541 | T | C | 27.6% | 196 | 0.5887 | | | | | | | Erns | S | | | | |
| 1543 | A | G | 3.0% | 199 | 0.1353 | | | | | | | Erns | NS | DG | | | |
| 1545 | C | T | 3.0% | 199 | 0.1545 | | | | | | | Erns | S | | | | |
| 1558 | C | T | 1.0% | 203 | 0.0553 | | | | | | | Erns | NS | TI | | | |
| 1561 | A | | | | | | T | 1.4% | 1892 | 0.0785 | | Erns | NS | QL | | | |
| 1564 | C | A | 1.0% | 206 | 0.0547 | | | | | | | Erns | NS | AD | | | |

| Ref Position | Ref Base | Serum Base | Serum SNP % | Serum Depth | Serum entropy | | Culture Base | Culture SNP % | Culture Depth | Culture entropy | | Features | S/NS | Residue change | Serum PCR call | P3 PCR call | P5 PCR call |
|---|---|---|---|---|---|---|---|---|---|---|---|---|---|---|---|---|---|
| 1574 | T | C | 1.7% | 241 | 0.0845 | | | | | | | Erns | S | | | | |
| 1582 | C | T | 5.7% | 246 | 0.2184 | | T | 3.8% | 2301 | 0.1599 | | Erns | NS | PL | C/t | | |
| 1596 | A | G | 1.8% | 221 | 0.1031 | | | | | | | Erns | NS | KE | | | |
| 1602 | G | A | 1.9% | 209 | 0.0799 | | | | | | | Erns | NS | GR | | | |
| 1603 | G | T | 1.9% | 207 | 0.0640 | | | | | | | Erns | NS | GV | | | |
| 1604 | A | G | 1.0% | 210 | 0.0538 | | G | 5.5% | 2580 | 0.2120 | | Erns | S | | | | |
| 1606 | A | G | 1.0% | 210 | 0.0538 | | | | | | | Erns | NS | KR | | | |
| 1608 | A | T | 0.9% | 222 | 0.0514 | | | | | | | Erns | NS | NY | | | |
| 1627 | T | C | 0.9% | 225 | 0.0508 | | | | | | | Erns | NS | IT | | | |
| 1633 | C | T | 3.4% | 234 | 0.1490 | | T | 9.5% | 3423 | 0.3145 | | Erns | NS | SL | | | |
| 1636 | A | T | 1.7% | 233 | 0.0987 | | | | | | | Erns | NS | QL | | | |
| 1638 | G | A | 1.8% | 228 | 0.0883 | | | | | | | Erns | NS | GS | | | |
| 1640 | T | C | 2.6% | 228 | 0.1097 | | | | | | | Erns | S | | | | |
| 1645 | G | A | 0.9% | 230 | 0.0499 | | | | | | | Erns | NS | CY | | | |
| 1646 | C | | | | | | T | 1.1% | 3328 | 0.0614 | | Erns | S | | | | |
| 1659 | G | A | 13.7% | 234 | 0.3990 | | A | 9.8% | 3313 | 0.3205 | | Erns | NS | AT | | | |
| 1660 | C | T | 3.0% | 234 | 0.1344 | | T | 2.1% | 3313 | 0.1051 | | Erns | NS | AV | | | |
| 1661 | T | A | 0.9% | 235 | 0.0490 | | | | | | | Erns | S | | | | |
| 1663 | C | | | | | | T | 1.3% | 3298 | 0.0734 | | Erns | NS | AV | | | |
| 1664 | G | A | 7.6% | 236 | 0.2628 | | A | 8.5% | 3295 | 0.2915 | | Erns | S | | | | |
| 1691 | C | T | 1.6% | 252 | 0.0925 | | | | | | | Erns | S | | | | |
| 1693 | G | T | 0.8% | 252 | 0.0463 | | | | | | | Erns | NS | CF | | | |
| 1699 | G | A | 1.2% | 256 | 0.0638 | | | | | | | Erns | NS | SN | | | |
| 1703 | G | A | 0.8% | 256 | 0.0457 | | | | | | | Erns | S | | | | |
| 1704 | T | C | 0.8% | 256 | 0.0457 | | | | | | | Erns | NS | FL | | | |
| 1725 | C | T | 2.0% | 247 | 0.0990 | | | | | | | Erns | NS | LF | | | |

| Ref Position | Ref Base | Serum Base | Serum SNP % | Serum Depth | Serum entropy | | Culture Base | Culture SNP % | Culture Depth | Culture entropy | | Features | S/NS | Residue change | Serum PCR call | P3 PCR call | P5 PCR call |
|---|---|---|---|---|---|---|---|---|---|---|---|---|---|---|---|---|---|
| 1727 | T | C | 1.2% | 254 | 0.0642 | | | | | | | Erns | S | | | | |
| 1730 | T | C | 0.8% | 248 | 0.0469 | | | | | | | Erns | S | | | | |
| 1733 | T | C | 0.8% | 242 | 0.0479 | | | | | | | Erns | S | | | | |
| 1735 | G | A | 22.4% | 241 | 0.5320 | | A | 7.1% | 2813 | 0.2568 | | Erns | NS | GE | G/a | G/a | |
| 1737 | A | G | 13.2% | 234 | 0.4420 | | G | 29.2% | 2805 | 0.6619 | | Erns | NS | MV | | A/g | A/g |
| 1748 | C | | | | | | T | 1.8% | 2703 | 0.0938 | | Erns | S | | | | |
| 1749 | T | C | 10.0% | 221 | 0.3544 | | C | 3.2% | 2697 | 0.1461 | | Erns | S | | | | |
| 1752 | G | | | | | | T | 1.1% | 2673 | 0.0626 | | Erns | STOP | E* | | | |
| 1753 | A | | | | | | C | 1.0% | 2623 | 0.0573 | | Erns | NS | EA | | | |
| 1758 | G | A | 0.9% | 211 | 0.0536 | | | | | | | Erns | NS | AT | | | |
| 1767 | G | A | 1.0% | 210 | 0.0538 | | | | | | | Erns | NS | GR | | | |
| 1793 | A | | | | | | G | 1.5% | 2240 | 0.0801 | | Erns | S | | | | |
| 1809 | A | T | 2.7% | 148 | 0.1243 | | | | | | | Erns | NS | IL | | | |
| 1811 | A | C | 1.4% | 144 | 0.0732 | | | | | | | Erns | S | | | | |
| 1817 | A | G | 14.4% | 139 | 0.4120 | | | | | | | Erns | S | | A/g | | |
| 1822 | A | G | 1.5% | 136 | 0.0766 | | | | | | | Erns | NS | KR | | | |
| 1826 | G | A | 1.7% | 121 | 0.0842 | | | | | | | Erns | S | | | | |
| 1859 | T | | | | | | C | 1.4% | 217 | 0.0729 | | Erns | S | | | | T/c |
| 1868 | C | | | | | | T | 5.0% | 241 | 0.1979 | | E1 | S | | | | |
| 1889 | A | | | | | | T | 9.6% | 335 | 0.3653 | | E1 | S | | | A/t | A/t |
| 1900 | A | T | 1.9% | 103 | 0.0958 | | | | | | | E1 | NS | YF | | | |
| 1909 | T | | | | | | A | 9.7% | 371 | 0.3149 | | E1 | NS | FY | | T/a | |
| 1915 | A | | | | | | G | 8.9% | 383 | 0.2997 | | E1 | NS | KR | A/g | | A/g |
| 1919 | T | | | | | | G | 1.2% | 416 | 0.0566 | | E1 | NS | NK | | | |
| 1969 | A | | | | | | T | 1.7% | 586 | 0.0811 | | E1 | S | | | | |
| 1973 | C | | | | | | T | 2.6% | 615 | 0.1206 | | E1 | S | | | | |

| Ref Position | Ref Base | Serum Base | Serum SNP % | Serum Depth | Serum entropy | | Culture Base | Culture SNP % | Culture Depth | Culture entropy | | Features | S/NS | Residue change | Serum PCR call | P3 PCR call | P5 PCR call |
|---|---|---|---|---|---|---|---|---|---|---|---|---|---|---|---|---|---|
| 2044 | | | | | | | | | | | | | NS | SF | C/t | | |
| 2075 | A | | | | | | G | 5.1% | 837 | 0.2025 | | E1 | S | | | | |
| 2086 | T | | | | | | C | 7.5% | 850 | 0.2671 | | E1 | NS | VA | T/c | | |
| 2104 | A | G | 2.0% | 100 | 0.0980 | | | | | | | E1 | NS | HR | | | |
| 2109 | T | | | | | | A | 1.1% | 923 | 0.0652 | | E1 | NS | ST | | | |
| 2113 | T | C | 2.0% | 100 | 0.0980 | | | | | | | E1 | NS | IT | | | |
| 2117 | G | A | 2.9% | 103 | 0.1317 | | A | 7.1% | 954 | 0.2569 | | E1 | S | | | | |
| 2123 | C | T | 1.9% | 104 | 0.0950 | | | | | | | E1 | S | | | | |
| 2132 | T | C | 14.9% | 101 | 0.4201 | | C | 7.1% | 916 | 0.2561 | | E1 | S | | T/c | T/c | |
| 2138 | | | | | | | | | | | | E1 | NS | MI | | | G/a |
| 2144 | C | | | | | | T | 5.5% | 981 | 0.2131 | | E1 | S | | | | |
| 2171 | G | | | | | | A | 3.1% | 967 | 0.1383 | | E1 | S | | | | |
| 2180 | A | T | 1.8% | 114 | 0.0883 | | | | | | | E1 | S | | | | |
| 2181 | A | | | | | | G | 1.3% | 948 | 0.0679 | | E1 | NS | TA | | | |
| 2189 | T | | | | | | C | 2.7% | 973 | 0.1232 | | E1 | S | | | | |
| 2221 | G | A | 37.4% | 115 | 0.6610 | | A | 4.5% | 1016 | 0.1844 | | E1 | NS | RK | | | G/a |
| 2225 | T | | | | | | C | 2.4% | 1023 | 0.1148 | | E1 | S | | | | |
| 2228 | C | | | | | | T | 1.8% | 1028 | 0.0921 | | E1 | S | | | | |
| 2240 | G | A | 1.7% | 116 | 0.0871 | | | | | | | E1 | S | | | | |
| 2241 | A | G | 43.5% | 115 | 0.6846 | | G | 5.9% | 960 | 0.2252 | | E1 | NS | ND | A/g | | |
| 2252 | T | A | 1.7% | 116 | 0.0871 | | | | | | | E1 | S | | | | |
| 2258 | G | A | 1.7% | 119 | 0.0853 | | A | 2.0% | 957 | 0.0975 | | E1 | S | | | | |
| 2271 | C | | | | | | A | 3.5% | 864 | 0.1480 | | E1 | NS | LM | | | |
| 2279 | C | | | | | | T | 1.1% | 945 | 0.0599 | | E1 | S | | | | |
| 2283 | G | T | 29.1% | 117 | 0.6027 | | | | | | | E1 | STOP | E* | G/t | | |
| 2284 | A | C | 28.8% | 118 | 0.6005 | | | | | | | E1 | NS | EA | | | |

| Ref Position | Ref Base | Serum Base | Serum SNP % | Serum Depth | Serum entropy | | Culture Base | Culture SNP % | Culture Depth | Culture entropy | | Features | S/NS | Residue change | Serum PCR call | P3 PCR call | P5 PCR call |
|---|---|---|---|---|---|---|---|---|---|---|---|---|---|---|---|---|---|
| 2289 | G | A | 1.7% | 117 | 0.0865 | | A | 1.2% | 895 | 0.0624 | | E1 | NS | GS | | | |
| 2296 | T | C | 1.7% | 118 | 0.0859 | | | | | | | E1 | NS | VA | | | |
| 2312 | G | A | 8.3% | 132 | 0.2868 | | A | 7.5% | 939 | 0.2652 | | E1 | S | | | | |
| 2339 | T | C | 3.5% | 142 | 0.1524 | | C | 1.1% | 1033 | 0.0590 | | E1 | S | | | | |
| 2343 | A | G | 27.7% | 141 | 0.5897 | | | | | | | E1 | NS | ND | A/g | | |
| 2352 | A | G | 23.7% | 139 | 0.5481 | | | | | | | E2 | NS | TA | | | |
| 2353 | C | T | 1.4% | 138 | 0.0758 | | | | | | | E2 | NS | TM | | | |
| 2366 | T | C | 1.4% | 146 | 0.0724 | | | | | | | E2 | S | | | | |
| 2369 | A | G | 1.4% | 140 | 0.0749 | | | | | | | E2 | S | | | | |
| 2378 | C | | | | | | G | 1.2% | 1004 | 0.0613 | | E2 | S | | | | |
| 2381 | T | | | | | | C | 5.1% | 1005 | 0.2007 | | E2 | S | | | | |
| 2384 | G | | | | | | A | 1.3% | 996 | 0.0692 | | E2 | S | | | | |
| 2396 | C | T | 4.0% | 126 | 0.1669 | | | | | | | E2 | S | | | | |
| 2421 | C | | | | | | T | 2.9% | 1126 | 0.1323 | | E2 | S | | | | |
| 2429 | G | A | 12.6% | 167 | 0.3782 | | A | 38.9% | 1158 | 0.6686 | | E2 | S | | | R | R |
| 2441 | G | | | | | | T | 1.1% | 1126 | 0.0590 | | E2 | S | | | | |
| 2467 | A | G | 22.7% | 163 | 0.5356 | | G | 10.1% | 1174 | 0.3281 | | E2 | NS | EG | A/g | A/g | |
| 2470 | A | | | | | | T | 5.0% | 1248 | 0.2047 | | E2 | NS | YF | | | |
| 2486 | C | A | 1.1% | 181 | 0.0608 | | | | | | | E2 | S | | | | |
| 2494 | A | T | 1.1% | 176 | 0.0622 | | | | | | | E2 | NS | DV | | | |
| 2495 | T | | | | | | C | 2.6% | 1361 | 0.1222 | | E2 | S | | | | |
| 2504 | C | | | | | | T | 2.0% | 1402 | 0.0958 | | E2 | S | | | | |
| 2507 | A | | | | | | G | 1.2% | 1405 | 0.0605 | | E2 | S | | | | |
| 2525 | A | | | | | | G | 7.7% | 1606 | 0.2709 | | E2 | S | | | | |
| 2532 | G | A | 45.9% | 233 | 0.7325 | | A | 79.4% | 1643 | 0.5082 | | E2 | NS | VI | R | A/g | A/g |
| 2533 | T | C | 5.6% | 232 | 0.2159 | | C | 3.7% | 1649 | 0.1583 | | E2 | NS | VA | | | |

| Ref Position | Ref Base | Serum Base | Serum SNP % | Serum Depth | Serum entropy | | Culture Base | Culture SNP % | Culture Depth | Culture entropy | | Features | S/NS | Residue change | Serum PCR call | P3 PCR call | P5 PCR call |
|---|---|---|---|---|---|---|---|---|---|---|---|---|---|---|---|---|---|
| 2538 | A | G | 1.7% | 229 | 0.0880 | | | | | | | E2 | NS | KE | | | |
| 2551 | G | A | 0.9% | 223 | 0.0512 | | | | | | | E2 | NS | SN | | | |
| 2554 | G | A | 0.9% | 223 | 0.0512 | | | | | | | E2 | NS | GE | | | |
| 2560 | T | C | 22.9% | 231 | 0.5365 | | C | 8.5% | 1759 | 0.2906 | | E2 | NS | VA | T/c | | |
| 2564 | G | | | | | | A | 2.2% | 1946 | 0.1061 | | E2 | S | | | | |
| 2581 | T | G | 1.6% | 246 | 0.0831 | | | | | | | E2 | NS | IR | | | |
| 2587 | G | A | 0.8% | 240 | 0.0482 | | | | | | | E2 | STOP | W* | | | |
| 2609 | A | | | | | | G | 2.0% | 2066 | 0.0993 | | E2 | S | | | | |
| 2628 | A | G | 9.5% | 241 | 0.3149 | | | | | | | E2 | NS | RG | | | |
| 2666 | C | T | 0.9% | 234 | 0.0492 | | T | 1.6% | 1994 | 0.0807 | | E2 | S | | | | |
| 2668 | T | C | 1.3% | 233 | 0.0688 | | | | | | | E2 | NS | LP | | | |
| 2712 | C | A | 27.0% | 222 | 0.5835 | | A | 75.9% | 1856 | 0.5675 | | E2 | NS | QK | C/a | A/c | A/c |
| 2729 | A | G | 4.5% | 224 | 0.1824 | | G | 4.9% | 1877 | 0.2058 | | E2 | S | | | | |
| 2745 | G | A | 0.9% | 228 | 0.0503 | | | | | | | E2 | NS | EK | | | |
| 2748 | T | C | 1.3% | 224 | 0.0711 | | | | | | | E2 | NS | FL | | | |
| 2755 | T | A | 1.4% | 217 | 0.0729 | | | | | | | E2 | NS | LH | | | |
| 2773 | A | G | 1.0% | 208 | 0.0542 | | | | | | | E2 | NS | KR | | | |
| 2776 | C | A | 1.0% | 197 | 0.0567 | | | | | | | E2 | NS | PH | | | |
| 2783 | G | A | 1.1% | 187 | 0.0592 | | | | | | | E2 | S | | | | |
| 2798 | C | | | | | | A | 1.4% | 1614 | 0.0747 | | E2 | NS | NK | | | |
| 2807 | G | | | | | | A | 1.8% | 1651 | 0.0934 | | E2 | S | | | | |
| 2816 | A | T | 1.1% | 189 | 0.0587 | | | | | | | E2 | S | | | | |
| 2817 | C | T | 20.1% | 189 | 0.5019 | | | | | | | E2 | NS | PS | C/t | | |
| 2820 | G | T | 12.8% | 187 | 0.3832 | | | | | | | E2 | NS | AS | G/t | | |
| 2821 | C | T | 1.1% | 186 | 0.0594 | | | | | | | E2 | S | | | | |
| 2856 | T | C | 0.8% | 236 | 0.0489 | | | | | | | E2 | NS | SP | | | |

| Ref Position | Ref Base | Serum Base | Serum SNP % | Serum Depth | Serum entropy | | Culture Base | Culture SNP % | Culture Depth | Culture entropy | | Features | S/NS | Residue change | Serum PCR call | P3 PCR call | P5 PCR call |
|---|---|---|---|---|---|---|---|---|---|---|---|---|---|---|---|---|---|
| 2871 | T | | | | | | G | 1.4% | 1969 | 0.0724 | | E2 | NS | LV | | | |
| 2873 | | | | | | | | | | | | | S | | | | A/g |
| 2877 | A | G | 0.9% | 223 | 0.0512 | | | | | | | E2 | NS | ND | | | |
| 2881 | G | A | 1.3% | 224 | 0.0711 | | | | | | | E2 | NS | RK | | | |
| 2898 | G | A | 1.4% | 213 | 0.0740 | | | | | | | E2 | NS | AT | | | |
| 2899 | C | T | 4.7% | 215 | 0.1881 | | | | | | | E2 | NS | AV | | | |
| 2900 | A | G | 1.9% | 215 | 0.0926 | | | | | | | E2 | S | | | | |
| 2904 | A | G | 21.3% | 225 | 0.5183 | | G | 9.4% | 1963 | 0.3109 | | E2 | NS | IV | A/g | | |
| 2912 | G | A | 0.9% | 234 | 0.0492 | | | | | | | E2 | S | | | | |
| 2922 | T | A | 13.3% | 240 | 0.4677 | | A | 32.5% | 2027 | 0.6303 | | E2 | NS | ST | T/a | T/a | W |
| 2930 | A | G | 1.3% | 234 | 0.0686 | | | | | | | E2 | S | | | | |
| 2936 | C | T | 9.7% | 227 | 0.3123 | | T | 4.3% | 1941 | 0.1852 | | E2 | S | | | | |
| 2954 | C | A | 8.3% | 205 | 0.2859 | | A | 32.6% | 1778 | 0.6314 | | E2 | S | | C/a | C/a | M |
| 2958 | C | T | 1.9% | 210 | 0.0943 | | | | | | | E2 | STOP | Q* | | | |
| 2960 | A | G | 8.1% | 210 | 0.2811 | | G | 6.9% | 1790 | 0.2494 | | E2 | S | | | | |
| 2962 | A | T | 0.9% | 211 | 0.0536 | | | | | | | E2 | NS | KM | | | |
| 2966 | T | C | 2.8% | 211 | 0.1167 | | | | | | | E2 | S | | | | |
| 2981 | C | A | 1.1% | 189 | 0.0587 | | | | | | | E2 | S | | | | |
| 2984 | C | | | | | | T | 3.1% | 1707 | 0.1384 | | E2 | S | | | | |
| 2985 | G | A | 1.0% | 192 | 0.0579 | | | | | | | E2 | NS | DN | | | |
| 2987 | C | A | 2.1% | 194 | 0.1004 | | | | | | | E2 | S | | | | |
| 2998 | G | A | 1.0% | 199 | 0.0562 | | | | | | | E2 | NS | GE | | | |
| 3017 | G | A | 1.0% | 203 | 0.0553 | | | | | | | E2 | S | | | | |
| 3048 | T | C | 1.4% | 210 | 0.0749 | | | | | | | E2 | NS | SP | | | |
| 3049 | C | T | 0.9% | 211 | 0.0536 | | | | | | | E2 | NS | SF | | | |
| 3083 | T | | | | | | C | 1.8% | 1138 | 0.0920 | | E2 | S | | | | |

| Ref Position | Ref Base | Serum Base | Serum SNP % | Serum Depth | Serum entropy | | Culture Base | Culture SNP % | Culture Depth | Culture entropy | | Features | S/NS | Residue change | Serum PCR call | P3 PCR call | P5 PCR call |
|---|---|---|---|---|---|---|---|---|---|---|---|---|---|---|---|---|---|
| 3084 | C | A | 1.0% | 206 | 0.0547 | | | | | | | E2 | NS | QK | | | |
| 3091 | G | A | 7.8% | 206 | 0.2730 | | A | 2.6% | 926 | 0.1174 | | E2 | NS | SN | | | |
| 3092 | T | A | 8.1% | 209 | 0.2747 | | A | 1.9% | 924 | 0.0928 | | E2 | NS | SR | | | |
| 3093 | G | A | 1.0% | 199 | 0.0562 | | | | | | | E2 | NS | EK | | | |
| 3101 | A | | | | | | G | 1.5% | 1019 | 0.0735 | | E2 | S | | | | |
| 3106 | A | | | | | | T | 1.6% | 1176 | 0.0881 | | E2 | NS | HL | | | |
| 3117 | G | A | 1.2% | 257 | 0.0494 | | | | | | | E2 | NS | GS | | | |
| 3122 | G | A | 16.9% | 260 | 0.4547 | | A | 32.5% | 1534 | 0.6595 | | E2 | S | | G/a | R | R |
| 3132 | G | A | 0.8% | 257 | 0.0455 | | | | | | | E2 | NS | EK | | | |
| 3133 | A | T | 1.7% | 238 | 0.0853 | | | | | | | E2 | NS | EV | | | |
| 3136 | A | G | 1.3% | 238 | 0.0677 | | | | | | | E2 | NS | NS | | | |
| 3137 | T | A | 0.8% | 238 | 0.0485 | | | | | | | E2 | NS | NK | | | |
| 3146 | C | T | 0.8% | 258 | 0.0454 | | | | | | | E2 | S | | | | |
| 3151 | G | A | 0.8% | 257 | 0.0455 | | | | | | | E2 | NS | RK | | | |
| 3162 | G | A | 22.8% | 254 | 0.5373 | | A | 4.4% | 3833 | 0.1844 | | E2 | NS | DN | G/a | | G/a |
| 3174 | A | G | 18.0% | 261 | 0.4715 | | G | 6.6% | 3882 | 0.2488 | | E2 | NS | ND | A/g | | |
| 3177 | A | G | 0.8% | 261 | 0.0450 | | | | | | | E2 | NS | RG | | | |
| 3182 | A | T | 2.3% | 261 | 0.1095 | | T | 9.4% | 3918 | 0.3171 | | E2 | NS | ED | | A/t | |
| 3184 | G | A | 0.8% | 261 | 0.0450 | | | | | | | E2 | NS | GD | | | |
| 3186 | G | A | 0.8% | 261 | 0.0450 | | | | | | | E2 | NS | VM | | | |
| 3194 | A | G | 1.5% | 260 | 0.0795 | | | | | | | E2 | NS | IM | | | |
| 3197 | A | | | | | | G | 1.6% | 3914 | 0.0834 | | E2 | S | | | | |
| 3200 | G | A | 0.8% | 263 | 0.0447 | | | | | | | E2 | S | | | | |
| 3201 | C | T | 0.8% | 262 | 0.0448 | | | | | | | E2 | NS | HY | | | |
| 3203 | C | A | 18.7% | 262 | 0.4819 | | | | | | | E2 | NS | HQ | C/a | | |
| 3207 | C | A | 17.2% | 262 | 0.4587 | | A | 6.4% | 3901 | 0.2397 | | E2 | NS | LM | C/a | | |

| Ref Position | Ref Base | Serum Base | Serum SNP % | Serum Depth | Serum entropy | | Culture Base | Culture SNP % | Culture Depth | Culture entropy | | Features | S/NS | Residue change | Serum PCR call | P3 PCR call | P5 PCR call |
|---|---|---|---|---|---|---|---|---|---|---|---|---|---|---|---|---|---|
| 3212 | A | T | 0.8% | 265 | 0.0444 | | | | | | | E2 | S | | | | |
| 3227 | A | C | 2.3% | 256 | 0.1260 | | | | | | | E2 | S | | | | |
| 3232 | C | A | 1.2% | 253 | 0.0644 | | | | | | | E2 | NS | TK | | | |
| 3247 | C | T | 22.9% | 236 | 0.5378 | | T | 4.4% | 3878 | 0.1828 | | E2 | NS | TI | C/t | | C/t |
| 3260 | T | A | 1.4% | 218 | 0.0567 | | | | | | | E2 | S | | | | |
| 3262 | A | G | 0.9% | 219 | 0.0520 | | | | | | | E2 | NS | KR | | | |
| 3278 | T | A | 0.9% | 215 | 0.0528 | | | | | | | E2 | S | | | | |
| 3284 | G | A | 1.4% | 207 | 0.0758 | | | | | | | E2 | S | | | | |
| 3287 | A | | | | | | G | 1.4% | 3892 | 0.0731 | | E2 | S | | | | |
| 3288 | T | C | 16.3% | 203 | 0.4439 | | C | 2.7% | 3891 | 0.1246 | | E2 | NS | YH | T/c | | T/c |
| 3298 | T | C | 3.1% | 191 | 0.1396 | | | | | | | E2 | NS | NS | IT | | T/c |
| 3315 | G | A | 1.2% | 171 | 0.0637 | | A | 3.3% | 3714 | 0.1483 | | E2 | NS | VI | | | |
| 3326 | G | T | 1.2% | 166 | 0.0652 | | | | | | | E2 | S | | | | |
| 3336 | T | G | 3.1% | 161 | 0.1384 | | G | 1.0% | 3588 | 0.0564 | | E2 | NS | FV | | | |
| 3337 | T | A | 2.5% | 161 | 0.1163 | | | | | | | E2 | NS | FY | | | |
| 3341 | C | T | 1.9% | 107 | 0.0929 | | | | | | | E2 | S | | | | |
| 3344 | C | | | | | | T | 1.0% | 2046 | 0.0609 | | E2 | S | | | | |
| 3354 | T | | | | | | G | 1.0% | 966 | 0.0628 | | E2 | NS | LV | | | |
| 3355 | T | | | | | | C | 1.5% | 940 | 0.0815 | | E2 | NS | LS | | | |
| 3356 | G | | | | | | A | 1.0% | 884 | 0.0578 | | E2 | S | | | | |
| 3366 | T | | | | | | A | 13.5% | 585 | 0.4322 | | E2 | NS | YN | | | |
| 3368 | T | | | | | | A | 1.9% | 520 | 0.0893 | | E2 | STOP | Y* | | | |
| 3401 | C | | | | | | T | 3.5% | 546 | 0.1510 | | E2 | S | | | | |
| 3411 | G | | | | | | A | 1.3% | 541 | 0.0628 | | E2 | NS | GR | | | |
| 3419 | T | | | | | | C | 3.1% | 554 | 0.1371 | | E2 | S | | | | |
| 3420 | C | | | | | | T | 1.3% | 557 | 0.0750 | | E2 | STOP | Q* | | | |

| Ref Position | Ref Base | Serum Base | Serum SNP % | Serum Depth | Serum entropy | | Culture Base | Culture SNP % | Culture Depth | Culture entropy | | Features | S/NS | Residue change | Serum PCR call | P3 PCR call | P5 PCR call |
|---|---|---|---|---|---|---|---|---|---|---|---|---|---|---|---|---|---|
| 3442 | C | | | | | | T | 1.9% | 531 | 0.0935 | | E2 | NS | AV | | | |
| 3468 | G | | | | | | T | 2.3% | 570 | 0.1088 | | E2 | NS | AS | | | |
| 3473 | | | | | | | | | | | | E2 | NS | ED | G/t | | |
| 3477 | A | | | | | | G | 1.4% | 552 | 0.0758 | | E2 | NS | IV | | | |
| 3479 | A | | | | | | T | 1.8% | 554 | 0.0904 | | E2 | S | | | | |
| 3488 | A | | | | | | G | 1.1% | 564 | 0.0589 | | E2 | NS | IM | | | |
| 3491 | | | | | | | | | | | | E2 | S | | G/a | | |
| 3565 | C | | | | | | T | 3.6% | 667 | 0.1550 | | E2 | NS | SL | | | |
| 3622 | A | | | | | | G | 8.2% | 643 | 0.2823 | | p7 | NS | DG | A/g | | |
| 3648 | T | | | | | | C | 9.2% | 638 | 0.3082 | | p7 | S | | | | |
| 3667 | T | | | | | | A | 1.3% | 706 | 0.0584 | | p7 | NS | LQ | | | |
| 3689 | G | | | | | | A | 1.6% | 749 | 0.0821 | | p7 | S | | | | |
| 3698 | | | | | | | | | | | | p7 | S | | G/a | | |
| 3714 | | | | | | | | | | | | p7 | S | | C/t | | |
| 3716 | A | | | | | | G | 6.3% | 827 | 0.2348 | | p7 | S | | | A/g | A/g |
| 3753 | T | | | | | | C | 1.3% | 851 | 0.0752 | | p7 | S | | | | |
| 3766 | A | G | 1.9% | 107 | 0.0929 | | | | | | | p7 | NS | DG | | | |
| 3768 | G | A | 5.5% | 109 | 0.2131 | | | | | | | p7 | NS | VM | | | |
| 3769 | T | | | | | | C | 2.5% | 831 | 0.1146 | | p7 | NS | VA | | | |
| 3775 | A | | | | | | G | 1.2% | 892 | 0.0665 | | p7 | NS | KR | | | |
| 3787 | G | T | 1.6% | 122 | 0.0836 | | | | | | | NS2 | NS | GV | | | |
| 3790 | A | G | 22.4% | 125 | 0.5278 | | | | | | | NS2 | NS | DG | A/g | | |
| 3795 | G | T | 1.6% | 125 | 0.0820 | | | | | | | NS2 | NS | GW | | | |
| 3801 | T | | | | | | C | 1.8% | 1023 | 0.0947 | | NS2 | S | | | | |
| 3802 | T | C | 3.1% | 127 | 0.1399 | | | | | | | NS2 | NS | LS | | | |
| 3803 | G | | | | | | A | 2.3% | 1028 | 0.1081 | | NS2 | S | | | | |

| Ref Position | Ref Base | Serum Base | Serum SNP % | Serum Depth | Serum entropy | | Culture Base | Culture SNP % | Culture Depth | Culture entropy | | Features | S/NS | Residue change | Serum PCR call | P3 PCR call | P5 PCR call |
|---|---|---|---|---|---|---|---|---|---|---|---|---|---|---|---|---|---|
| 3812 | A | | | | | | G | 3.6% | 1034 | 0.1520 | | NS2 | NS | IM | | | |
| 3831 | G | A | 2.6% | 156 | 0.1192 | | | | | | | NS2 | NS | VI | | | |
| 3833 | T | A | 1.3% | 151 | 0.0704 | | | | | | | NS2 | S | | | | |
| 3842 | C | A | 1.3% | 160 | 0.0672 | | | | | | | NS2 | S | | | | |
| 3843 | G | T | 1.3% | 160 | 0.0672 | | | | | | | NS2 | NS | VF | | | |
| 3857 | C | T | 1.2% | 164 | 0.0659 | | | | | | | NS2 | S | | | | |
| 3860 | A | | | | | | G | 1.3% | 1240 | 0.0770 | | NS2 | NS | IM | | | |
| 3882 | G | A | 1.1% | 177 | 0.0619 | | A | 1.8% | 1305 | 0.0894 | | NS2 | NS | VI | | | |
| 3891 | G | A | 8.1% | 173 | 0.2810 | | A | 6.6% | 1287 | 0.2506 | | NS2 | NS | VI | | | |
| 3902 | A | G | 1.2% | 167 | 0.0649 | | | | | | | NS2 | S | | | | |
| 3934 | C | | | | | | T | 1.4% | 1234 | 0.0700 | | NS2 | NS | AV | | | |
| 3936 | | | | | | | | | | | | NS2 | NS | PS | | | C/t |
| 3959 | A | | | | | | G | 2.9% | 1257 | 0.1300 | | NS2 | S | | | | |
| 3967 | C | G | 1.3% | 159 | 0.0675 | | | | | | | NS2 | NS | TS | | | |
| 3968 | C | A | 1.2% | 162 | 0.0665 | | | | | | | NS2 | S | | | | |
| 4004 | C | T | 1.4% | 143 | 0.0736 | | | | | | | NS2 | S | | | | |
| 4048 | C | T | 1.5% | 132 | 0.0785 | | | | | | | NS2 | NS | SL | | | |
| 4066 | G | A | 7.0% | 128 | 0.2411 | | A | 28.9% | 1067 | 0.6123 | | NS2 | S | | G/a | R | G/a |
| 4075 | C | T | 6.2% | 129 | 0.2325 | | T | 2.6% | 1069 | 0.1212 | | NS2 | S | | | | C/t |
| 4083 | G | | | | | | A | 1.4% | 1074 | 0.0735 | | NS2 | NS | GS | | | |
| 4100 | A | G | 1.8% | 113 | 0.0889 | | | | | | | NS2 | S | | | | |
| 4104 | G | C | 1.6% | 126 | 0.0815 | | | | | | | NS2 | NS | VL | | | |
| 4112 | C | T | 1.5% | 133 | 0.0780 | | | | | | | NS2 | S | | | | |
| 4134 | C | T | 1.6% | 125 | 0.0820 | | | | | | | NS2 | S | | | | |
| 4136 | A | G | 1.6% | 125 | 0.0820 | | | | | | | NS2 | S | | | | |
| 4143 | C | T | 1.7% | 117 | 0.0865 | | | | | | | NS2 | S | | | | |

| Ref Position | Ref Base | Serum Base | Serum SNP % | Serum Depth | Serum entropy | | Culture Base | Culture SNP % | Culture Depth | Culture entropy | | Features | S/NS | Residue change | Serum PCR call | P3 PCR call | P5 PCR call |
|---|---|---|---|---|---|---|---|---|---|---|---|---|---|---|---|---|---|
| 4147 | A | T | 1.6% | 127 | 0.0810 | | | | | | | NS2 | NS | YF | | | |
| 4151 | G | T | 1.6% | 123 | 0.0831 | | | | | | | NS2 | S | | | | |
| 4161 | A | G | 1.5% | 136 | 0.0766 | | | | | | | NS2 | NS | TA | | | |
| 4164 | G | | | | | | A | 1.6% | 1141 | 0.0811 | | NS2 | NS | VI | | | |
| 4174 | T | G | 3.8% | 133 | 0.1602 | | G | 3.3% | 1152 | 0.1429 | | NS2 | NS | MR | | | |
| 4195 | G | A | 1.4% | 143 | 0.0736 | | | | | | | NS2 | NS | GD | | | |
| 4202 | G | A | 1.4% | 142 | 0.0740 | | | | | | | NS2 | S | | | | |
| 4203 | T | | | | | | C | 5.9% | 1251 | 0.2232 | | NS2 | S | | | | |
| 4212 | A | G | 25.0% | 144 | 0.5623 | | G | 39.0% | 1248 | 0.6689 | | NS2 | NS | MV | A/g | R | G/a |
| 4217 | A | | | | | | G | 1.3% | 1264 | 0.0713 | | NS2 | S | | | | |
| 4224 | C | T | 1.4% | 144 | 0.0732 | | | | | | | NS2 | S | | | | |
| 4226 | G | T | 1.4% | 144 | 0.0732 | | T | 3.5% | 1273 | 0.1503 | | NS2 | S | | | | |
| 4234 | T | C | 28.4% | 141 | 0.5964 | | C | 1.3% | 1261 | 0.0714 | | NS2 | NS | IT | Y | | |
| 4241 | A | G | 1.4% | 139 | 0.0753 | | | | | | | NS2 | S | | | | |
| 4247 | C | A | 2.8% | 141 | 0.1290 | | | | | | | NS2 | S | | | | |
| 4256 | A | G | 2.8% | 144 | 0.1269 | | G | 2.5% | 1194 | 0.1174 | | NS2 | S | | | | |
| 4263 | A | G | 21.5% | 144 | 0.5209 | | G | 35.2% | 1169 | 0.6489 | | NS2 | NS | IV | A/g | R | R |
| 4275 | C | T | 2.9% | 140 | 0.1297 | | | | | | | NS2 | NS | PS | | | |
| 4280 | C | T | 1.4% | 138 | 0.0758 | | | | | | | NS2 | S | | | | |
| 4283 | T | C | 1.4% | 142 | 0.0740 | | C | 4.2% | 1131 | 0.1756 | | NS2 | S | | | | T/c |
| 4298 | A | G | 4.0% | 149 | 0.1534 | | | | | | | NS2 | S | | | | |
| 4305 | C | T | 2.7% | 149 | 0.1236 | | T | 8.9% | 1113 | 0.3089 | | NS2 | S | | | C/t | C/t |
| 4314 | A | G | 1.3% | 149 | 0.0712 | | G | 4.9% | 1107 | 0.1949 | | NS2 | NS | IV | | | |
| 4326 | A | | | | | | G | 1.6% | 1162 | 0.0835 | | NS2 | NS | IV | | | |
| 4359 | A | C | 3.7% | 164 | 0.1802 | | | | | | | NS2 | NS | KQ | | | |
| 4367 | T | C | 1.2% | 171 | 0.0637 | | | | | | | NS2 | S | | | | |

| Ref Position | Ref Base | Serum Base | Serum SNP % | Serum Depth | Serum entropy | | Culture Base | Culture SNP % | Culture Depth | Culture entropy | | Features | S/NS | Residue change | Serum PCR call | P3 PCR call | P5 PCR call |
|---|---|---|---|---|---|---|---|---|---|---|---|---|---|---|---|---|---|
| 4406 | A | T | 1.0% | 194 | 0.0574 | | | | | | | NS2 | S | | | | |
| 4412 | T | | | | | | A | 1.9% | 1289 | 0.0903 | | NS2 | S | | | | |
| 4436 | G | A | 2.2% | 185 | 0.1043 | | | | | | | NS2 | S | | | | |
| 4450 | T | A | 1.1% | 189 | 0.0587 | | | | | | | NS2 | NS | IK | | | |
| 4466 | C | G | 1.1% | 190 | 0.0584 | | | | | | | NS2 | S | | | | |
| 4472 | G | | | | | | A | 1.0% | 1643 | 0.0576 | | NS2 | S | | | | |
| 4478 | G | | | | | | A | 1.1% | 1673 | 0.0621 | | NS2 | S | | | | |
| 4487 | T | C | 1.0% | 195 | 0.0572 | | | | | | | NS2 | S | | | | |
| 4506 | T | C | 17.5% | 189 | 0.4631 | | C | 30.1% | 1822 | 0.6115 | | NS2 | S | | | T/c | T/c |
| 4508 | A | G | 1.1% | 189 | 0.0587 | | | | | | | NS2 | S | | | | |
| 4541 | A | G | 2.3% | 172 | 0.1105 | | | | | | | NS2 | NS | IM | | | |
| 4556 | G | | | | | | A | 1.6% | 1702 | 0.0815 | | NS2 | S | | | | |
| 4562 | T | C | 2.5% | 157 | 0.1187 | | | | | | | NS2 | S | | | | |
| 4565 | A | | | | | | C | 2.3% | 1707 | 0.1089 | | NS2 | S | | | | |
| 4572 | A | G | 13.3% | 150 | 0.3927 | | G | 41.2% | 1691 | 0.6776 | | NS2 | NS | IV | | A/g | R |
| 4574 | C | T | 29.5% | 149 | 0.6068 | | | | | | | NS2 | S | | | | |
| 4591 | T | A | 1.4% | 146 | 0.0724 | | | | | | | NS2 | NS | MK | | | |
| 4604 | A | G | 7.4% | 135 | 0.2641 | | G | 24.8% | 1656 | 0.5733 | | NS2 | S | | | R | |
| 4610 | T | C | 1.6% | 127 | 0.0810 | | | | | | | NS2 | S | | | | |
| 4622 | G | T | 3.4% | 116 | 0.1739 | | | | | | | NS2 | NS | ED | | | |
| 4641 | G | | | | | | A | 1.9% | 996 | 0.1042 | | NS2 | NS | EK | | | |
| 4647 | G | | | | | | A | 1.6% | 898 | 0.0897 | | NS2 | NS | EK | | | |
| 4700 | A | | | | | | G | 5.0% | 968 | 0.1973 | | NS2 | S | | | A/g | |
| 4701 | A | G | 1.9% | 108 | 0.0922 | | G | 2.2% | 969 | 0.1015 | | NS2 | NS | IV | | | |
| 4706 | A | G | 3.8% | 106 | 0.1607 | | | | | | | NS2 | NS | IM | | | |
| 4715 | A | | | | | | G | 1.3% | 985 | 0.0702 | | NS2 | S | | | | |

| Ref Position | Ref Base | Serum Base | Serum SNP % | Serum Depth | Serum entropy | | Culture Base | Culture SNP % | Culture Depth | Culture entropy | | Features | S/NS | Residue change | Serum PCR call | P3 PCR call | P5 PCR call |
|---|---|---|---|---|---|---|---|---|---|---|---|---|---|---|---|---|---|
| 4739 | G | | | | | | A | 2.7% | 1205 | 0.1226 | | NS2 | S | | | | |
| 4750 | G | A | 3.2% | 124 | 0.1425 | | | | | | | NS2 | NS | GE | | | |
| 4763 | C | | | | | | T | 5.3% | 1165 | 0.2062 | | NS2 | S | | | | |
| 4766 | C | T | 3.0% | 135 | 0.1335 | | | | | | | NS2 | S | | | | |
| 4772 | G | A | 1.5% | 135 | 0.0771 | | | | | | | NS2 | S | | | | |
| 4776 | A | | | | | | G | 1.3% | 1199 | 0.0709 | | NS2 | NS | IV | | | |
| 4778 | C | A | 1.5% | 133 | 0.0780 | | | | | | | NS2 | S | | | | |
| 4806 | A | T | 1.4% | 142 | 0.0740 | | | | | | | NS2 | NS | NY | | | |
| 4820 | C | T | 1.3% | 149 | 0.0712 | | | | | | | NS2 | S | | | | |
| 4827 | A | G | 1.3% | 152 | 0.0701 | | | | | | | NS2 | NS | IV | | | |
| 4874 | A | | | | | | T | 2.9% | 1566 | 0.1303 | | NS2 | S | | | | |
| 4880 | T | C | 2.5% | 162 | 0.1329 | | | | | | | NS2 | S | | | | |
| 4889 | T | A | 1.8% | 163 | 0.0722 | | | | | | | NS2 | NS | HQ | | | |
| 4902 | A | T | 1.2% | 166 | 0.0652 | | | | | | | NS2 | NS | TS | | | |
| 4919 | A | G | 1.9% | 159 | 0.0936 | | | | | | | NS2 | S | | | | |
| 4936 | G | | | | | | A | 1.1% | 1321 | 0.0687 | | NS2 | NS | RK | | | |
| 4967 | C | T | 1.4% | 143 | 0.0736 | | | | | | | NS2 | S | | | | |
| 4978 | G | A | 1.5% | 135 | 0.0771 | | | | | | | NS2 | NS | GD | | | |
| 4995 | T | | | | | | A | 1.1% | 1128 | 0.0637 | | NS2 | NS | YN | | | |
| 5000 | T | C | 1.5% | 132 | 0.0785 | | | | | | | NS2 | S | | | | |
| 5015 | T | C | 1.6% | 126 | 0.0815 | | C | 3.7% | 1116 | 0.1574 | | NS2 | S | | | | |
| 5018 | C | T | 1.6% | 127 | 0.0810 | | | | | | | NS2 | S | | | | |
| 5056 | T | C | 1.6% | 124 | 0.0826 | | | | | | | NS2 | NS | LP | | | |
| 5072 | A | | | | | | G | 3.9% | 767 | 0.1651 | | NS2 | S | | | | |
| 5076 | | | | | | | | | | | | NS2 | NS | LV | | | C/g |
| 5177 | G | | | | | | A | 1.3% | 544 | 0.0688 | | NS3 | S | | | | |

| Ref Position | Ref Base | Serum Base | Serum SNP % | Serum Depth | Serum entropy | | Culture Base | Culture SNP % | Culture Depth | Culture entropy | | Features | S/NS | Residue change | Serum PCR call | P3 PCR call | P5 PCR call |
|---|---|---|---|---|---|---|---|---|---|---|---|---|---|---|---|---|---|
| 5237 | T | | | | | | C | 2.8% | 499 | 0.1279 | | NS3 | S | | | | |
| 5267 | G | | | | | | A | 1.0% | 579 | 0.0577 | | NS3 | S | | | | |
| 5275 | T | | | | | | A | 1.9% | 589 | 0.0928 | | NS3 | NS | LQ | | | |
| 5304 | T | | | | | | G | 1.2% | 598 | 0.0637 | | NS3 | NS | WG | | | |
| 5319 | C | | | | | | A | 1.8% | 610 | 0.0903 | | NS3 | NS | QK | | | |
| 5327 | G | | | | | | A | 2.5% | 610 | 0.1109 | | NS3 | S | | | | |
| 5339 | T | | | | | | C | 2.8% | 605 | 0.1337 | | NS3 | S | | | | |
| 5343 | C | | | | | | A | 1.4% | 584 | 0.0666 | | NS3 | NS | HN | | | |
| 5347 | T | | | | | | A | 2.5% | 590 | 0.1284 | | NS3 | NS | VE | | | |
| 5375 | C | | | | | | T | 1.6% | 964 | 0.0802 | | NS3 | S | | | | |
| 5432 | G | | | | | | A | 1.6% | 1144 | 0.0809 | | NS3 | S | | | | |
| 5459 | A | | | | | | C | 1.6% | 1122 | 0.0947 | | NS3 | S | | | | |
| 5477 | C | T | 1.9% | 105 | 0.0943 | | | | | | | NS3 | S | | | | |
| 5499 | G | T | 1.9% | 103 | 0.0958 | | | | | | | NS3 | STOP | E* | | | |
| 5504 | A | T | 1.9% | 103 | 0.0958 | | | | | | | NS3 | S | | | | |
| 5519 | G | | | | | | A | 3.3% | 1138 | 0.1480 | | NS3 | S | | | | |
| 5522 | C | | | | | | A | 3.0% | 1160 | 0.1310 | | NS3 | S | | | | |
| 5525 | A | G | 1.9% | 104 | 0.0950 | | | | | | | NS3 | S | | | | |
| 5531 | A | G | 1.9% | 106 | 0.0936 | | | | | | | NS3 | S | | | | |
| 5539 | A | G | 2.0% | 100 | 0.0980 | | | | | | | NS3 | NS | HR | | | |
| 5561 | A | G | 1.8% | 112 | 0.0896 | | | | | | | NS3 | S | | | | |
| 5567 | G | A | 1.8% | 114 | 0.0883 | | | | | | | NS3 | S | | | | |
| 5585 | C | A | 1.8% | 111 | 0.0902 | | | | | | | NS3 | S | | | | |
| 5588 | A | T | 1.6% | 122 | 0.0836 | | | | | | | NS3 | S | | | | |
| 5591 | G | C | 1.6% | 123 | 0.0831 | | | | | | | NS3 | S | | | | |
| 5633 | G | | | | | | A | 4.8% | 1178 | 0.1912 | | NS3 | S | | | | |

| Ref Position | Ref Base | Serum Base | Serum SNP % | Serum Depth | Serum entropy | | Culture Base | Culture SNP % | Culture Depth | Culture entropy | | Features | S/NS | Residue change | Serum PCR call | P3 PCR call | P5 PCR call |
|---|---|---|---|---|---|---|---|---|---|---|---|---|---|---|---|---|---|
| 5654 | C | | | | | | T | 1.8% | 1246 | 0.0942 | | NS3 | S | | | | |
| 5655 | G | T | 1.6% | 126 | 0.0815 | | | | | | | NS3 | STOP | G* | | | |
| 5684 | G | A | 1.5% | 134 | 0.0776 | | | | | | | NS3 | S | | | | |
| 5705 | C | T | 1.5% | 130 | 0.0795 | | | | | | | NS3 | S | | | | |
| 5720 | C | T | 3.4% | 118 | 0.1480 | | T | 2.8% | 1512 | 0.1269 | | NS3 | S | | | | |
| 5723 | C | T | 2.3% | 132 | 0.1085 | | | | | | | NS3 | S | | | | |
| 5730 | A | G | 1.5% | 135 | 0.0771 | | | | | | | NS3 | NS | TA | | | |
| 5777 | A | T | 20.7% | 188 | 0.5106 | | T | 4.5% | 1861 | 0.1828 | | NS3 | S | | | | |
| 5813 | T | C | 1.0% | 192 | 0.0579 | | | | | | | NS3 | S | | | | |
| 5817 | G | T | 2.1% | 194 | 0.1004 | | | | | | | NS3 | NS | AS | | | |
| 5822 | A | G | 2.1% | 195 | 0.1000 | | | | | | | NS3 | S | | | | |
| 5828 | A | G | 1.0% | 194 | 0.0574 | | | | | | | NS3 | S | | | | |
| 5831 | A | | | | | | G | 2.8% | 1845 | 0.1264 | | NS3 | S | | | | |
| 5834 | A | | | | | | G | 3.5% | 1834 | 0.1519 | | NS3 | S | | | | A/g |
| 5836 | C | | | | | | A | 1.0% | 1831 | 0.0577 | | NS3 | NS | TN | | | |
| 5837 | C | | | | | | A | 1.1% | 1833 | 0.0727 | | NS3 | S | | | | |
| 5851 | A | C | 0.9% | 211 | 0.0536 | | | | | | | NS3 | NS | KT | | | |
| 5868 | A | G | 2.0% | 205 | 0.0961 | | | | | | | NS3 | NS | IV | | | |
| 5876 | A | G | 0.9% | 213 | 0.0532 | | | | | | | NS3 | S | | | | |
| 5891 | G | A | 0.9% | 231 | 0.0497 | | | | | | | NS3 | S | | | | |
| 5894 | C | T | 2.6% | 235 | 0.1188 | | | | | | | NS3 | S | | | | |
| 5900 | A | T | 0.9% | 233 | 0.0494 | | | | | | | NS3 | S | | | | |
| 5903 | A | | | | | | G | 2.3% | 1846 | 0.1086 | | NS3 | S | | | | |
| 5915 | G | A | 0.8% | 239 | 0.0484 | | | | | | | NS3 | S | | | | |
| 5917 | C | T | 0.8% | 236 | 0.0489 | | | | | | | NS3 | NS | AV | | | |
| 5927 | C | T | 0.8% | 239 | 0.0484 | | | | | | | NS3 | S | | | | |

| Ref Position | Ref Base | Serum Base | Serum SNP % | Serum Depth | Serum entropy | | Culture Base | Culture SNP % | Culture Depth | Culture entropy | | Features | S/NS | Residue change | Serum PCR call | P3 PCR call | P5 PCR call |
|---|---|---|---|---|---|---|---|---|---|---|---|---|---|---|---|---|---|
| 5999 | C | T | 1.0% | 209 | 0.0540 | | | | | | | NS3 | S | | | | |
| 6000 | A | G | 1.0% | 207 | 0.0544 | | | | | | | NS3 | NS | MV | | | |
| 6003 | G | A | 1.0% | 205 | 0.0549 | | | | | | | NS3 | NS | AT | | | |
| 6008 | C | | | | | | T | 1.4% | 1975 | 0.0759 | | NS3 | S | | | | |
| 6016 | C | A | 1.0% | 195 | 0.0572 | | | | | | | NS3 | NS | TN | | | |
| 6029 | T | A | 1.0% | 193 | 0.0577 | | | | | | | NS3 | STOP | Y* | | | |
| 6044 | A | | | | | | G | 2.0% | 1815 | 0.1038 | | NS3 | S | | | | |
| 6050 | T | C | 1.2% | 162 | 0.0665 | | | | | | | NS3 | S | | | | |
| 6080 | A | G | 1.3% | 153 | 0.0697 | | | | | | | NS3 | S | | | | |
| 6083 | C | T | 9.4% | 149 | 0.3116 | | T | 3.8% | 1715 | 0.1618 | | NS3 | S | | | | |
| 6089 | C | T | 16.9% | 142 | 0.4543 | | T | 9.4% | 1630 | 0.3106 | | NS3 | S | | | | |
| 6104 | A | G | 1.5% | 130 | 0.0795 | | | | | | | NS3 | S | | | | |
| 6212 | C | | | | | | T | 26.4% | 1213 | 0.5767 | | NS3 | S | | C/t | C/t | C/t |
| 6299 | T | A | 2.7% | 110 | 0.1251 | | A | 8.3% | 1318 | 0.2842 | | NS3 | S | | | | T/a |
| 6320 | G | A | 1.8% | 112 | 0.0896 | | | | | | | NS3 | S | | | | |
| 6328 | T | A | 1.6% | 123 | 0.0831 | | | | | | | NS3 | NS | MK | | | |
| 6359 | G | A | 23.8% | 147 | 0.5489 | | | | | | | NS3 | S | | G/a | | |
| 6377 | G | A | 1.3% | 150 | 0.0708 | | | | | | | NS3 | S | | | | |
| 6385 | A | G | 1.3% | 158 | 0.0679 | | | | | | | NS3 | NS | KR | | | |
| 6389 | A | | | | | | G | 1.6% | 1772 | 0.0835 | | NS3 | S | | | | |
| 6395 | A | G | 1.3% | 156 | 0.0686 | | | | | | | NS3 | S | | | | |
| 6424 | A | G | 1.2% | 170 | 0.0640 | | | | | | | NS3 | NS | YC | | | |
| 6431 | G | A | 1.1% | 176 | 0.0622 | | | | | | | NS3 | S | | | | |
| 6437 | C | T | 1.1% | 174 | 0.0628 | | | | | | | NS3 | S | | | | |
| 6457 | T | C | 1.1% | 175 | 0.0625 | | | | | | | NS3 | NS | VA | | | |
| 6460 | C | | | | | | T | 1.2% | 1840 | 0.0629 | | NS3 | NS | TI | | | |

| Ref Position | Ref Base | Serum Base | Serum SNP % | Serum Depth | Serum entropy | | Culture Base | Culture SNP % | Culture Depth | Culture entropy | | Features | S/NS | Residue change | Serum PCR call | P3 PCR call | P5 PCR call |
|---|---|---|---|---|---|---|---|---|---|---|---|---|---|---|---|---|---|
| 6461 | A | | | | | | G | 1.1% | 1847 | 0.0603 | | NS3 | S | | | | |
| 6464 | A | | | | | | G | 2.2% | 1926 | 0.1070 | | NS3 | S | | | | |
| 6474 | T | G | 1.1% | 184 | 0.0600 | | | | | | | NS3 | NS | YD | | | |
| 6480 | A | G | 27.8% | 187 | 0.5911 | | | | | | | NS3 | NS | | A/g | | |
| 6482 | T | C | 1.1% | 187 | 0.0592 | | | | | | | NS3 | S | | | | |
| 6485 | G | A | 1.1% | 188 | 0.0589 | | | | | | | NS3 | S | | | | |
| 6488 | C | A | 1.1% | 189 | 0.0587 | | | | | | | NS3 | S | | | | |
| 6509 | A | T | 1.1% | 186 | 0.0594 | | | | | | | NS3 | S | | | | |
| 6519 | C | G | 1.0% | 195 | 0.0572 | | | | | | | NS3 | NS | PA | | | |
| 6525 | T | C | 1.1% | 188 | 0.0589 | | | | | | | NS3 | S | | | | |
| 6539 | G | A | 1.1% | 185 | 0.0597 | | A | 1.5% | 1858 | 0.0782 | | NS3 | S | | | | |
| 6545 | A | C | 1.1% | 185 | 0.0597 | | | | | | | NS3 | S | | | | |
| 6547 | G | T | 1.1% | 190 | 0.0584 | | | | | | | NS3 | NS | GV | | | |
| 6579 | T | C | 2.6% | 196 | 0.1188 | | | | | | | NS3 | NS | SP | | | |
| 6585 | A | G | 1.1% | 184 | 0.0600 | | G | 1.6% | 1750 | 0.0802 | | NS3 | NS | IV | | | |
| 6590 | C | T | 3.2% | 189 | 0.1610 | | | | | | | NS3 | S | | | | |
| 6606 | C | T | 1.0% | 191 | 0.0582 | | | | | | | NS3 | NS | LF | | | |
| 6620 | T | C | 3.7% | 190 | 0.1578 | | | | | | | NS3 | S | | | | |
| 6636 | C | A | 1.1% | 178 | 0.0616 | | | | | | | NS3 | NS | QK | | | |
| 6638 | G | A | 2.7% | 185 | 0.1096 | | | | | | | NS3 | S | | | | |
| 6644 | A | | | | | | G | 1.6% | 1773 | 0.0812 | | NS3 | S | | | | |
| 6659 | A | T | 2.0% | 197 | 0.1133 | | | | | | | NS3 | S | | | | |
| 6662 | T | C | 2.0% | 199 | 0.0831 | | | | | | | NS3 | S | | | | |
| 6692 | C | T | 1.6% | 188 | 0.0819 | | | | | | | NS3 | S | | | | |
| 6710 | G | A | 1.0% | 197 | 0.0567 | | | | | | | NS3 | S | | | | |
| 6779 | A | G | 0.9% | 220 | 0.0518 | | | | | | | NS3 | S | | | | |

| Ref Position | Ref Base | Serum Base | Serum SNP % | Serum Depth | Serum entropy | | Culture Base | Culture SNP % | Culture Depth | Culture entropy | | Features | S/NS | Residue change | Serum PCR call | P3 PCR call | P5 PCR call |
|---|---|---|---|---|---|---|---|---|---|---|---|---|---|---|---|---|---|
| 6782 | A | G | 5.3% | 227 | 0.2307 | | G | 1.9% | 1790 | 0.0924 | | NS3 | S | | | | |
| 6808 | G | A | 0.9% | 219 | 0.0520 | | | | | | | NS3 | STOP | W* | | | |
| 6824 | G | A | 0.9% | 212 | 0.0534 | | A | 6.1% | 1792 | 0.2424 | | NS3 | S | | | | |
| 6833 | G | A | 1.9% | 208 | 0.0950 | | | | | | | NS3 | S | | | | |
| 6836 | A | G | 1.0% | 208 | 0.0542 | | | | | | | NS3 | S | | | | |
| 6855 | C | T | 3.4% | 204 | 0.1494 | | | | | | | NS3 | S | | | | |
| 6860 | C | | | | | | T | 2.6% | 1683 | 0.1189 | | NS3 | S | | | | |
| 6877 | A | G | 1.0% | 194 | 0.0574 | | | | | | | NS3 | NS | EG | | | |
| 6878 | A | T | 2.1% | 193 | 0.1008 | | | | | | | NS3 | NS | ED | | | |
| 6881 | C | A | 4.1% | 194 | 0.1719 | | | | | | | NS3 | NS | DE | | | |
| 6882 | T | C | 2.1% | 195 | 0.1000 | | | | | | | NS3 | S | | | | |
| 6885 | C | T | 3.0% | 198 | 0.1358 | | | | | | | NS3 | NS | PS | | | |
| 6911 | C | | | | | | T | 2.4% | 1734 | 0.1187 | | NS3 | S | | | | |
| 6916 | C | T | 1.1% | 178 | 0.0616 | | | | | | | NS3 | NS | TI | | | |
| 6926 | A | G | 1.2% | 168 | 0.0646 | | | | | | | NS3 | S | | | | |
| 6932 | G | A | 1.8% | 163 | 0.0722 | | | | | | | NS3 | S | | | | |
| 6950 | C | | | | | | T | 2.1% | 1799 | 0.1002 | | NS3 | S | | | | |
| 6971 | A | G | 1.2% | 163 | 0.0662 | | | | | | | NS3 | S | | | | |
| 6988 | T | | | | | | A | 1.0% | 1679 | 0.0634 | | NS3 | NS | IK | | | |
| 6998 | A | G | 6.2% | 145 | 0.2326 | | | | | | | NS3 | S | | | | |
| 7007 | A | G | 5.8% | 137 | 0.2225 | | G | 3.8% | 1433 | 0.1605 | | NS3 | S | | | | |
| 7016 | C | | | | | | T | 1.7% | 1498 | 0.0923 | | NS3 | S | | | | |
| 7049 | A | | | | | | G | 5.2% | 1508 | 0.1996 | | NS3 | S | | | | |
| 7052 | A | G | 1.4% | 146 | 0.0724 | | | | | | | NS3 | S | | | | |
| 7058 | T | G | 1.4% | 144 | 0.0732 | | | | | | | NS3 | NS | DE | | | |
| 7071 | A | G | 1.2% | 171 | 0.0637 | | G | 3.2% | 1792 | 0.1410 | | NS3 | NS | IV | | | |

| Ref Position | Ref Base | Serum Base | Serum SNP % | Serum Depth | Serum entropy | | Culture Base | Culture SNP % | Culture Depth | Culture entropy | | Features | S/NS | Residue change | Serum PCR call | P3 PCR call | P5 PCR call |
|---|---|---|---|---|---|---|---|---|---|---|---|---|---|---|---|---|---|
| 7088 | T | C | 2.3% | 172 | 0.1105 | | | | | | | NS3 | S | | | | |
| 7095 | C | A | 1.2% | 169 | 0.0643 | | | | | | | NS3 | NS | LM | | | |
| 7103 | T | C | 1.1% | 174 | 0.0628 | | C | 1.7% | 1800 | 0.0853 | | NS3 | S | | | | |
| 7127 | C | T | 1.1% | 174 | 0.0628 | | | | | | | NS3 | S | | | | |
| 7138 | A | G | 1.2% | 164 | 0.0659 | | | | | | | NS3 | NS | NS | | | |
| 7147 | T | C | 1.1% | 179 | 0.0613 | | | | | | | NS3 | NS | VA | | | |
| 7150 | T | C | 1.6% | 184 | 0.0653 | | | | | | | NS3 | NS | VA | | | |
| 7165 | C | G | 1.0% | 192 | 0.0579 | | | | | | | NS3 | NS | AG | | | |
| 7169 | G | T | 1.6% | 188 | 0.0819 | | | | | | | NS3 | S | | | | |
| 7183 | G | T | 1.1% | 182 | 0.0605 | | | | | | | NS3 | NS | GV | | | |
| 7199 | G | A | 1.2% | 169 | 0.0643 | | | | | | | NS4A | S | | | | |
| 7203 | G | A | 1.2% | 162 | 0.0665 | | | | | | | NS4A | NS | AT | | | |
| 7221 | T | A | 1.2% | 168 | 0.0646 | | | | | | | NS4A | NS | FI | | | |
| 7226 | G | | | | | | A | 1.1% | 1745 | 0.0675 | | NS4A | S | | | | |
| 7233 | G | T | 1.1% | 181 | 0.0608 | | | | | | | NS4A | NS | GC | | | |
| 7239 | C | A | 1.0% | 192 | 0.0579 | | | | | | | NS4A | NS | QK | | | |
| 7244 | T | A | 11.7% | 188 | 0.3609 | | A | 37.7% | 2116 | 0.6624 | | NS4A | S | | | | |
| 7247 | A | G | 3.7% | 189 | 0.1459 | | G | 1.1% | 2142 | 0.0635 | | NS4A | S | | | | |
| 7253 | G | A | 1.1% | 186 | 0.0594 | | | | | | | NS4A | S | | | | |
| 7254 | A | G | 1.1% | 184 | 0.0600 | | | | | | | NS4A | NS | RG | | | |
| 7256 | G | A | 1.1% | 185 | 0.0597 | | | | | | | NS4A | S | | | | |
| 7259 | T | A | 1.1% | 186 | 0.0594 | | | | | | | NS4A | NS | HQ | | | |
| 7277 | T | | | | | | C | 4.6% | 2165 | 0.1838 | | NS4A | S | | | | |
| 7328 | T | C | 1.3% | 149 | 0.0712 | | | | | | | NS4A | S | | | | |
| 7337 | C | | | | | | T | 1.3% | 2146 | 0.0644 | | NS4A | S | | | | |
| 7368 | T | C | 3.6% | 165 | 0.1562 | | | | | | | NS4A | S | | | | |

| Ref Position | Ref Base | Serum Base | Serum SNP % | Serum Depth | Serum entropy | | Culture Base | Culture SNP % | Culture Depth | Culture entropy | | Features | S/NS | Residue change | Serum PCR call | P3 PCR call | P5 PCR call |
|---|---|---|---|---|---|---|---|---|---|---|---|---|---|---|---|---|---|
| 7379 | G | A | 1.3% | 159 | 0.0675 | | | | | | | NS4A | S | | | | |
| 7416 | T | | | | | | A | 4.3% | 2355 | 0.1889 | | NS4B | NS | | | | |
| 7427 | A | G | 1.5% | 134 | 0.0776 | | | | | | | NS4B | S | | | | |
| 7436 | G | T | 7.2% | 138 | 0.2962 | | T | 36.8% | 2351 | 0.6808 | | NS4B | S | | G/t | | K |
| 7465 | G | | | | | | A | 1.4% | 1908 | 0.0761 | | NS4B | NS | RK | | | |
| 7472 | A | G | 1.6% | 122 | 0.0836 | | | | | | | NS4B | S | | | | |
| 7475 | C | T | 1.6% | 123 | 0.0831 | | | | | | | NS4B | S | | | | |
| 7484 | G | A | 1.6% | 125 | 0.0820 | | | | | | | NS4B | S | | | | |
| 7485 | T | G | 1.6% | 124 | 0.0826 | | | | | | | NS4B | NS | FV | | | |
| 7497 | G | A | 1.5% | 130 | 0.0795 | | | | | | | NS4B | NS | VI | | | |
| 7519 | T | | | | | | C | 1.8% | 1634 | 0.0999 | | NS4B | NS | VA | | | T/c |
| 7520 | A | G | 6.2% | 146 | 0.2315 | | G | 7.2% | 1632 | 0.2586 | | NS4B | S | | | | |
| 7521 | | | | | | | | | | | | NS4B | NS | KE | | | A/g |
| 7522 | A | | | | | | G | 2.1% | 1608 | 0.1001 | | NS4B | NS | KR | | | |
| 7524 | | | | | | | | | | | | NS4B | NS | KR | | | A/g |
| 7527 | T | | | | | | A | 1.1% | 1637 | 0.0632 | | NS4B | NS | LI | | | |
| 7547 | | | | | | | | | | | | NS4B | S | | | | A/g |
| 7550 | T | A | 2.7% | 148 | 0.1243 | | | | | | | NS4B | NS | DE | | | |
| 7573 | G | C | 1.2% | 165 | 0.0655 | | | | | | | NS4B | NS | GA | | | |
| 7579 | G | T | 1.2% | 172 | 0.0634 | | | | | | | NS4B | NS | WL | | | |
| 7582 | G | A | 1.2% | 173 | 0.0631 | | | | | | | NS4B | NS | GE | | | |
| 7589 | T | C | 1.2% | 172 | 0.0634 | | | | | | | NS4B | S | | | | |
| 7597 | T | A | 1.1% | 190 | 0.0584 | | | | | | | NS4B | NS | LH | | | |
| 7613 | T | C | 1.0% | 205 | 0.0549 | | | | | | | NS4B | S | | | | |
| 7623 | G | A | 1.0% | 207 | 0.0544 | | | | | | | NS4B | NS | GR | | | |
| 7625 | G | A | 1.0% | 208 | 0.0542 | | | | | | | NS4B | S | | | | |

| Ref Position | Ref Base | Serum Base | Serum SNP % | Serum Depth | Serum entropy | | Culture Base | Culture SNP % | Culture Depth | Culture entropy | | Features | S/NS | Residue change | Serum PCR call | P3 PCR call | P5 PCR call |
|---|---|---|---|---|---|---|---|---|---|---|---|---|---|---|---|---|---|
| 7634 | A | G | 1.4% | 209 | 0.0752 | | | | | | | NS4B | S | | | | |
| 7643 | T | C | 0.9% | 216 | 0.0526 | | | | | | | NS4B | S | | | | |
| 7652 | G | | | | | | A | 4.1% | 1695 | 0.1702 | | NS4B | S | | | | |
| 7662 | C | G | 0.9% | 216 | 0.0526 | | | | | | | NS4B | NS | LV | | | |
| 7667 | C | T | 0.9% | 214 | 0.0530 | | | | | | | NS4B | S | | | | |
| 7670 | C | T | 3.3% | 214 | 0.1440 | | T | 1.4% | 1623 | 0.0718 | | NS4B | S | | | | |
| 7694 | C | T | 0.9% | 213 | 0.0532 | | T | 4.0% | 1699 | 0.1680 | | NS4B | S | | | | |
| 7697 | C | A | 22.6% | 212 | 0.5349 | | | | | | | NS4B | S | | C/a | | |
| 7749 | T | C | 1.0% | 210 | 0.0538 | | | | | | | NS4B | NS | FL | | | |
| 7760 | T | C | 10.5% | 210 | 0.3354 | | C | 5.0% | 1716 | 0.1971 | | NS4B | S | | | | |
| 7762 | C | A | 1.0% | 210 | 0.0538 | | | | | | | NS4B | NS | SY | | | |
| 7772 | G | A | 1.0% | 210 | 0.0538 | | | | | | | NS4B | S | | | | |
| 7802 | G | | | | | | A | 1.4% | 1741 | 0.0727 | | NS4B | S | | | | G/a |
| 7811 | C | T | 2.1% | 194 | 0.1004 | | | | | | | NS4B | S | | | | |
| 7815 | C | T | 8.0% | 188 | 0.2782 | | T | 2.7% | 1732 | 0.1231 | | NS4B | S | | | | |
| 7846 | A | G | 1.1% | 178 | 0.0616 | | | | | | | NS4B | NS | YC | | | |
| 7856 | C | A | 1.1% | 179 | 0.0613 | | | | | | | NS4B | S | | | | |
| 7862 | A | G | 2.4% | 164 | 0.0973 | | | | | | | NS4B | S | | | | |
| 7878 | C | T | 1.1% | 182 | 0.0605 | | | | | | | NS4B | S | | | | |
| 7895 | T | | | | | | C | 1.2% | 1820 | 0.0646 | | NS4B | S | | | | |
| 7899 | A | G | 1.1% | 189 | 0.0587 | | | | | | | NS4B | NS | TA | | | |
| 7910 | A | G | 1.1% | 188 | 0.0589 | | | | | | | NS4B | S | | | | |
| 7922 | C | | | | | | A | 2.2% | 1671 | 0.1040 | | NS4B | S | | | | |
| 7923 | C | A | 2.8% | 213 | 0.1283 | | | | | | | NS4B | NS | PT | | | |
| 7926 | A | T | 0.9% | 215 | 0.0528 | | | | | | | NS4B | NS | TS | | | |
| 7934 | G | | | | | | A | 2.4% | 1756 | 0.1129 | | NS4B | S | | | | |

| Ref Position | Ref Base | Serum Base | Serum SNP % | Serum Depth | Serum entropy | | Culture Base | Culture SNP % | Culture Depth | Culture entropy | | Features | S/NS | Residue change | Serum PCR call | P3 PCR call | P5 PCR call |
|---|---|---|---|---|---|---|---|---|---|---|---|---|---|---|---|---|---|
| 7950 | C | T | 0.9% | 219 | 0.0520 | | | | | | | NS4B | S | | | | |
| 7955 | C | | | | | | T | 1.3% | 1664 | 0.0683 | | NS4B | S | | | | |
| 7981 | C | T | 0.6% | 315 | 0.0385 | | | | | | | NS4B | NS | SL | | | |
| 7982 | A | G | 0.6% | 313 | 0.0387 | | | | | | | NS4B | S | | | | |
| 7991 | A | G | 0.9% | 336 | 0.0510 | | | | | | | NS4B | S | | | | |
| 8001 | G | T | 1.9% | 324 | 0.0922 | | | | | | | NS4B | NS | DY | | | |
| 8003 | T | C | 3.2% | 317 | 0.1401 | | C | 1.7% | 2765 | 0.0846 | | NS4B | S | | | | |
| 8018 | G | A | 6.6% | 334 | 0.2428 | | A | 3.5% | 3706 | 0.1507 | | NS4B | S | | | | |
| 8030 | A | C | 1.2% | 333 | 0.0544 | | C | 4.2% | 3680 | 0.1855 | | NS4B | S | | | | |
| 8042 | A | G | 1.8% | 332 | 0.0904 | | | | | | | NS4B | NS | IM | | | |
| 8090 | A | | | | | | G | 1.6% | 3291 | 0.0803 | | NS4B | S | | | | |
| 8103 | G | | | | | | A | 1.6% | 3153 | 0.0828 | | NS4B | NS | AT | | | |
| 8112 | G | A | 0.7% | 295 | 0.0406 | | | | | | | NS4B | NS | AT | | | |
| 8114 | C | A | 0.7% | 297 | 0.0404 | | | | | | | NS4B | S | | | | |
| 8126 | C | | | | | | T | 1.3% | 3055 | 0.0684 | | NS4B | S | | | | |
| 8130 | C | T | 0.7% | 295 | 0.0406 | | | | | | | NS4B | STOP | Q* | | | |
| 8132 | G | C | 0.7% | 291 | 0.0411 | | | | | | | NS4B | NS | QH | | | |
| 8142 | T | C | 0.7% | 289 | 0.0413 | | | | | | | NS4B | S | | | | |
| 8145 | C | T | 0.7% | 289 | 0.0413 | | | | | | | NS4B | NS | LF | | | |
| 8162 | A | G | 3.0% | 263 | 0.1362 | | | | | | | NS4B | S | | | | |
| 8175 | G | A | 0.8% | 260 | 0.0451 | | | | | | | NS4B | NS | DN | | | |
| 8183 | A | G | 0.8% | 258 | 0.0454 | | | | | | | NS4B | S | | | | |
| 8188 | C | T | 1.0% | 198 | 0.0565 | | | | | | | NS4B | NS | TM | | | |
| 8189 | G | A | 1.0% | 201 | 0.0558 | | | | | | | NS4B | S | | | | |
| 8195 | G | A | 2.0% | 196 | 0.0996 | | A | 1.5% | 2094 | 0.0848 | | NS4B | S | | | | |
| 8201 | T | C | 1.4% | 145 | 0.0728 | | | | | | | NS4B | S | | | | |

| Ref Position | Ref Base | Serum Base | Serum SNP % | Serum Depth | Serum entropy | | Culture Base | Culture SNP % | Culture Depth | Culture entropy | | Features | S/NS | Residue change | Serum PCR call | P3 PCR call | P5 PCR call |
|---|---|---|---|---|---|---|---|---|---|---|---|---|---|---|---|---|---|
| 8211 | C | | | | | | A | 1.0% | 864 | 0.0589 | | NS4B | NS | PT | | | |
| 8223 | A | | | | | | C | 1.4% | 292 | 0.0606 | | NS4B | NS | IL | | | |
| 8232 | T | | | | | | C | 1.8% | 109 | 0.0915 | | NS4B | S | | | | |
| 8236 | T | A | 2.2% | 135 | 0.1066 | | | | | | | NS4B | NS | FY | | | |
| 8257 | G | A | 1.7% | 120 | 0.0848 | | | | | | | NS4B | NS | GD | | | |
| 8258 | T | C | 1.7% | 119 | 0.0853 | | | | | | | NS4B | S | | | | |
| 8264 | C | T | 24.3% | 136 | 0.6573 | | T | 6.3% | 347 | 0.2362 | | NS4B | S | | C/t | | |
| 8270 | A | T | 1.6% | 127 | 0.0810 | | | | | | | NS4B | NS | RS | | | |
| 8330 | G | A | 1.2% | 165 | 0.0655 | | | | | | | NS4B | S | | | | |
| 8341 | G | A | 1.2% | 162 | 0.0665 | | | | | | | NS4B | NS | GD | | | |
| 8357 | A | G | 2.4% | 165 | 0.1141 | | | | | | | NS4B | S | | | | |
| 8358 | C | A | 1.2% | 167 | 0.0649 | | | | | | | NS4B | NS | LM | | | |
| 8375 | T | | | | | | C | 1.7% | 1397 | 0.0868 | | NS4B | S | | | | |
| 8378 | C | T | 2.4% | 168 | 0.1125 | | | | | | | NS4B | S | | | | |
| 8399 | A | G | 1.3% | 157 | 0.0682 | | | | | | | NS4B | S | | | | |
| 8442 | T | C | 2.4% | 166 | 0.1136 | | | | | | | NS5A | S | | | | |
| 8460 | A | G | 2.5% | 158 | 0.1181 | | | | | | | NS5A | NS | KE | | | |
| 8476 | G | A | 1.3% | 149 | 0.0712 | | | | | | | NS5A | NS | GE | | | |
| 8480 | G | C | 1.3% | 150 | 0.0708 | | | | | | | NS5A | S | | | | |
| 8483 | G | A | 1.3% | 149 | 0.0712 | | | | | | | NS5A | S | | | | |
| 8485 | A | G | 1.4% | 147 | 0.0720 | | | | | | | NS5A | NS | KR | | | |
| 8489 | A | G | 2.7% | 146 | 0.1256 | | G | 1.7% | 1798 | 0.0814 | | NS5A | NS | IM | | | |
| 8535 | G | | | | | | A | 4.7% | 1733 | 0.1888 | | NS5A | NS | GS | | | |
| 8549 | C | T | 1.4% | 139 | 0.0753 | | | | | | | NS5A | S | | | | |
| 8553 | T | G | 1.4% | 139 | 0.0753 | | | | | | | NS5A | NS | LV | | | |
| 8561 | A | | | | | | T | 1.6% | 1614 | 0.0873 | | NS5A | S | | | | |

| Ref Position | Ref Base | Serum Base | Serum SNP % | Serum Depth | Serum entropy | | Culture Base | Culture SNP % | Culture Depth | Culture entropy | | Features | S/NS | Residue change | Serum PCR call | P3 PCR call | P5 PCR call |
|---|---|---|---|---|---|---|---|---|---|---|---|---|---|---|---|---|---|
| 8572 | T | C | 5.2% | 134 | 0.2051 | | | | | | | NS5A | NS | LS | | | |
| 8584 | T | C | 11.5% | 130 | 0.3576 | | C | 1.4% | 1760 | 0.0763 | | NS5A | NS | IT | | | |
| 8591 | C | T | 2.1% | 140 | 0.0819 | | | | | | | NS5A | S | | | | |
| 8618 | C | | | | | | A | 1.5% | 1985 | 0.0762 | | NS5A | S | | | | |
| 8627 | A | G | 9.9% | 162 | 0.3224 | | | | | | | NS5A | S | | | | |
| 8633 | T | | | | | | C | 1.5% | 2074 | 0.0745 | | NS5A | S | | | | |
| 8634 | A | G | 1.8% | 164 | 0.0913 | | | | | | | NS5A | NS | KE | | | |
| 8636 | A | | | | | | G | 1.3% | 2073 | 0.0702 | | NS5A | S | | | | |
| 8637 | C | | | | | | T | 1.1% | 2072 | 0.0635 | | NS5A | NS | LF | | | |
| 8639 | T | C | 1.2% | 166 | 0.0652 | | | | | | | NS5A | S | | | | |
| 8645 | G | | | | | | A | 5.7% | 1993 | 0.2192 | | NS5A | S | | | | |
| 8666 | C | T | 2.0% | 151 | 0.0975 | | | | | | | NS5A | S | | | | |
| 8669 | G | | | | | | T | 1.9% | 1889 | 0.0968 | | NS5A | S | | | | |
| 8674 | A | G | 38.7% | 150 | 0.6672 | | G | 3.5% | 1872 | 0.1526 | | NS5A | NS | KR | A/g | | A/g |
| 8677 | A | G | 1.4% | 145 | 0.0728 | | | | | | | NS5A | NS | NS | | | |
| 8689 | G | C | 1.3% | 158 | 0.0679 | | | | | | | NS5A | NS | RT | | | |
| 8693 | G | | | | | | A | 1.4% | 1829 | 0.0704 | | NS5A | S | | | | |
| 8694 | C | A | 2.5% | 163 | 0.0978 | | | | | | | NS5A | NS | PT | | | |
| 8698 | T | C | 1.2% | 166 | 0.0652 | | | | | | | NS5A | NS | VA | | | |
| 8723 | T | C | 1.3% | 156 | 0.0686 | | | | | | | NS5A | S | | | | |
| 8746 | A | G | 1.2% | 166 | 0.0652 | | | | | | | NS5A | NS | KR | | | |
| 8750 | A | G | 2.4% | 164 | 0.1147 | | | | | | | NS5A | S | | | | |
| 8771 | A | | | | | | G | 1.2% | 1880 | 0.0618 | | NS5A | S | | | | |
| 8780 | C | T | 1.1% | 175 | 0.0625 | | | | | | | NS5A | S | | | | |
| 8783 | C | T | 1.1% | 176 | 0.0622 | | | | | | | NS5A | S | | | | |
| 8787 | A | T | 1.1% | 179 | 0.0613 | | | | | | | NS5A | STOP | K* | | | |

| Ref Position | Ref Base | Serum Base | Serum SNP % | Serum Depth | Serum entropy | | Culture Base | Culture SNP % | Culture Depth | Culture entropy | | Features | S/NS | Residue change | Serum PCR call | P3 PCR call | P5 PCR call |
|---|---|---|---|---|---|---|---|---|---|---|---|---|---|---|---|---|---|
| 8816 | T | C | 1.2% | 164 | 0.0659 | | C | 5.3% | 1456 | 0.2356 | | NS5A | S | | | | |
| 8820 | G | A | 1.2% | 164 | 0.0659 | | | | | | | NS5A | NS | GR | | | |
| 8821 | G | A | 1.2% | 166 | 0.0652 | | | | | | | NS5A | NS | GE | | | |
| 8827 | C | A | 1.8% | 169 | 0.0892 | | A | 1.9% | 1509 | 0.1027 | | NS5A | NS | TK | | | |
| 8828 | A | G | 1.2% | 169 | 0.0643 | | | | | | | NS5A | S | | | | |
| 8829 | C | A | 1.2% | 168 | 0.0646 | | | | | | | NS5A | NS | LI | | | |
| 8831 | C | G | 1.2% | 166 | 0.0652 | | | | | | | NS5A | S | | | | |
| 8843 | T | | | | | | C | 3.1% | 1513 | 0.1368 | | NS5A | S | | | | |
| 8848 | G | T | 1.2% | 166 | 0.0652 | | | | | | | NS5A | NS | WL | | | |
| 8868 | A | T | 1.2% | 162 | 0.0665 | | | | | | | NS5A | NS | IL | | | |
| 8876 | G | | | | | | A | 4.2% | 1885 | 0.1728 | | NS5A | S | | | | |
| 8879 | G | A | 1.2% | 161 | 0.0669 | | | | | | | NS5A | S | | | | |
| 8886 | A | G | 1.3% | 159 | 0.0675 | | | | | | | NS5A | NS | RG | | | |
| 8891 | C | T | 1.3% | 158 | 0.0679 | | | | | | | NS5A | S | | | | |
| 8912 | C | T | 1.2% | 166 | 0.0652 | | | | | | | NS5A | S | | | | |
| 8915 | A | G | 1.2% | 167 | 0.0649 | | | | | | | NS5A | S | | | | |
| 8963 | T | A | 2.2% | 136 | 0.0839 | | | | | | | NS5A | STOP | C* | | | |
| 8964 | G | C | 1.5% | 135 | 0.0771 | | | | | | | NS5A | NS | AP | | | |
| 8981 | C | A | 1.5% | 137 | 0.0762 | | A | 1.9% | 1870 | 0.0985 | | NS5A | NS | NK | | | |
| 8983 | C | | | | | | A | 1.3% | 1837 | 0.0746 | | NS5A | NS | TK | | | |
| 8984 | A | T | 3.1% | 130 | 0.1374 | | T | 3.7% | 1829 | 0.1740 | | NS5A | S | | | | |
| 8987 | G | | | | | | A | 1.4% | 1753 | 0.0765 | | NS5A | S | | | | |
| 9001 | T | | | | | | A | 1.5% | 1632 | 0.0862 | | NS5A | NS | MK | | | |
| 9026 | C | T | 3.2% | 156 | 0.1418 | | T | 1.0% | 1726 | 0.0559 | | NS5A | S | | | | |
| 9035 | T | | | | | | C | 4.2% | 1724 | 0.1741 | | NS5A | S | | | | |
| 9039 | T | A | 1.3% | 157 | 0.0682 | | | | | | | NS5A | NS | ST | | | |

| Ref Position | Ref Base | Serum Base | Serum SNP % | Serum Depth | Serum entropy | | Culture Base | Culture SNP % | Culture Depth | Culture entropy | | Features | S/NS | Residue change | Serum PCR call | P3 PCR call | P5 PCR call |
|---|---|---|---|---|---|---|---|---|---|---|---|---|---|---|---|---|---|
| 9056 | T | | | | | | C | 1.2% | 1604 | 0.0577 | | NS5A | S | | | | |
| 9059 | T | C | 8.5% | 142 | 0.2896 | | | | | | | NS5A | S | | | | |
| 9073 | G | A | 1.5% | 137 | 0.0762 | | | | | | | NS5A | NS | RK | | | |
| 9079 | A | | | | | | C | 1.1% | 1516 | 0.0585 | | NS5A | NS | NT | | | |
| 9083 | T | C | 1.5% | 132 | 0.0785 | | | | | | | NS5A | S | | | | |
| 9089 | G | | | | | | A | 2.9% | 1288 | 0.1430 | | NS5A | S | | | | |
| 9093 | G | A | 2.3% | 133 | 0.1078 | | A | 3.2% | 1392 | 0.1425 | | NS5A | NS | EK | | | |
| 9098 | A | | | | | | C | 2.5% | 1664 | 0.1178 | | NS5A | S | | | | |
| 9140 | G | | | | | | A | 1.1% | 2229 | 0.0599 | | NS5A | S | | | | |
| 9150 | A | G | 1.5% | 197 | 0.0788 | | G | 3.1% | 2281 | 0.1386 | | NS5A | NS | MV | | | |
| 9160 | T | A | 1.0% | 193 | 0.0577 | | | | | | | NS5A | NS | IK | | | |
| 9165 | C | T | 1.0% | 203 | 0.0553 | | | | | | | NS5A | NS | PS | | | |
| 9170 | G | | | | | | A | 1.0% | 2391 | 0.0591 | | NS5A | S | | | | |
| 9171 | C | | | | | | T | 1.7% | 2399 | 0.0852 | | NS5A | S | | | | |
| 9182 | A | G | 2.3% | 215 | 0.1105 | | | | | | | NS5A | S | | | | |
| 9183 | G | A | 0.9% | 215 | 0.0528 | | | | | | | NS5A | NS | VI | | | |
| 9188 | C | T | 0.9% | 217 | 0.0524 | | | | | | | NS5A | S | | | | |
| 9199 | T | C | 0.9% | 222 | 0.0514 | | | | | | | NS5A | NS | VA | | | |
| 9209 | C | T | 0.9% | 235 | 0.0490 | | | | | | | NS5A | S | | | | |
| 9210 | A | | | | | | T | 2.8% | 2757 | 0.1275 | | NS5A | NS | SC | | | |
| 9225 | G | A | 2.1% | 236 | 0.1026 | | | | | | | NS5A | NS | VI | | | |
| 9239 | A | G | 1.7% | 233 | 0.0868 | | | | | | | NS5A | S | | | | |
| 9248 | T | C | 16.6% | 235 | 0.4494 | | C | 8.1% | 2829 | 0.2819 | | NS5A | S | | T/c | | |
| 9267 | A | | | | | | G | 1.4% | 3026 | 0.0695 | | NS5A | NS | RG | | | |
| 9269 | G | A | 3.1% | 255 | 0.1395 | | | | | | | NS5A | S | | | | |
| 9271 | G | A | 0.8% | 256 | 0.0457 | | | | | | | NS5A | NS | GE | | | |

| Ref Position | Ref Base | Serum Base | Serum SNP % | Serum Depth | Serum entropy | | Culture Base | Culture SNP % | Culture Depth | Culture entropy | | Features | S/NS | Residue change | Serum PCR call | P3 PCR call | P5 PCR call |
|---|---|---|---|---|---|---|---|---|---|---|---|---|---|---|---|---|---|
| 9272 | A | G | 0.8% | 255 | 0.0458 | | | | | | | NS5A | S | | | | |
| 9275 | C | T | 1.6% | 255 | 0.0807 | | | | | | | NS5A | S | | | | |
| 9293 | G | A | 15.7% | 254 | 0.4355 | | | | | | | NS5A | S | | G/a | | |
| 9311 | G | A | 1.7% | 232 | 0.0871 | | | | | | | NS5A | S | | | | |
| 9313 | T | A | 1.7% | 233 | 0.0868 | | | | | | | NS5A | NS | IK | | | |
| 9315 | G | A | 0.9% | 233 | 0.0494 | | | | | | | NS5A | NS | EK | | | |
| 9323 | C | T | 0.9% | 219 | 0.0520 | | | | | | | NS5A | S | | | | |
| 9324 | G | T | 0.9% | 219 | 0.0520 | | | | | | | NS5A | NS | AS | | | |
| 9337 | A | G | 1.9% | 215 | 0.0926 | | | | | | | NS5A | NS | QR | | | |
| 9343 | C | T | 0.9% | 212 | 0.0534 | | | | | | | NS5A | NS | TM | | | |
| 9353 | T | C | 3.6% | 221 | 0.1557 | | | | | | | NS5A | S | | | | |
| 9362 | T | | | | | | A | 1.5% | 2223 | 0.0772 | | NS5A | NS | DE | | | |
| 9374 | C | A | 0.9% | 222 | 0.0514 | | | | | | | NS5A | STOP | Y* | | | |
| 9381 | C | T | 1.8% | 227 | 0.0886 | | | | | | | NS5A | NS | PS | | | |
| 9383 | T | C | 2.2% | 228 | 0.1202 | | C | 2.8% | 2313 | 0.1446 | | NS5A | S | | | | |
| 9387 | C | A | 1.3% | 231 | 0.0693 | | | | | | | NS5A | NS | LM | | | |
| 9392 | G | A | 0.9% | 227 | 0.0505 | | A | 5.8% | 2252 | 0.2236 | | NS5A | S | | | | |
| 9396 | A | G | 0.9% | 233 | 0.0494 | | | | | | | NS5A | NS | ND | | | |
| 9406 | T | A | 2.7% | 225 | 0.0979 | | | | | | | NS5A | NS | VE | | | |
| 9407 | G | A | 0.9% | 225 | 0.0508 | | | | | | | NS5A | S | | | | |
| 9411 | G | | | | | | A | 1.1% | 2081 | 0.0655 | | NS5A | NS | EK | | | |
| 9415 | T | | | | | | A | 1.1% | 2059 | 0.0591 | | NS5A | NS | IK | | | |
| 9430 | C | | | | | | T | 1.1% | 2056 | 0.0614 | | NS5A | NS | AV | | | |
| 9437 | T | | | | | | A | 10.9% | 2162 | 0.3501 | | NS5A | S | | | | T/a |
| 9443 | T | | | | | | C | 4.2% | 2206 | 0.1747 | | NS5A | S | | | | |
| 9449 | C | T | 0.9% | 233 | 0.0494 | | | | | | | NS5A | S | | | | |

| Ref Position | Ref Base | Serum Base | Serum SNP % | Serum Depth | Serum entropy | | Culture Base | Culture SNP % | Culture Depth | Culture entropy | | Features | S/NS | Residue change | Serum PCR call | P3 PCR call | P5 PCR call |
|---|---|---|---|---|---|---|---|---|---|---|---|---|---|---|---|---|---|
| 9472 | C | T | 0.9% | 227 | 0.0505 | | | | | | | NS5A | NS | SL | | | |
| 9480 | G | T | 0.8% | 251 | 0.0464 | | | | | | | NS5A | NS | AS | | | |
| 9482 | A | T | 0.8% | 253 | 0.0461 | | | | | | | NS5A | S | | | | |
| 9489 | G | A | 0.8% | 255 | 0.0458 | | | | | | | NS5A | NS | AT | | | |
| 9491 | T | C | 0.8% | 256 | 0.0457 | | | | | | | NS5A | S | | | | |
| 9509 | T | C | 25.2% | 258 | 0.5645 | | | | | | | NS5A | S | | T/c | | |
| 9515 | T | C | 1.6% | 252 | 0.0815 | | | | | | | NS5A | S | | | | |
| 9516 | G | | | | | | A | 2.6% | 2190 | 0.1178 | | NS5A | NS | GS | | | |
| 9517 | G | A | 0.8% | 253 | 0.0461 | | | | | | | NS5A | NS | GD | | | |
| 9518 | T | C | 0.8% | 252 | 0.0463 | | | | | | | NS5A | S | | | | |
| 9519 | A | | | | | | G | 3.1% | 2189 | 0.1368 | | NS5A | NS | ND | | | |
| 9546 | G | A | 0.8% | 266 | 0.0443 | | | | | | | NS5A | NS | AT | | | |
| 9577 | G | A | 0.7% | 296 | 0.0405 | | | | | | | NS5A | NS | RK | | | |
| 9582 | A | T | 0.7% | 299 | 0.0402 | | | | | | | NS5A | NS | IF | | | |
| 9585 | G | A | 0.7% | 301 | 0.0399 | | | | | | | NS5A | NS | DN | | | |
| 9594 | C | A | 0.7% | 304 | 0.0396 | | | | | | | NS5A | NS | LI | | | |
| 9596 | A | G | 1.3% | 303 | 0.0702 | | | | | | | NS5A | S | | | | |
| 9602 | A | G | 1.0% | 307 | 0.0425 | | | | | | | NS5A | S | | | | |
| 9607 | T | A | 1.0% | 303 | 0.0430 | | | | | | | NS5A | NS | VD | | | |
| 9618 | G | A | 0.7% | 306 | 0.0394 | | | | | | | NS5A | NS | GR | | | |
| 9626 | C | | | | | | T | 1.2% | 2550 | 0.0630 | | NS5A | S | | | | |
| 9641 | C | A | 0.7% | 283 | 0.0420 | | | | | | | NS5A | S | | | | |
| 9660 | G | C | 0.7% | 287 | 0.0416 | | | | | | | NS5A | NS | GR | | | |
| 9663 | A | G | 0.7% | 281 | 0.0423 | | | | | | | NS5A | NS | KE | | | |
| 9671 | A | G | 0.7% | 282 | 0.0422 | | | | | | | NS5A | S | | | | |
| 9718 | A | G | 0.9% | 222 | 0.0514 | | G | 1.9% | 2281 | 0.1001 | | NS5A | NS | EG | | | |

| Ref Position | Ref Base | Serum Base | Serum SNP % | Serum Depth | Serum entropy | | Culture Base | Culture SNP % | Culture Depth | Culture entropy | | Features | S/NS | Residue change | Serum PCR call | P3 PCR call | P5 PCR call |
|---|---|---|---|---|---|---|---|---|---|---|---|---|---|---|---|---|---|
| 9721 | G | | | | | | A | 1.1% | 2232 | 0.0578 | | NS5A | NS | RK | | | |
| 9724 | A | C | 2.6% | 194 | 0.1055 | | | | | | | NS5A | NS | QP | | | |
| 9727 | T | C | 1.0% | 193 | 0.0577 | | C | 1.1% | 2078 | 0.0683 | | NS5A | NS | VA | | | |
| 9764 | G | A | 1.9% | 208 | 0.0950 | | | | | | | NS5A | S | | | | |
| 9789 | G | | | | | | A | 1.1% | 2122 | 0.0586 | | NS5A | NS | DN | | | |
| 9791 | T | | | | | | G | 1.4% | 2134 | 0.0724 | | NS5A | NS | DE | | | |
| 9802 | T | A | 1.1% | 187 | 0.0592 | | | | | | | NS5A | NS | LQ | | | |
| 9814 | T | C | 3.1% | 196 | 0.1369 | | | | | | | NS5A | NS | VA | | | |
| 9837 | A | T | 0.9% | 213 | 0.0532 | | | | | | | NS5A | STOP | K* | | | |
| 9854 | G | A | 1.0% | 192 | 0.0579 | | | | | | | NS5A | S | | | | |
| 9865 | G | A | 2.8% | 251 | 0.1167 | | A | 5.9% | 2091 | 0.2309 | | NS5A | NS | RK | | | |
| 9881 | A | G | 0.8% | 258 | 0.0454 | | | | | | | NS5A | S | | | | |
| 9893 | G | A | 0.7% | 307 | 0.0393 | | | | | | | NS5A | S | | | | |
| 9923 | G | A | 5.1% | 352 | 0.2018 | | A | 2.6% | 2355 | 0.1294 | | NS5A | S | | | | |
| 9951 | A | C | 1.0% | 395 | 0.0566 | | C | 3.6% | 3373 | 0.1576 | | NS5B | NS | TP | | | |
| 9952 | C | T | 1.0% | 404 | 0.0555 | | T | 2.7% | 3442 | 0.1280 | | NS5B | NS | TI | | | |
| 9953 | C | | | | | | A | 1.1% | 3431 | 0.0612 | | NS5B | S | | | | |
| 9957 | C | T | 1.0% | 412 | 0.0547 | | | | | | | NS5B | S | | | | |
| 9962 | T | | | | | | G | 3.2% | 3467 | 0.1586 | | NS5B | NS | FL | | | |
| 9966 | G | | | | | | A | 1.8% | 3494 | 0.0965 | | NS5B | NS | EK | | | |
| 9970 | T | | | | | | G | 1.2% | 3459 | 0.0659 | | NS5B | NS | LR | | | |
| 9986 | G | T | 1.1% | 435 | 0.0705 | | | | | | | NS5B | S | | | | |
| 9987 | C | T | 1.1% | 438 | 0.0701 | | | | | | | NS5B | NS | PS | | | |
| 9988 | C | A | 0.7% | 449 | 0.0401 | | | | | | | NS5B | NS | PH | | | |
| 9998 | G | T | 0.7% | 461 | 0.0393 | | | | | | | NS5B | NS | KN | | | |
| 10001 | C | T | 0.9% | 463 | 0.0497 | | | | | | | NS5B | S | | | | |

| Ref Position | Ref Base | Serum Base | Serum SNP % | Serum Depth | Serum entropy | | Culture Base | Culture SNP % | Culture Depth | Culture entropy | | Features | S/NS | Residue change | Serum PCR call | P3 PCR call | P5 PCR call |
|---|---|---|---|---|---|---|---|---|---|---|---|---|---|---|---|---|---|
| 10015 | T | C | 0.6% | 468 | 0.0388 | | | | | | | NS5B | NS | MT | | | |
| 10052 | G | A | 0.7% | 454 | 0.0398 | | A | 3.1% | 4125 | 0.1439 | | NS5B | S | | | | |
| 10058 | T | C | 0.7% | 451 | 0.0400 | | | | | | | NS5B | S | | | | |
| 10123 | A | G | 0.8% | 359 | 0.0483 | | | | | | | NS5B | NS | YC | | | |
| 10127 | G | T | 1.7% | 229 | 0.0880 | | | | | | | NS5B | S | | | | |
| 10142 | C | T | 1.0% | 205 | 0.0549 | | | | | | | NS5B | S | | | | |
| 10148 | A | | | | | | G | 2.0% | 2507 | 0.0943 | | NS5B | S | | | | |
| 10160 | G | A | 1.5% | 195 | 0.0795 | | | | | | | NS5B | S | | | | |
| 10167 | A | T | 1.1% | 190 | 0.0584 | | | | | | | NS5B | NS | RW | | | |
| 10180 | C | | | | | | T | 1.4% | 2392 | 0.0759 | | NS5B | NS | TI | | | |
| 10182 | G | A | 1.1% | 181 | 0.0608 | | | | | | | NS5B | NS | VI | | | |
| 10210 | T | A | 2.0% | 152 | 0.0970 | | A | 1.4% | 2361 | 0.0803 | | NS5B | NS | LH | | | |
| 10238 | C | T | 6.6% | 152 | 0.2426 | | T | 25.6% | 2400 | 0.5691 | | NS5B | S | | C/t | Y | C/t |
| 10254 | C | A | 15.7% | 140 | 0.4349 | | | | | | | NS5B | NS | LI | C/a | | |
| 10286 | C | T | 2.8% | 145 | 0.1262 | | | | | | | NS5B | S | | | | |
| 10298 | T | C | 4.9% | 143 | 0.1954 | | | | | | | NS5B | S | | T/c | | |
| 10322 | G | | | | | | A | 2.0% | 2293 | 0.0983 | | NS5B | S | | | | |
| 10325 | C | A | 1.3% | 149 | 0.0712 | | | | | | | NS5B | S | | | | |
| 10340 | A | G | 1.1% | 174 | 0.0628 | | | | | | | NS5B | S | | | | |
| 10341 | A | G | 1.1% | 175 | 0.0625 | | | | | | | NS5B | NS | SG | | | |
| 10346 | A | G | 1.6% | 186 | 0.0647 | | | | | | | NS5B | S | | | | |
| 10349 | T | G | 1.6% | 187 | 0.0822 | | | | | | | NS5B | S | | | | |
| 10353 | A | T | 1.0% | 194 | 0.0574 | | | | | | | NS5B | NS | RW | | | |
| 10355 | G | A | 1.5% | 196 | 0.0620 | | | | | | | NS5B | S | | | | |
| 10359 | G | A | 2.4% | 207 | 0.1001 | | | | | | | NS5B | NS | EK | | | |
| 10364 | A | G | 0.9% | 212 | 0.0534 | | | | | | | NS5B | S | | | | |

| Ref Position | Ref Base | Serum Base | Serum SNP % | Serum Depth | Serum entropy | | Culture Base | Culture SNP % | Culture Depth | Culture entropy | | Features | S/NS | Residue change | Serum PCR call | P3 PCR call | P5 PCR call |
|---|---|---|---|---|---|---|---|---|---|---|---|---|---|---|---|---|---|
| 10365 | T | C | 0.9% | 212 | 0.0534 | | | | | | | NS5B | S | | | | |
| 10379 | G | A | 1.4% | 213 | 0.0740 | | | | | | | NS5B | S | | | | |
| 10382 | C | | | | | | T | 12.0% | 2393 | 0.3663 | | NS5B | S | | | | C/t |
| 10399 | G | | | | | | A | 5.9% | 2333 | 0.2246 | | NS5B | NS | SN | | | |
| 10409 | G | A | 0.8% | 250 | 0.0466 | | A | 2.4% | 2130 | 0.1117 | | NS5B | S | | | | |
| 10467 | A | G | 1.1% | 380 | 0.0584 | | | | | | | NS5B | NS | KE | | | |
| 10470 | T | C | 1.3% | 385 | 0.0604 | | | | | | | NS5B | S | | | | |
| 10473 | T | | | | | | C | 1.2% | 3390 | 0.0674 | | NS5B | S | | | | |
| 10497 | C | T | 0.8% | 391 | 0.0450 | | | | | | | NS5B | NS | HY | | | |
| 10508 | G | C | 1.0% | 400 | 0.0629 | | | | | | | NS5B | S | | | | |
| 10553 | G | A | 3.8% | 395 | 0.1615 | | A | 4.4% | 3444 | 0.1860 | | NS5B | S | | | | |
| 10568 | G | A | 1.5% | 399 | 0.0780 | | | | | | | NS5B | S | | | | |
| 10573 | C | T | 1.0% | 384 | 0.0579 | | | | | | | NS5B | NS | AV | | | |
| 10598 | C | | | | | | A | 1.7% | 3458 | 0.0843 | | NS5B | S | | | | |
| 10610 | G | A | 1.4% | 361 | 0.0730 | | | | | | | NS5B | S | | | | |
| 10625 | C | | | | | | T | 1.6% | 3678 | 0.0869 | | NS5B | S | | | | |
| 10634 | A | G | 1.2% | 325 | 0.0664 | | | | | | | NS5B | S | | | | |
| 10638 | T | C | 1.2% | 322 | 0.0559 | | | | | | | NS5B | S | | | | |
| 10641 | G | A | 0.9% | 321 | 0.0409 | | A | 1.9% | 3707 | 0.0934 | | NS5B | NS | VI | | | |
| 10643 | C | | | | | | T | 1.0% | 3720 | 0.0560 | | NS5B | S | | | | |
| 10645 | G | A | 0.9% | 320 | 0.0411 | | | | | | | NS5B | NS | RK | | | |
| 10646 | G | T | 1.3% | 319 | 0.0674 | | | | | | | NS5B | NS | RS | | | |
| 10653 | A | G | 0.6% | 309 | 0.0391 | | | | | | | NS5B | NS | KE | | | |
| 10658 | C | T | 1.7% | 299 | 0.0850 | | | | | | | NS5B | S | | | | |
| 10659 | G | T | 1.4% | 292 | 0.0819 | | | | | | | NS5B | NS | GW | | | |
| 10676 | T | C | 1.0% | 311 | 0.0544 | | | | | | | NS5B | S | | | | |

| Ref Position | Ref Base | Serum Base | Serum SNP % | Serum Depth | Serum entropy | | Culture Base | Culture SNP % | Culture Depth | Culture entropy | | Features | S/NS | Residue change | Serum PCR call | P3 PCR call | P5 PCR call |
|---|---|---|---|---|---|---|---|---|---|---|---|---|---|---|---|---|---|
| 10679 | T | A | 1.0% | 311 | 0.0421 | | | | | | | NS5B | STOP | Y* | | | |
| 10696 | A | C | 0.8% | 265 | 0.0444 | | | | | | | NS5B | NS | KT | | | |
| 10697 | G | | | | | | A | 2.5% | 2071 | 0.1290 | | NS5B | S | | | | |
| 10700 | C | T | 1.5% | 272 | 0.0766 | | T | 3.0% | 2067 | 0.1389 | | NS5B | S | | | | |
| 10703 | G | | | | | | A | 2.0% | 2081 | 0.1104 | | NS5B | S | | | | |
| 10706 | A | G | 0.7% | 291 | 0.0411 | | | | | | | NS5B | S | | | | |
| 10708 | G | | | | | | A | 2.3% | 2114 | 0.1071 | | NS5B | NS | RK | | | |
| 10712 | T | C | 1.0% | 297 | 0.0565 | | | | | | | NS5B | S | | | | |
| 10715 | T | | | | | | C | 1.3% | 2054 | 0.0667 | | NS5B | S | | | | |
| 10721 | C | T | 1.0% | 293 | 0.0571 | | T | 1.0% | 2009 | 0.0563 | | NS5B | S | | | | |
| 10724 | C | T | 1.0% | 294 | 0.0441 | | | | | | | NS5B | S | | | | |
| 10740 | C | T | 0.9% | 343 | 0.0502 | | | | | | | NS5B | S | | | | |
| 10781 | G | T | 0.8% | 489 | 0.0475 | | | | | | | NS5B | NS | ED | | | |
| 10799 | C | | | | | | T | 2.7% | 3104 | 0.1297 | | NS5B | S | | | | C/t |
| 10820 | T | C | 4.1% | 484 | 0.1721 | | C | 2.3% | 3133 | 0.1123 | | NS5B | S | | | | |
| 10839 | G | A | 0.9% | 469 | 0.0491 | | | | | | | NS5B | NS | VI | | | |
| 10853 | A | G | 1.5% | 474 | 0.0700 | | G | 7.0% | 2901 | 0.2575 | | NS5B | S | | | | |
| 10859 | A | | | | | | G | 1.9% | 2822 | 0.0986 | | NS5B | S | | | | |
| 10872 | T | C | 1.0% | 477 | 0.0582 | | | | | | | NS5B | S | | | | |
| 10886 | T | C | 1.5% | 468 | 0.0707 | | | | | | | NS5B | S | | | | |
| 10889 | T | A | 1.3% | 462 | 0.0693 | | | | | | | NS5B | NS | NK | | | |
| 10890 | A | T | 1.3% | 461 | 0.0694 | | | | | | | NS5B | STOP | K* | | | |
| 10910 | T | C | 1.0% | 401 | 0.0559 | | | | | | | NS5B | S | | | | |
| 10918 | A | | | | | | T | 1.1% | 2578 | 0.0645 | | NS5B | NS | NI | | | |
| 10931 | C | T | 0.8% | 381 | 0.0460 | | | | | | | NS5B | S | | | | |
| 10943 | T | C | 0.8% | 375 | 0.0466 | | | | | | | NS5B | S | | | | |

| Ref Position | Ref Base | Serum Base | Serum SNP % | Serum Depth | Serum entropy | | Culture Base | Culture SNP % | Culture Depth | Culture entropy | | Features | S/NS | Residue change | Serum PCR call | P3 PCR call | P5 PCR call |
|---|---|---|---|---|---|---|---|---|---|---|---|---|---|---|---|---|---|
| 10997 | T | | | | | | C | 2.6% | 1898 | 0.1379 | | NS5B | S | | | | T/c |
| 11007 | T | A | 2.2% | 185 | 0.1043 | | | | | | | NS5B | NS | YN | | | |
| 11009 | C | A | 1.8% | 166 | 0.0904 | | | | | | | NS5B | STOP | Y* | | | |
| 11012 | C | A | 2.6% | 151 | 0.1223 | | | | | | | NS5B | STOP | Y* | | | |
| 11015 | T | | | | | | A | 2.6% | 1409 | 0.1412 | | NS5B | STOP | Y* | | | |
| 11030 | C | T | 1.3% | 158 | 0.0679 | | | | | | | NS5B | S | | | | |
| 11045 | C | | | | | | T | 5.4% | 1539 | 0.2099 | | NS5B | S | | | | |
| 11070 | C | T | 1.1% | 180 | 0.0610 | | | | | | | NS5B | NS | PS | | | |
| 11107 | A | T | 1.0% | 207 | 0.0544 | | | | | | | NS5B | NS | NI | | | |
| 11108 | C | T | 44.1% | 213 | 0.6862 | | T | 5.7% | 1497 | 0.2199 | | NS5B | S | | C/t | | C/t |
| 11129 | G | A | 1.7% | 235 | 0.0862 | | | | | | | NS5B | S | | | | |
| 11130 | C | T | 0.8% | 236 | 0.0489 | | | | | | | NS5B | NS | PS | | | |
| 11131 | C | A | 0.8% | 237 | 0.0487 | | | | | | | NS5B | NS | PQ | | | |
| 11162 | T | C | 0.8% | 261 | 0.0450 | | | | | | | NS5B | S | | | | |
| 11165 | C | A | 0.8% | 264 | 0.0445 | | | | | | | NS5B | S | | | | |
| 11168 | A | G | 0.8% | 265 | 0.0444 | | | | | | | NS5B | S | | | | |
| 11187 | T | C | 0.8% | 252 | 0.0463 | | | | | | | NS5B | NS | CR | | | |
| 11189 | C | T | 1.2% | 256 | 0.0638 | | | | | | | NS5B | S | | | | |
| 11202 | A | G | 1.5% | 263 | 0.0788 | | | | | | | NS5B | NS | IV | | | |
| 11204 | C | T | 1.5% | 266 | 0.0780 | | | | | | | NS5B | S | | | | |
| 11207 | A | G | 0.7% | 268 | 0.0440 | | | | | | | NS5B | S | | | | |
| 11210 | C | T | 0.7% | 270 | 0.0437 | | | | | | | NS5B | S | | | | |
| 11219 | C | T | 1.9% | 263 | 0.1070 | | T | 1.2% | 1893 | 0.0693 | | NS5B | S | | | | |
| 11224 | G | T | 0.7% | 267 | 0.0441 | | | | | | | NS5B | NS | RM | | | |
| 11235 | A | T | 0.8% | 262 | 0.0448 | | | | | | | NS5B | NS | IF | | | |
| 11243 | C | T | 0.7% | 270 | 0.0437 | | | | | | | NS5B | S | | | | |

| Ref Position | Ref Base | Serum Base | Serum SNP % | Serum Depth | Serum entropy | | Culture Base | Culture SNP % | Culture Depth | Culture entropy | | Features | S/NS | Residue change | Serum PCR call | P3 PCR call | P5 PCR call |
|---|---|---|---|---|---|---|---|---|---|---|---|---|---|---|---|---|---|
| 11246 | T | C | 0.8% | 266 | 0.0443 | | | | | | | NS5B | S | | | | |
| 11252 | C | T | 0.8% | 250 | 0.0466 | | | | | | | NS5B | S | | | | |
| 11254 | A | T | 0.8% | 248 | 0.0469 | | | | | | | NS5B | NS | DV | | | |
| 11259 | T | C | 0.8% | 256 | 0.0457 | | | | | | | NS5B | NS | FL | | | |
| 11262 | T | C | 0.8% | 254 | 0.0460 | | | | | | | NS5B | S | | | | |
| 11266 | T | C | 0.8% | 254 | 0.0460 | | | | | | | NS5B | NS | IT | | | |
| 11273 | G | | | | | | A | 1.4% | 2161 | 0.0806 | | NS5B | S | | | | |
| 11295 | G | T | 0.8% | 246 | 0.0472 | | | | | | | NS5B | NS | AS | | | |
| 11306 | G | A | 0.7% | 273 | 0.0433 | | | | | | | NS5B | S | | | | |
| 11309 | G | A | 0.7% | 272 | 0.0434 | | | | | | | NS5B | NS | MI | | | |
| 11314 | T | G | 0.7% | 272 | 0.0434 | | | | | | | NS5B | NS | IS | | | |
| 11328 | G | T | 0.8% | 261 | 0.0450 | | | | | | | NS5B | NS | GC | | | |
| 11339 | A | G | 0.7% | 267 | 0.0441 | | | | | | | NS5B | S | | | | |
| 11348 | G | | | | | | A | 3.0% | 2890 | 0.1351 | | NS5B | S | | | | |
| 11351 | A | | | | | | G | 1.3% | 2855 | 0.0715 | | NS5B | S | | | | |
| 11355 | G | | | | | | A | 1.6% | 2847 | 0.0827 | | NS5B | NS | EK | | | |
| 11357 | A | G | 7.8% | 230 | 0.2745 | | G | 28.3% | 2830 | 0.5958 | | NS5B | S | | A/g | A/g | A/g |
| 11360 | A | G | 0.9% | 229 | 0.0501 | | | | | | | NS5B | S | | | | |
| 11362 | T | | | | | | A | 1.2% | 2777 | 0.0723 | | NS5B | NS | MK | | | |
| 11369 | T | | | | | | A | 4.4% | 2813 | 0.1854 | | NS5B | S | | | | |
| 11372 | C | T | 0.9% | 231 | 0.0497 | | | | | | | NS5B | S | | | | |
| 11393 | G | T | 1.0% | 207 | 0.0544 | | | | | | | NS5B | NS | ED | | | |
| 11444 | T | A | 0.8% | 243 | 0.0477 | | | | | | | NS5B | NS | SR | | | |
| 11451 | G | T | 0.7% | 273 | 0.0433 | | | | | | | NS5B | NS | AS | | | |
| 11515 | G | A | 1.1% | 358 | 0.0613 | | | | | | | NS5B | NS | RK | | | |
| 11519 | T | C | 1.0% | 383 | 0.0484 | | | | | | | NS5B | NS | SW | | | |

| Ref Position | Ref Base | Serum Base | Serum SNP % | Serum Depth | Serum entropy | | Culture Base | Culture SNP % | Culture Depth | Culture entropy | | Features | S/NS | Residue change | Serum PCR call | P3 PCR call | P5 PCR call |
|---|---|---|---|---|---|---|---|---|---|---|---|---|---|---|---|---|---|
| 11531 | C | | | | | | T | 4.8% | 2812 | 0.1929 | | NS5B | S | | | | |
| 11571 | T | | | | | | A | 3.5% | 4138 | 0.1541 | | NS5B | NS | WR | | | |
| 11579 | G | A | 1.0% | 573 | 0.0648 | | | | | | | NS5B | S | | | | |
| 11584 | T | C | 1.4% | 572 | 0.0736 | | | | | | | NS5B | NS | VA | | | |
| 11606 | C | T | 1.6% | 569 | 0.0813 | | | | | | | NS5B | S | | | | |
| 11629 | C | T | 0.9% | 568 | 0.0504 | | | | | | | NS5B | NS | AV | | | |
| 11630 | G | A | 0.9% | 567 | 0.0505 | | | | | | | NS5B | S | | | | |
| 11636 | A | G | 0.7% | 558 | 0.0425 | | G | 6.1% | 3970 | 0.2291 | | NS5B | S | | | | |
| 11639 | A | G | 0.7% | 556 | 0.0427 | | | | | | | NS5B | S | | | | |
| 11681 | T | C | 1.0% | 521 | 0.0541 | | C | 1.6% | 3863 | 0.0798 | | NS5B | S | | | | |
| 11702 | T | C | 1.1% | 528 | 0.0554 | | | | | | | NS5B | S | | | | |
| 11765 | G | A | 0.9% | 429 | 0.0529 | | | | | | | NS5B | S | | | | |
| 11786 | C | | | | | | T | 4.5% | 2482 | 0.1900 | | NS5B | S | | | | |
| 11797 | G | A | 0.8% | 247 | 0.0471 | | | | | | | NS5B | NS | RK | | | |
| 11798 | A | G | 1.6% | 249 | 0.0823 | | | | | | | NS5B | S | | | | |
| 11800 | T | A | 0.8% | 252 | 0.0463 | | A | 1.6% | 2288 | 0.0924 | | NS5B | NS | IK | | | |
| 11807 | G | T | 2.7% | 261 | 0.1396 | | | | | | | NS5B | NS | QH | | | |
| 11820 | G | A | 1.2% | 256 | 0.0638 | | | | | | | NS5B | NS | VI | | | |
| 11828 | G | A | 1.7% | 241 | 0.0845 | | | | | | | NS5B | S | | | | |
| 11832 | G | | | | | | A | 1.1% | 2297 | 0.0600 | | NS5B | NS | EK | | | |
| 11844 | C | | | | | | A | 4.9% | 2518 | 0.1955 | | NS5B | NS | LI | | | |
| 11884 | A | T | 0.7% | 279 | 0.0425 | | | | | | | NS5B | NS | HL | | | |
| 11889 | T | A | 0.8% | 266 | 0.0443 | | | | | | | NS5B | NS | YN | | | |
| 11897 | T | C | 8.5% | 272 | 0.2898 | | C | 27.2% | 2539 | 0.5965 | | NS5B | S | | | T/c | |
| 11900 | C | T | 1.1% | 285 | 0.0584 | | | | | | | NS5B | S | | | | |
| 11903 | A | G | 1.0% | 287 | 0.0581 | | | | | | | NS5B | S | | | | |

| Ref Position | Ref Base | Serum Base | Serum SNP % | Serum Depth | Serum entropy | | Culture Base | Culture SNP % | Culture Depth | Culture entropy | | Features | S/NS | Residue change | Serum PCR call | P3 PCR call | P5 PCR call |
|---|---|---|---|---|---|---|---|---|---|---|---|---|---|---|---|---|---|
| 11906 | T | | | | | | C | 1.5% | 2637 | 0.0786 | | NS5B | S | | | | |
| 11930 | T | | | | | | C | 11.0% | 2349 | 0.3457 | | NS5B | S | | | T/c | T/c |
| 11936 | A | G | 0.6% | 312 | 0.0388 | | G | 1.5% | 2325 | 0.0781 | | NS5B | S | | | | |
| 11937 | C | | | | | | T | 2.1% | 2328 | 0.1037 | | NS5B | S | | | | |
| 11939 | | | | | | | | | | | | NS5B | S | | | | A/g |
| 11957 | G | A | 0.8% | 396 | 0.0445 | | | | | | | NS5B | S | | | | |
| 11993 | G | A | 1.1% | 446 | 0.0534 | | | | | | | NS5B | S | | | | |
| 11996 | A | | | | | | G | 1.9% | 2048 | 0.0909 | | NS5B | S | | | | A/g |
| 11999 | T | C | 0.9% | 446 | 0.0512 | | | | | | | NS5B | S | | | | |
| 12010 | A | | | | | | C | 2.5% | 1943 | 0.1144 | | NS5B | NS | NT | | | |
| 12012 | A | G | 1.9% | 423 | 0.0866 | | | | | | | NS5B | NS | IV | | | |
| 12029 | A | | | | | | G | 1.8% | 1742 | 0.0959 | | NS5B | S | | | | |
| 12033 | G | A | 1.2% | 431 | 0.0550 | | | | | | | NS5B | NS | VI | | | |
| 12035 | C | T | 0.7% | 441 | 0.0407 | | | | | | | NS5B | S | | | | |
| 12079 | T | | | | | | C | 1.7% | 3770 | 0.0867 | | UTR | NA | | | | T/c |
| 12084 | T | C | 5.5% | 656 | 0.2097 | | | | | | | UTR | NA | | | | |
| 12093 | G | A | 2.3% | 652 | 0.1095 | | A | 1.7% | 3967 | 0.0839 | | UTR | NA | | | | |
| 12095 | T | C | 4.0% | 648 | 0.1683 | | C | 5.0% | 3965 | 0.2021 | | UTR | NA | | | | |
| 12107 | C | T | 2.6% | 618 | 0.1202 | | | | | | | UTR | NA | | | | |
| 12110 | A | G | 1.0% | 614 | 0.0612 | | | | | | | UTR | NA | | | | |
| 12112 | T | C | 1.8% | 611 | 0.0933 | | | | | | | UTR | NA | | | | |
| 12114 | T | C | 1.1% | 610 | 0.0627 | | C | 3.7% | 3789 | 0.1657 | | UTR | NA | | | | |
| 12116 | T | | | | | | C | 2.8% | 3788 | 0.1328 | | UTR | NA | | | | |
| 12133 | T | C | 1.2% | 590 | 0.0649 | | | | | | | UTR | NA | | | | |
| 12176 | C | T | 3.4% | 532 | 0.1478 | | T | 3.4% | 3425 | 0.1522 | | UTR | NA | | | | |
| 12178 | G | A | 0.8% | 530 | 0.0444 | | | | | | | UTR | NA | | | | |

| Ref Position | Ref Base | Serum Base | Serum SNP % | Serum Depth | Serum entropy | | Culture Base | Culture SNP % | Culture Depth | Culture entropy | | Features | S/NS | Residue change | Serum PCR call | P3 PCR call | P5 PCR call |
|---|---|---|---|---|---|---|---|---|---|---|---|---|---|---|---|---|---|
| 12193 | T | A | 0.6% | 466 | 0.0389 | | | | | | | UTR | NA | | | | |
| 12199 | A | | | | | | G | 8.7% | 3288 | 0.2958 | | UTR | NA | | | | |
| 12211 | C | T | 0.8% | 364 | 0.0478 | | | | | | | UTR | NA | | | | |
| 12223 | A | T | 0.9% | 346 | 0.0384 | | | | | | | UTR | NA | | | | |
| 12228 | A | G | 0.9% | 340 | 0.0505 | | | | | | | UTR | NA | | | | |
| 12238 | T | G | 0.9% | 325 | 0.0405 | | | | | | | UTR | NA | | | | |
| 12249 | A | T | 0.6% | 313 | 0.0387 | | | | | | | UTR | NA | | | | |
| 12279 | C | | | | | | T | 2.3% | 1043 | 0.1321 | | UTR | NA | | | | |
| 12280 | C | | | | | | A | 1.3% | 862 | 0.0683 | | UTR | NA | | | | |

**NADL variant base sites**

| NADL position | MRI103 position | NADL Base | Variant Base | NADL SNP % | NADL Depth | NADL entropy | Features |
|---|---|---|---|---|---|---|---|
| 25 | 10 | T | A | 2.1% | 1056 | 0.1181 | UTR |
| 34 | 19 | T | C | 2.4% | 1602 | 0.1207 | UTR |
| 1830 | 1816 | G | A | 6.3% | 9383 | 0.2348 | E1 |
| 2564 | 2549 | C | A | 2.9% | 8143 | 0.1349 | E2 |
| 4022 | 4007 | A | T | 3.8% | 4759 | 0.1643 | NS2-3 |
| 4068 | 4053 | T | G | 3.4% | 4673 | 0.1529 | NS2-3 |
| 5872 | 5588 | A | G | 3.7% | 5148 | 0.1614 | NS2-3 |
| 10838 | 10554 | A | G | 4.5% | 15956 | 0.1932 | NS5B |
| 10869 | 10585 | A | G | 2.2% | 13461 | 0.1138 | UTR |
| 10890 | 10606 | T | G | 3.9% | 12493 | 0.1777 | UTR |
| 12293 | 12009 | A | G | 2.1% | 12725 | 0.1018 | UTR |
| 12367 | 12082 | A | G | 2.4% | 19659 | 0.1152 | UTR |
